# Relaxed evolutionary constraints on mitochondrial electron-transfer-associated variants in human cancers

**DOI:** 10.64898/2026.10.21.760641

**Authors:** Ji-Yong Sung, Jae-Ho Cheong

**Affiliations:** Department of Neurosurgery, Seoul National University Bundang Hospital, Seoul National University College of Medicine, Seongnam-si, Republic of Korea; Department of Surgery, Yonsei University College of Medicine, Seoul 03722, Republic of Korea

## Abstract

Mitochondrial electron transfer depends on quantum-mechanical coupling between redox cofactors, but its evolutionary constraints in cancer remain unclear. We combined the human complex III cryo-EM structure, Pathways calculations and Marcus theory to predict how missense substitutions in mitochondrial cytochrome b (MT-CYB), which coordinates the Q-cycle haem centres, affect electron-transfer kinetics. Comparing 524 somatic mutations from TCGA and PCAWG with 597 germline variant sites, we found that charge-changing substitutions near the haems were depleted in the germline (observed/expected, 0.12) but enriched in tumours (1.81). Greater predicted kinetic perturbations corresponded to stronger germline depletion and tumour enrichment, predominantly at low heteroplasmy and independently of tumour purity. Predicted kinetic effects correlated strongly with electrostatic perturbations (r = 0.94), precluding identification of tunnelling-specific selection. These findings support relaxed purifying selection on MT-CYB variants predicted to disrupt mitochondrial electron transfer during the evolution of diverse human cancers, without demonstrating adaptive benefits or selection on tunnelling itself.

## Introduction

Electron flow through the mitochondrial respiratory chain involves successive electron-transfer reactions between redox cofactors separated by subnanometre to nanometre-scale distances within and between protein subunits ^1,2^ At the level of individual steps, electron transfer can proceed through quantum-mechanical tunnelling, mediated by electronic coupling between spatially separated donor and acceptor states. The magnitude of this coupling generally decreases approximately exponentially with distance and is influenced by the covalent and non-covalent structure of the intervening protein environment. ^1,2^ Within the framework of non-adiabatic Marcus theory, electron-transfer rates are determined by the interplay of donor–acceptor electronic coupling, the thermodynamic driving force and nuclear reorganization energy ^3^ Decades of experimental and theoretical studies have established that biological redox centres are spatially organized to support electron transfer at rates compatible with their physiological functions, suggesting that the structural and energetic determinants of electron transfer are subject to evolutionary constraints. ^4^ Whether the physical determinants of mitochondrial electron transfer are subject to altered evolutionary constraints during tumour development remains poorly understood. Somatic mutations in mitochondrial DNA (mtDNA) are common across human cancers, and large-scale genomic studies have revealed distinct mutational patterns among respiratory complexes and tumour types. In particular, previous analyses have identified complex- and tissue-dependent differences in the distribution of missense and truncating mutations, suggesting that mitochondrial respiratory functions may experience different selective constraints in cancer. ^5,6^ Meanwhile, population-scale studies have demonstrated strong purifying selection across the human mitochondrial genome, with functionally constrained residues showing pronounced three-dimensional clustering within respiratory proteins. ^7^ However, these investigations have primarily characterized mutational selection in terms of gene identity, respiratory complex, predicted functional consequence or evolutionary conservation. Whether cancer-associated differences in mutational constraint can be quantitatively linked to the structural and energetic determinants of mitochondrial electron-transfer kinetics remains largely unexplored. Previous pan-cancer studies have revealed respiratory-complex-dependent patterns of mtDNA mutations, including an enrichment of truncating alterations in subsets of complex I genes and a contrasting predominance of missense variation in complex III.

Yet these studies classify variants by gene, complex or evolutionary conservation; none has asked whether the physical process of electron tunnelling, as predicted from structure, is under differential selection in the germline and in tumours. These distinct mutational patterns raise the possibility that different respiratory complexes are subject to different functional constraints during tumour evolution, motivating a mechanistic investigation of how MT-CYB missense variants affect the physical determinants of electron transfer. Here, we address this question by focusing on cytochrome b (MT-CYB), the only mtDNA-encoded subunit of complex III and a central component of its proton-motive Q cycle. ^8^

Complexes I and II provide major routes of electron entry into the ubiquinone pool, whereas complex III oxidizes ubiquinol supplied by these and other pathways and couples electron transfer to cytochrome c with proton translocation through the bifurcated Q cycle. MT-CYB forms the structural framework of the Qo and Qi quinone-binding sites and coordinates both haem bL and haem bH, which mediate electron transfer along the low-potential branch of the Q cycle. Consequently, missense substitutions in MT-CYB can potentially alter the electrostatic environment and redox potentials of these cofactors, thereby modulating electron-transfer kinetics. In contrast, the principal FMN–iron–sulfur electron-transfer chain of complex I resides predominantly within nuclear-encoded peripheral-arm subunits, while its mtDNA-encoded ND subunits contribute primarily to quinone-site and membrane-arm functions. These structural features make MT-CYB a particularly suitable model for quantitatively linking mitochondrial genetic variation to the biophysical determinants of electron transfer.

Using the cryo-EM structure of human complex III, ^9^ we compute Pathways electronic couplings for the electron-transfer steps of cytochrome b and combine them with Marcus theory to predict the effects of all possible missense substitutions on electron-transfer rates. We then evaluate these predictions against 524 somatic MT-CYB missense mutations from TCGA and PCAWG and 597 germline variant sites from HelixMTdb, examining their relationships with tumour heteroplasmy, tumour purity, and nuclear mutational signatures, ^10^ as well as an analogous electron-transfer step in complex IV.

## Results

### A structural and electron-tunnelling map of cytochrome b

We built the analysis on the cryo-EM structure of human kidney complex III (PDB 9CG3, 2.96 Å). Its cytochrome b sequence matches the rCRS translation of MT-CYB at all 378 modelled positions (residues 2–379; residues 1 and 380 are not resolved in the map), so every modelled residue maps to an mtDNA codon (**Fig. 1e**). No quinone or inhibitor is modelled in 9CG3, so we located the quinone pockets geometrically. The Qi pocket (∼227 Å³) is lined by His201, Ser205, Phe220, Tyr224 and Asp228, and the Qo pocket (∼60 Å³) by Tyr131 and Glu271 of the PEWY motif, matching the known sites (**Fig. 1b**). Pathways electronic couplings reproduced the expected hierarchy of the low-potential chain (**Fig. 1c**). Coupling between the Qo pocket and haem bL (edge distance 6.4 Å; log □□|T_DA| = −2.7) and between haem bH and the Qi pocket (3.8 Å; −2.0) far exceeds the transmembrane haem bL→bH step (9.3 Å; −5.7). In this simplified Pathways analysis, single side-chain truncations changed the bL→bH coupling little, consistent with compensating alternative paths. We therefore modelled missense effects primarily through the driving-force term of Marcus theory, using estimated shifts in haem redox potentials; this modelling choice does not rule out mutation-induced changes in geometry, reorganization energy or proton-coupled Q-cycle chemistry (**Fig. 1a,d**). To capture this, we estimated for each substitution the screened Coulomb shift that its net charge change imposes on each redox centre. We converted these shifts into rate changes for three Q-cycle steps (Qo semiquinone→bL, bL→bH, bH→Qi). For example, G48E is predicted to accelerate bL→bH about 30-fold and G116D to slow it about 130-fold (**Fig. 1d**). We grouped residues into five structural classes (Qo site, ISP interface, Qi site, haem b environment, other) for the selection analyses that follow (**Fig. 1e**).

**Figure 1.**
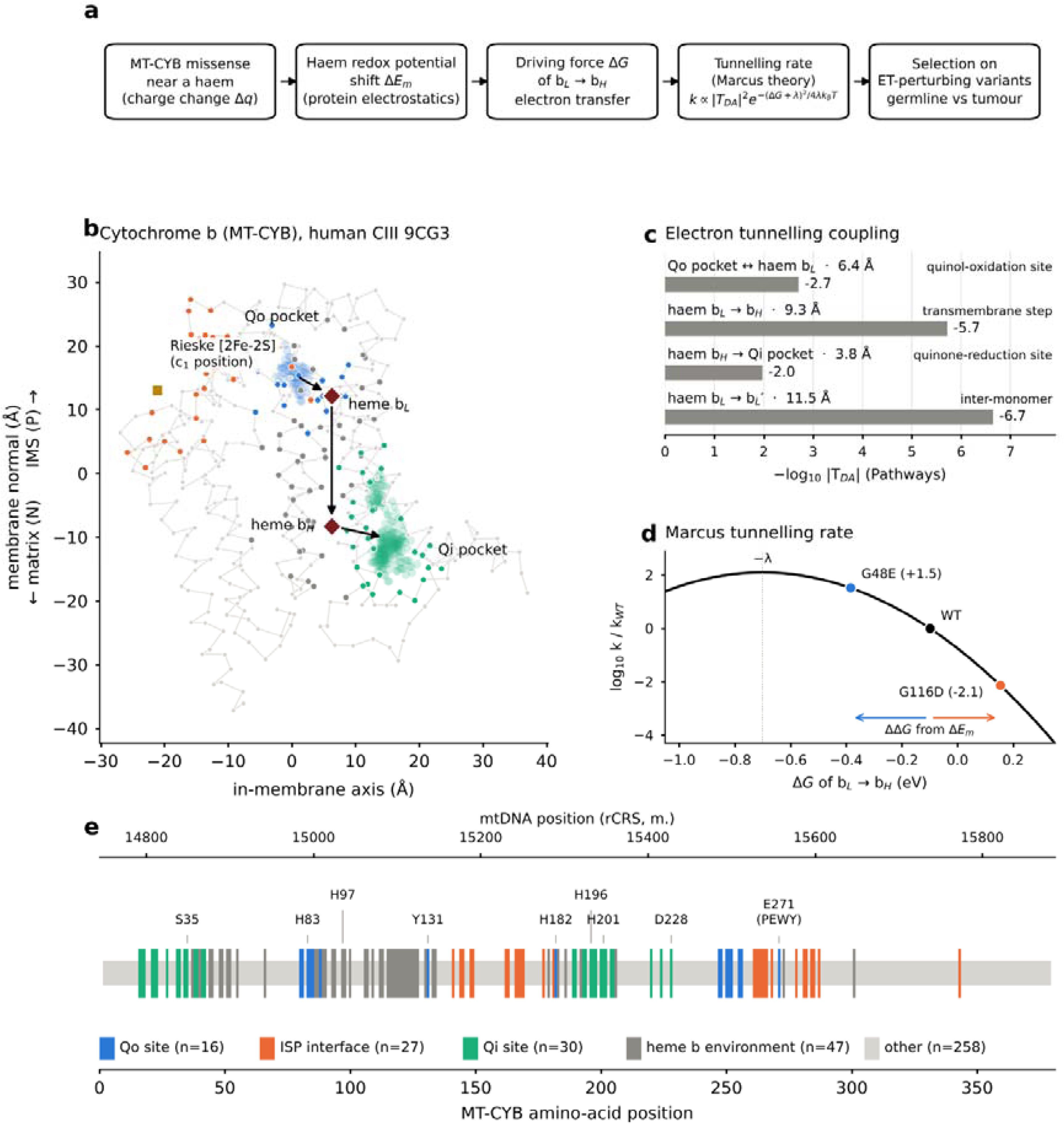
Structural and quantum electron-tunnelling framework for cytochrome b variants. a, Conceptual model. A missense variant that changes the net charge of a residue near a haem shifts the haem redox midpoint potential (ΔEm) through protein electrostatics. This alters the driving force (ΔG) of the transmembrane haem bL→bH electron transfer (ET) and therefore its non-adiabatic tunnelling rate (k), given by Marcus theory together with the donor– acceptor electronic coupling |TDA|. Selection on such ET-perturbing variants is compared between the germline and tumours in Figs 2 and 3. b, Cα trace of cytochrome b (chain J) from the cryo-EM structure of human kidney respiratory complex III (PDB 9CG3, 2.96 Å), projected onto the membrane normal (haem bH→bL axis; IMS, intermembrane space, P side, at top) and the principal in-membrane axis. Residues are coloured by functional class as in e. Translucent clouds show the Qo and Qi quinone pockets, detected geometrically in the ligand-free structure. Diamonds, haem iron atoms; square, Rieske [2Fe–2S] cluster (c position in this structure). Arrows show the low-potential electron path from the Qo site through haems bL and bH to the Qi site. c, Electronic coupling (−log |TDA|) for the ET steps of cytochrome b, calculated with the Pathways model. Numbers next to each step give the edge-to-edge donor–acceptor distance. Shorter bars indicate stronger coupling. d, Marcus dependence of the bL→bH tunnelling rate on driving force, relative to wild type (WT; ΔG0 = −0.10 eV, λ = 0.70 eV, T = 310 K). Coloured points show two tumour-observed variants positioned by their electrostatically estimated driving-force change (ΔΔG; effective dielectric constant ε = 8): G48E (ΔΔG = −0.29 eV, 30-fold acceleration) and G116D (ΔΔG = +0.25 eV, ∼130-fold deceleration). Dotted line, activationless optimum (ΔG = −λ). e, Functional classification of the 378 modelled MT-CYB residues (2–379), aligned with mtDNA coordinates on the revised Cambridge Reference Sequence (rCRS; top axis). Taller bars mark classified residues. Labels indicate haem bL ligands (H83, H182), haem bH ligands (H97, H196), Qi residues (H201, D228), Qo residues (Y131, E271 of the PEWY motif) and the disease-associated residue S35. Numbers of residues per class are given in the key. Pathways couplings used decay factors of 0.6 per covalent bond with distance-dependent hydrogen-bond and through-space terms. Parameter values and sensitivity analyses (ε = 4, 8 and 16; λ scaled by 0.7×, 1× and 1.4× the reference value of each step, i.e. 0.49–0.98 eV for bL→bH) are given in Methods. Source data are provided as a Source Data file.

### Charge-changing substitutions near the haems are purged from the germline but enriched in tumours

We analysed somatic mtDNA calls from 10,132 TCGA exomes and 2,536 PCAWG genomes, and germline variant sites from HelixMTdb. Somatic calls at sites carried by more than ten HelixMTdb individuals were removed as likely germline polymorphisms (for example m.15497G>A, G251S, homoplasmic in 690 individuals). Because this filter preferentially removes tolerated sites, expected counts for somatic mutations were computed over the same restricted set of sites. Without this restriction, enrichment estimates were inflated by up to 50% (Methods). After filtering, 524 somatic MT-CYB missense mutations remained (**Fig. 2a**). Expected counts came from each cohort’s own strand-aware trinucleotide spectrum. The model was well calibrated on synonymous variants in every class and cohort (all P ≥ 0.17; Supplementary Fig. 2).

**Figure 2.**
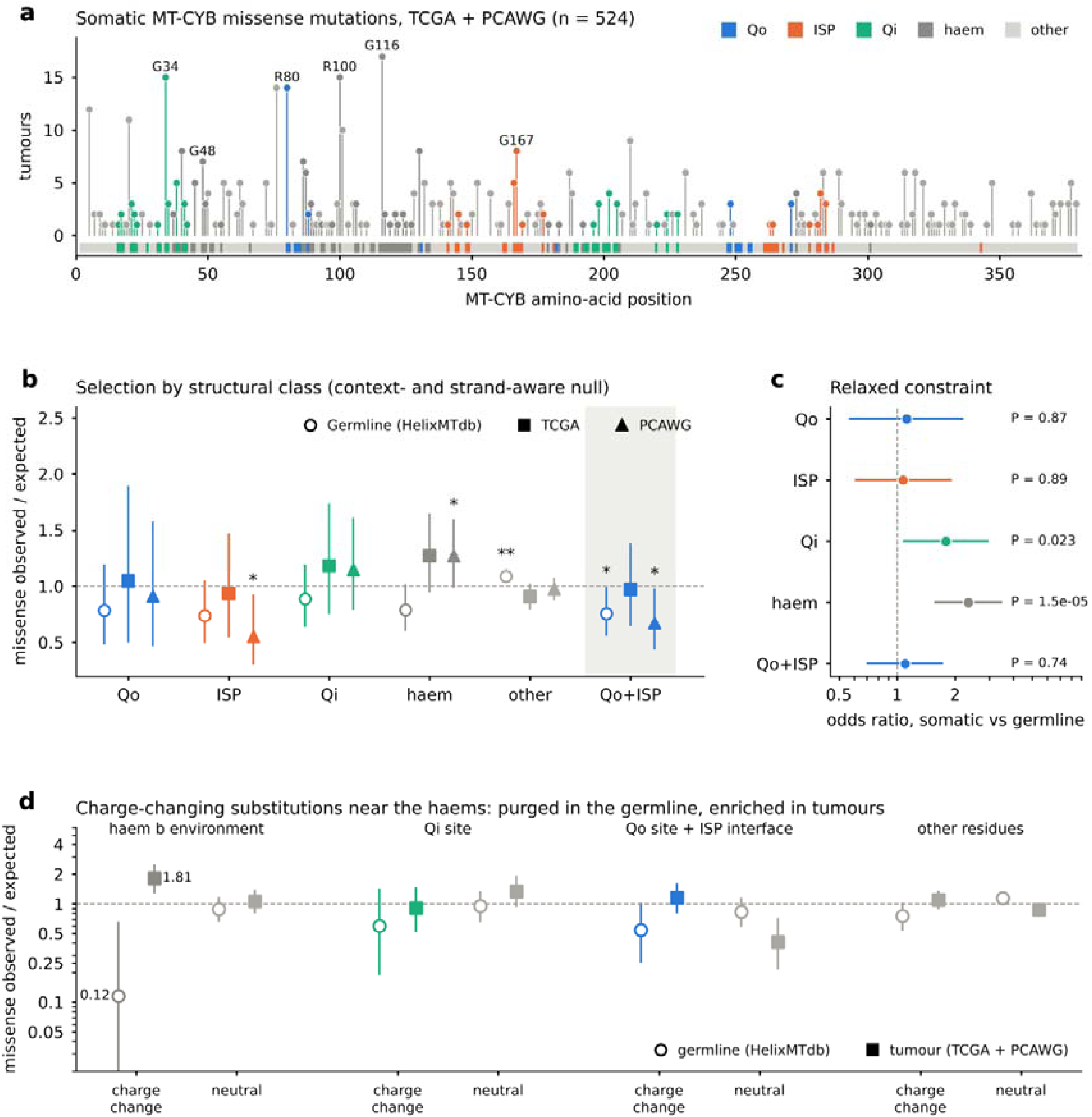
Charge-changing substitutions near the cytochrome b haems are purged from the germline but enriched in tumours. a, Somatic MT-CYB missense mutations in 2,536 PCAWG whole genomes and 10,132 TCGA exomes (n = 524 mutations), coloured by functional class (Fig. 1e). Mutations at sites carrying the same allele in more than ten HelixMTdb individuals were excluded as likely germline polymorphisms. Recurrent positions are labelled. b, Observed/expected ratio of missense mutations in each structural class for germline variant sites (HelixMTdb; open circles) and somatic mutations (TCGA, squares; PCAWG, triangles). The shaded group pools the Qo site and ISP interface, which share the bL leak path and are shown separately to its left. Expected counts were derived from each cohort’s own strand-aware trinucleotide mutation spectrum, estimated from all mtDNA single-nucleotide variants excluding missense variants in the analysed genes. Error bars, 95% binomial confidence intervals; *, P < 0.05; **, P < 0.01 (two-sided exact binomial test). c, Odds ratio for the share of missense variants falling in each class, comparing somatic with germline variants (two-sided Fisher’s exact test). Error bars, 95% confidence intervals. d, Observed/expected ratio of missense variants split by structural class and by whether the substitution changes residue net charge (|Δq| ≥ 0.9), for germline sites (open circles) and pooled somatic mutations (squares). Charge-changing substitutions in the haem b environment were depleted in the germline (1 observed versus 8.6 expected; O/E = 0.12, P = 0.003) and enriched in tumours (O/E = 1.81, P = 4.7 × 10□□), whereas charge-neutral substitutions at the same residues were close to expectation. Qo-site and ISP-interface residues were pooled for this analysis owing to low counts. Error bars, 95% binomial confidence intervals. Source data are provided as a Source Data file.

In the germline, missense variants were depleted from all structured classes (observed/expected 0.73–0.89) and in excess among other residues (1.09; **Fig. 2b**). In tumours, the haem b environment showed the opposite pattern (1.27 in TCGA, 1.27 in PCAWG). Its share of missense variants was 2.3-fold higher in tumours than in the germline (P = 1.5 × 10□□; **Fig. 2c**). The Qi site showed a weaker shift (odds ratio 1.79, P = 0.023). Qo and ISP-interface variants were not enriched; the ISP interface was depleted in PCAWG (0.55, P = 0.017). The haem b signal was carried entirely by substitutions that change residue charge (**Fig. 2d**). Only one charge-changing haem-environment variant was observed in the germline against 8.6 expected (0.12, P = 0.003), whereas tumours carried 1.81-fold more than expected (P = 4.7 × 10□). Charge-neutral substitutions at the same residues were close to expectation in both (0.89 and 1.06). Results were similar without the polymorphism filter (**Supplementary Fig. 1**). The 789 patients shared between TCGA and PCAWG did not drive these pooled estimates: repeating the analysis after removing the 628 overlapping PCAWG samples that carried mtDNA calls left the haem-environment charge-change enrichment unchanged (O/E = 1.84, P = 1.3 × 10 ³) and the tunnelling-perturbation enrichment essentially so (O/E = 1.25, P = 0.016).

### Variants predicted to perturb electron tunnelling show opposite selection in germline and tumours

We next scored every possible MT-CYB missense substitution by its predicted effect on the three Q-cycle tunnelling rates, combining Marcus driving-force changes with Pathways couplings. Variants predicted to change at least one rate by two-fold or more were depleted in the germline (observed/expected 0.53, P = 3 × 10□□) and enriched in tumours (1.31, P = 0.001; **Fig. 3a**). The tumour signal was significant in PCAWG (1.36, P = 0.004) and directionally consistent in TCGA (1.23, P = 0.11). The effect was graded (**Fig. 3b**). As the predicted rate change rose from <0.1 to ≥1 log□□ units, germline observed/expected fell from 1.08 to 0.47 and tumour observed/expected rose from 0.90 to 1.67 (P < 0.001 for the top bin). The pattern held across effective dielectric constants of 4–16 and reorganization energies of 0.7–1.4 times the reference values (germline 0.53–0.59; tumours 1.24–1.42; **Fig. 3c**). The predicted tunnelling perturbation is, however, almost collinear with the electrostatic shift at the nearest redox centre (r = 0.94 across all possible substitutions; **Fig. 3d**). In quasi-Poisson models adjusting for that shift, charge change, hydropathy, side-chain volume, burial and distance, the tunnelling-specific component was not significant in any cohort (P = 0.34, 0.87 and 0.07; **Fig. 3e**). Population data therefore show that tumours tolerate variants that would perturb tunnelling, but cannot attribute this to tunnelling rather than to electrostatic perturbation of the cofactors in general. Several tumour-observed variants lie far from the collinear trend and make opposite predictions. G116D, G48E, G34D and R100Q are predicted to perturb tunnelling more than their electrostatic shift implies, whereas R80H, E202K and G231D are predicted to do the reverse. Measuring haem potentials and bL→bH rates for these variants would separate the two explanations. The analogous analysis of the CuA→haem a step of complex IV (MT-CO1, MT-CO2) reproduced the germline depletion but showed no tumour enrichment (**Supplementary Fig. 3**). In TCGA, tumours carrying high-perturbation MT-CYB missense mutations were not enriched for truncating mutations in complex I (ND genes) (14.9% versus 17.3%; Fisher P = 0.74). Tumours carrying high-perturbation MT-CYB missense mutations were not consistently enriched for truncating mutations in complex I (ND genes). In TCGA the association was absent (cancer-type-stratified Cochran–Mantel–Haenszel odds ratio 0.56, P = 0.20); in PCAWG a nominal positive association (1.67, P = 0.023) did not survive removal of the samples shared with TCGA (1.52, P = 0.10), and the pooled de-duplicated estimate was null (1.17, P = 0.48). This co-occurrence test does not constitute a matched analysis of electron-transfer perturbation or selection across complexes I and III.

**Figure 3.**
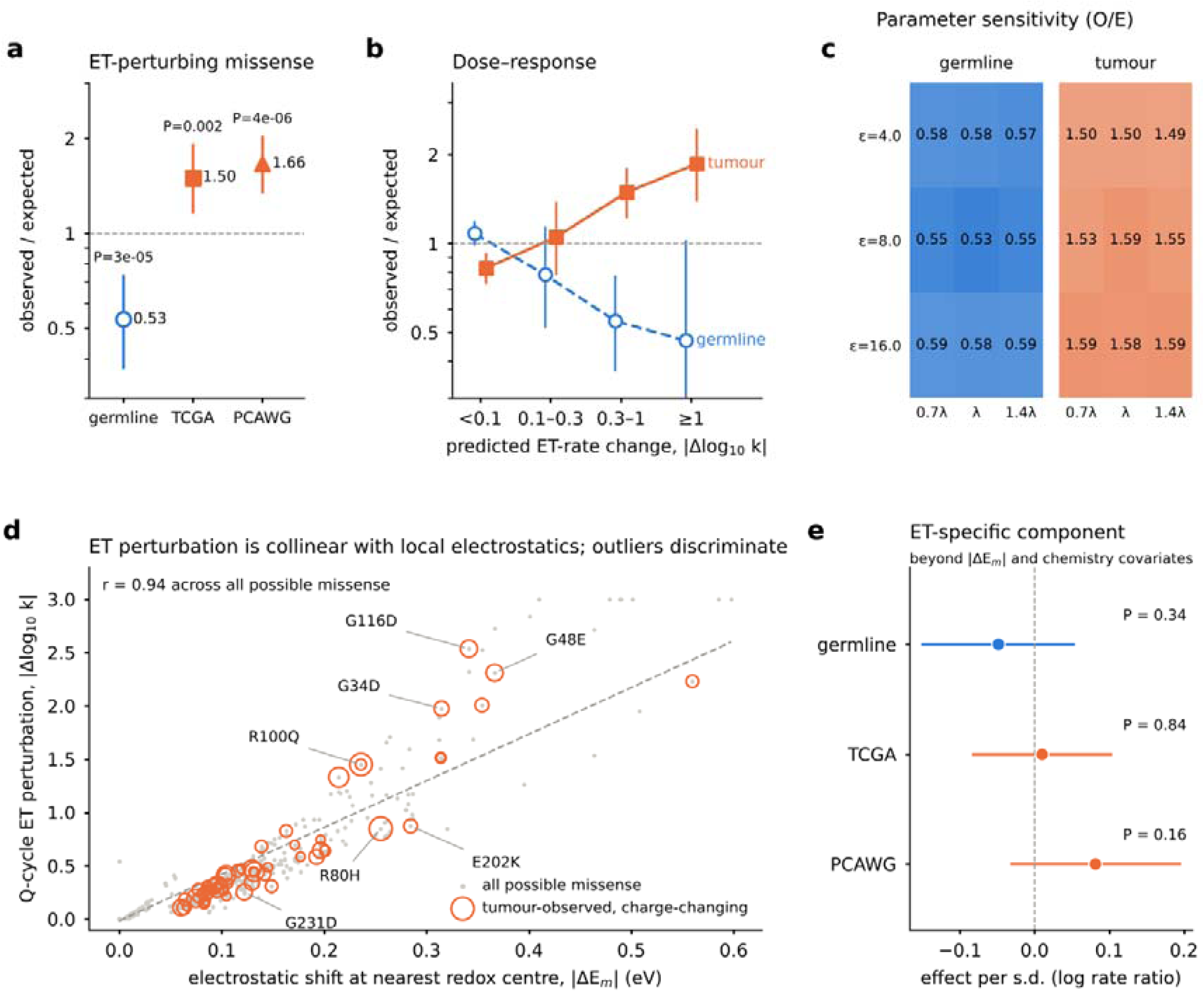
Tumours accumulate, and the germline purges, cytochrome b variants predicted to perturb quantum electron tunnelling. a, Observed/expected ratio of high-perturbation missense variants, defined as those predicted to change the rate of at least one cytochrome b electron-transfer (ET) step by ≥2-fold (|Δlog□□ k| ≥ 0.3), in germline variant sites (HelixMTdb) and somatic mutations (TCGA, PCAWG). Rates were calculated with a three-step Q-cycle model (Qo semiquinone → haem bL → haem bH → Qi quinone) combining Marcus non-adiabatic ET theory with Pathways electronic couplings (Fig. 1c,d). Driving-force changes were estimated from the screened Coulomb shift of each redox-centre potential caused by the substitution’s net charge change. Expected counts were derived from each cohort’s own strand-aware trinucleotide mutation spectrum across all possible MT-CYB missense substitutions. Error bars, 95% Poisson confidence intervals; two-sided Poisson tests. b, Observed/expected ratio across bins of predicted ET-rate change for germline sites (open circles, dashed line) and pooled somatic mutations (squares, solid line). c, Observed/expected ratio of high-perturbation variants across effective dielectric constants (ε = 4, 8, 16) and reorganization energies (0.7×, 1×, 1.4× the reference value of each step). d, Predicted Q-cycle ET perturbation versus the electrostatic potential shift at the nearest redox centre for all possible MT-CYB missense substitutions (grey) and for charge-changing substitutions observed in tumours (orange circles; area proportional to the number of tumours). Dashed line, least-squares fit. Labelled variants lie furthest above (G116D, G48E, G34D, R100Q) or below (R80H, E202K, G231D) the fit and are predicted to discriminate ET-specific from purely electrostatic effects experimentally. e, Effect of the ET-specific component of the perturbation score (the score orthogonalized to the electrostatic shift and the charge change) in quasi-Poisson regression with spectrum offsets, adjusting for electrostatic shift, charge change, hydropathy change, side-chain volume change, burial and distance to the nearest redox centre. Error bars, 95% confidence intervals. Source data are provided as a Source Data file.

### Tunnelling-perturbing variants are tolerated only below a heteroplasmy threshold

If tumours positively selected tunnelling-perturbing variants, these should expand towards homoplasmy. Instead, their enrichment was confined to variants carried by a minority of mtDNA copies (**Fig. 4a**). At variant allele fractions below 0.5, observed/expected was 1.53–1.84 in TCGA and 1.44–1.52 in PCAWG (all P ≤ 0.018). At 0.5 or above it fell to 0.84 and 0.86 (not significant). Consistent with a genuine functional cost, the heteroplasmy of high-perturbation missense mutations resembled that of truncating (loss-of-function) MT-CYB mutations (median VAF 0.32 versus 0.37; two-sided Mann–Whitney P = 0.44) and was lower than that of low-perturbation missense mutations (0.57; P = 0.006), as expected if these variants are deleterious rather than neutral. This pattern was not explained by tumour purity. It persisted with heteroplasmy corrected by consensus purity estimates (1.97 and 1.61 below 0.5; 1.04 above) or by ABSOLUTE, and in tumours with purity ≥ 0.7 (**Supplementary Fig. 6**). Median purity did not differ between tumours carrying perturbing and other MT-CYB missense variants (0.79 versus 0.79, P = 0.68). Germline variants showed the same direction: perturbing variants made up 8.3% of sites seen only in the heteroplasmic state and 5.3% of sites homoplasmic in at least one person (odds ratio 1.63, P = 0.19; **Fig. 4b**). The tumour pattern was shared across cancer types (heterogeneity P = 0.09; **Fig. 4c**). Together these results fit a threshold model (**Fig. 4d**). One interpretation is that wild-type mtDNA buffers functional defects at lower mutant fractions, whereas higher fractions become less tolerable. Cross-sectional tumour VAF is not a direct measure of a universal biochemical threshold or longitudinal purging, and it may additionally reflect mtDNA copy number and tumour subclonal composition.

**Figure 4.**
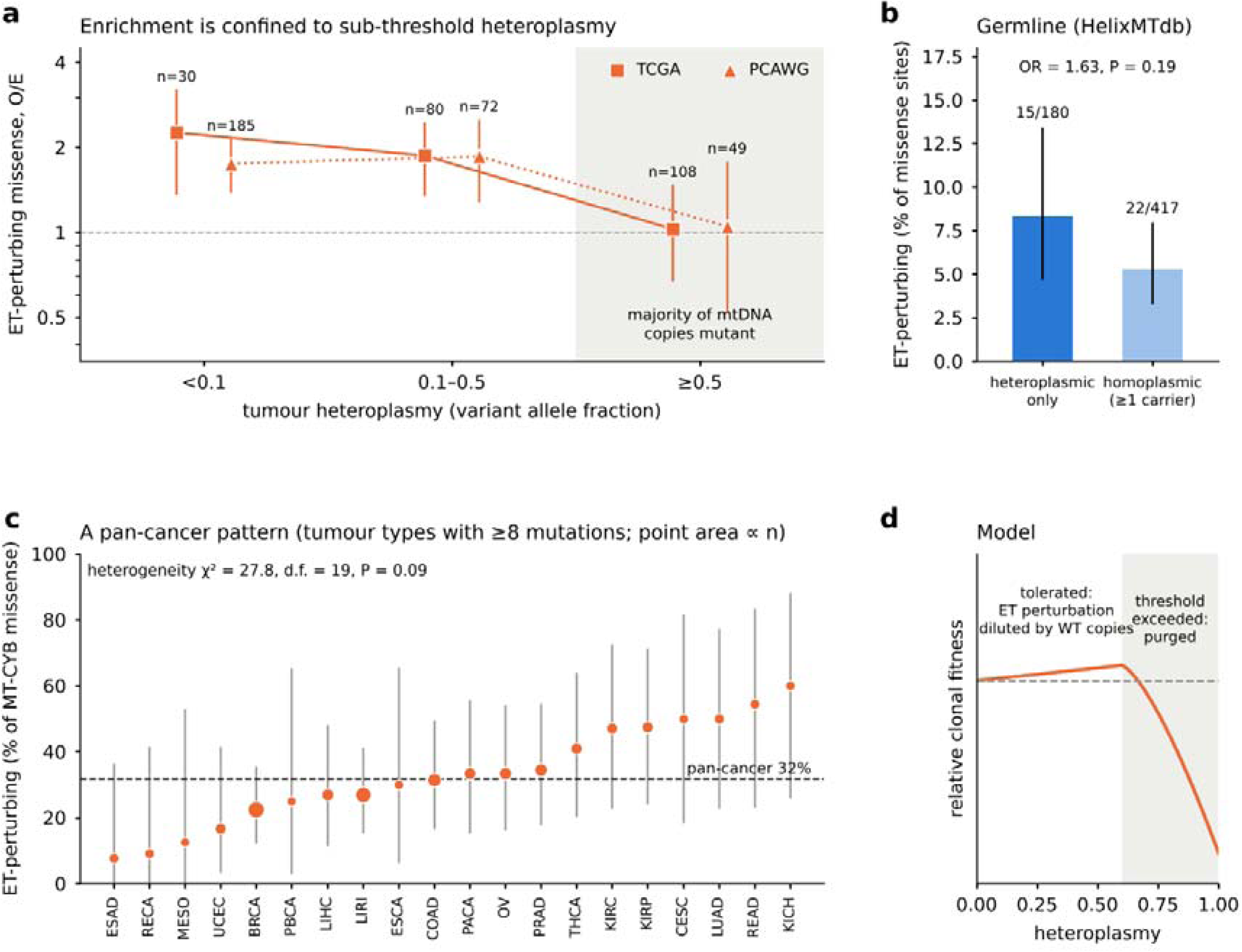
ET-perturbing MT-CYB variants are tolerated only below a heteroplasmy threshold. a, Observed/expected ratio of high-perturbation missense mutations (predicted ≥2-fold change in at least one Q-cycle ET step; Fig. 3a) among somatic MT-CYB missense mutations, stratified by tumour variant allele fraction (VAF), in TCGA (squares, solid line) and PCAWG (triangles, dotted line). The expected fraction was derived from each cohort’s strand-aware trinucleotide mutation spectrum across all possible MT-CYB missense substitutions. Shading marks strata in which most mtDNA copies carry the variant. Error bars, 95% Poisson confidence intervals; numbers give missense mutations per stratum. b, Percentage of ET-perturbing variants among germline MT-CYB missense sites in HelixMTdb observed only in the heteroplasmic state versus sites homoplasmic in at least one carrier (two-sided Fisher’s exact test). Error bars, 95% binomial confidence intervals. c, Percentage of ET-perturbing variants among somatic MT-CYB missense mutations in tumour types with at least eight mutations (TCGA and PCAWG project codes). Point area is proportional to the number of mutations. Dashed line, pan-cancer value. Heterogeneity was assessed by χ² test. d, Schematic model. ET-perturbing variants are tolerated, and accumulate, while wild-type mtDNA copies buffer the respiratory chain. Once heteroplasmy exceeds the biochemical threshold, they reduce clonal fitness and are purged. The curve is illustrative and not fitted to data. Source data are provided as a Source Data file.

### No detectable effect on nuclear oxidative mutagenesis

Increased electron leakage at the Qo site could raise mitochondrial superoxide and, downstream, oxidative damage to nuclear DNA. We tested this with single-base substitution signatures fitted to 9,382 TCGA exomes (COSMIC v3.4; median reconstruction cosine 0.88; **Supplementary Fig. 4**). Known tissue patterns were recovered, including SBS7 in melanoma, SBS4 in lung cancers and SBS18 most often in colorectal, gastric and oesophageal cancers. Among 3,744 tumours with adequate MT-CYB coverage, the 22 carrying Qo-site or ISP-interface missense variants did not show more SBS18 than tumours without MT-CYB missense variants (exposure ratio 0.58, 95% CI 0.33–1.05, P = 0.14; **Supplementary Fig. 5**). SBS36 (0.83, P = 0.55) and G>T at GG/GGG motifs (0.84, P = 0.19) were also unchanged. The upper confidence bound argues against a large increase in nuclear oxidative mutagenesis, although exome-based SBS18 detection is limited (present in 6% of tumours).

## Discussion

We find that human cancers accumulate cytochrome b (MT-CYB) variants predicted to perturb mitochondrial electron-transfer kinetics, despite strong depletion of these variants in the germline.^11^ The signal is concentrated in charge-changing substitutions near the haem b centres, ^8^ scales with the predicted magnitude of electron-transfer perturbation, and is observed across two tumour cohorts generated using different sequencing approaches. Importantly, the enrichment is confined predominantly to variants with low heteroplasmy, consistent with relaxed purifying selection rather than strong positive selection. These findings suggest that the functional constraints governing mitochondrial electron transfer, which are strongly maintained in the human germline, may be partially relaxed during tumour evolution. The biological significance of this finding relates to the distinctive role of complex III in mitochondrial bioenergetics. Whereas complexes I and II provide major routes of electron entry into the ubiquinone pool, complex III oxidizes ubiquinol supplied by these and other metabolic pathways through the bifurcated Q cycle. MT-CYB coordinates haem bL and bH and forms much of the structural framework of the quinone-binding sites, making its electron-transfer kinetics sensitive to changes in local electrostatics. Previous pan-cancer studies identified distinct mtDNA mutation patterns across respiratory complexes, including enrichment of truncating alterations in subsets of complex I genes and predominance of missense variation in complex III. Our findings extend these observations by linking differential mutational constraint to predicted perturbations of haem electrostatics and electron-transfer kinetics. Such changes could potentially influence CoQ redox balance, mitochondrial respiration, pyrimidine biosynthesis and ROS signalling. However, these functional consequences remain hypothetical, as mitochondrial metabolism and tumour-cell fitness were not directly assessed. The heteroplasmy-dependent distribution provides insight into the selective constraints acting on these variants. High-perturbation mutations were enriched at variant allele fractions below 0.5, but not at higher fractions, and their distribution resembled that of truncating MT-CYB mutations. This pattern is consistent with wild-type mtDNA partially buffering deleterious effects at low heteroplasmy, whereas increasing mutant fractions may compromise respiratory function and cellular fitness. ^12,13^ Such a threshold-dependent model is compatible with established mitochondrial genetic principles and strong germline purifying selection. ^14–17^ Nevertheless, tumour variant allele fraction does not directly establish a biochemical threshold, and cross-sectional data cannot demonstrate longitudinal elimination of deleterious variants. Thus, the findings support relaxed purifying selection without excluding context-dependent adaptive effects. An important mechanistic limitation is that predicted electron-transfer perturbations are strongly correlated with local electrostatic changes (r = 0.94). Although individual electron-transfer steps involve quantum-mechanical electronic coupling, the modelled effects of MT-CYB substitutions arise predominantly from shifts in haem redox potentials and thermodynamic driving forces. After adjustment for electrostatic and physicochemical variables, the tunnelling-specific component was not statistically significant. Therefore, our findings do not establish that quantum tunnelling itself is under selection, but rather identify relaxed constraints on variants predicted to alter the electrostatic and kinetic determinants of electron transfer. Direct measurement of haem midpoint potentials and electron-transfer kinetics, particularly for variants with divergent predictions such as G116D, G48E and G34D versus R80H and E202K, will be essential to distinguish these mechanisms. Our initial hypothesis that Qo-site mutations might increase electron leakage and promote nuclear oxidative mutagenesis was not supported. Tumours carrying Qo-site or ISP-interface variants showed no significant increase in SBS18, SBS36 or G>T substitutions at GG/GGG motifs.^10,18–20^ Although these results argue against a substantial increase in the measured nuclear oxidative-mutation signatures, limited sample size and exome-based detection sensitivity constrain this conclusion. Moreover, the absence of nuclear oxidative mutagenesis does not exclude localized mitochondrial ROS signalling, which requires direct functional investigation. Several limitations warrant consideration. The analysis relies on a single cryo-EM structure and empirical Pathways and screened-Coulomb models that do not explicitly account for conformational dynamics, protonation states, quinone occupancy or local electrostatic heterogeneity. ^21^ Predicted rate changes should therefore be interpreted primarily as relative estimates. Although tumour-purity correction did not eliminate the heteroplasmy-dependent enrichment, mtDNA copy number and clonal structure were not directly modelled. The analogous complex IV analysis did not reproduce tumour enrichment, ^22–24^ but differences in electron-transfer architecture and modelling scope, together with the absence of an equivalent analysis of complex I, preclude definitive claims of complex III specificity. Furthermore, the enrichment was not reproduced in the MSK-IMPACT targeted-panel cohort (539 MT-CYB missense mutations; haem-environment charge-change O/E = 1.10, P = 0.64; tunnelling-perturbation O/E = 0.94, P = 0.54). This cohort’s heteroplasmy distribution (median VAF 0.5) under-samples the low-variant-allele-fraction range in which the signal resides; confirmation in independent cohorts with comparable sensitivity is needed. ^5,25^ Future studies combining direct electron-transfer measurements with mitochondrial functional assays and heteroplasmy-controlled cellular models should determine whether MT-CYB variants are merely tolerated or confer context-dependent adaptive effects. In conclusion, our findings connect cancer-associated differences in mitochondrial mutational constraint to structure-based predictions of electron-transfer perturbation. MT-CYB variants predicted to disrupt haem electrostatics and electron-transfer kinetics are strongly constrained in the germline but relatively tolerated in cancers at low heteroplasmy. This supports relaxed purifying selection on mitochondrial electron-transfer-associated functions without establishing selection on quantum tunnelling itself or a direct tumour fitness benefit. More broadly, our study provides a biophysical framework linking mitochondrial bioenergetics to somatic evolution and metabolic adaptation in cancer.

## Methods

### Structure and pocket detection

We used the cryo-EM structure of human kidney respiratory complex III (PDB 9CG3, 2.96 Å). ^9^ The cytochrome b chain was confirmed to match the rCRS translation of MT-CYB (NC_012920, positions 14,747–15,887, vertebrate mitochondrial code) at all 378 modelled residues (2–379; residues 1 and 380 are not resolved in the map). ^26,27^ Hydrogens and waters were removed. Because no quinone or inhibitor is modelled, the Qo and Qi pockets were detected geometrically: grid points on a 1-Å lattice that were ≥3.0 Å from any heavy atom and had ≥60 atoms within 8 Å were clustered by connectivity; the cluster adjacent to haem bL lined by Glu271 was assigned Qo, ^28–30^ and the cluster adjacent to haem bH was assigned Qi. Haem bL and bH and their ligating histidines, and the CuA, haem a and haem a3 centres of complex IV (PDB 5Z62), were identified from metal–ligand distances. ^31,32^

### Pathways couplings

Electronic couplings were computed with the Beratan–Onuchic Pathways model on heavy atoms. ^33,34^ Each atom was a node; edges were covalent (decay 0.6, including metal–ligand bonds), hydrogen-bond (0.6²·exp[−1.7(R−2.8)] for N/O pairs 2.5–3.5 Å) or through-space (0.5·0.6·exp[−1.7(R−1.4)] within 6 Å). ^35,36^ Edge weights were −ln(decay); the optimal coupling between donor and acceptor cofactor atoms was the Dijkstra shortest path. Per-residue importance was the drop in optimal coupling on removing a residue’s side-chain atoms. Leakage couplings to the quinone pockets used pocket grid points as through-space acceptors.

### Marcus electron-transfer perturbation score

For each possible missense substitution we estimated the change in net residue charge (D, E = −1; K, R = +1; H = +0.1 at pH 7.4) and the screened Coulomb shift it imposes on each redox centre, ΔEm = 14.4·Δq/(ε·r) eV (r = side-chain-centroid to centre, floored at 3 Å). Each electron-transfer step’s driving-force change was the acceptor-minus-donor differential. Rate changes followed the Marcus non-adiabatic expression, log10(k/k_WT) = −[(ΔG+λ)²−(ΔG0+λ)²]/(4λk_BT·ln10), at 310 K. ^37^ Reference values were ΔG0 = −0.10 eV and λ = 0.70 eV (bL→bH), ΔG0 = −0.20 eV and λ = 0.80 eV (Qo→bL), ΔG0 = −0.02 eV and λ = 0.70 eV (bH→Qi), and ΔG0 = −0.05 eV and λ = 0.40 eV (CuA→haem a).^8,21,22^ The bL→bH coupling change from side-chain truncation was added. The Q-cycle score was the root-sum-square of the three rate changes, with per-step and composite scores capped at 3 log10 units; high-perturbation variants were those changing ≥1 step by ≥2-fold (|Δlog10 k| ≥ 0.3). Sensitivity was assessed over ε = 4, 8 and 16 and λ scaled by 0.7×, 1× and 1.4× the reference value of each step.^38,39^

### Variant data

Somatic mtDNA mutations for TCGA (10,132 exomes) and PCAWG (2,536 genomes), and HelixMTdb germline variant sites, were taken from Gorelick et al. (github.com/reznik-lab/mtdna-mutations).^5,6,40,41^ Single-nucleotide variants were used; the reference allele was confirmed against rCRS. Somatic calls at sites carried by more than ten HelixMTdb individuals were removed as likely germline polymorphisms. ^42^ Variant consequences were annotated in the vertebrate mitochondrial code and cross-checked against the published HGVSp (100% agreement). ^43,44^ Patients shared between TCGA and PCAWG were identified by barcode (789 patients); pooled estimates were confirmed to be robust to removing the overlapping PCAWG samples.

### Null model and classification

Expected counts were derived from each cohort’s own 192-channel strand-aware trinucleotide mutation spectrum, ^6,7,42^ estimated from all mtDNA single-nucleotide variants excluding missense variants in the analysed genes. Crucially, because the polymorphism filter preferentially removes tolerated sites, expected counts for the filtered somatic calls were computed over the same restricted universe of sites; germline comparisons retained the full site universe, because the polymorphism filter is defined by germline carrier counts and would otherwise remove the germline variants themselves. Residues 1 and 380 are not resolved in 9CG3, so the five germline missense variants at these residues were excluded from the structure-based analyses (**Figs 1, 3** and **4**), which therefore comprise 597 germline sites and 524 somatic mutations. Omitting this restriction inflated enrichment by up to ∼50%. The model was validated on synonymous variants. ^45,46^ Residues were classified hierarchically as Qo site, ISP interface (within 4.5 Å of the Rieske subunit), Qi site, haem b environment or other. ^47^ Charge-changing substitutions were those with |Δq| ≥ 0.9.

### Nuclear mutational signatures

For 30 TCGA cancer types, somatic single-nucleotide variants (GDC ensemble-masked MAFs) ^40,48^ were converted to SBS96 counts using the MAF CONTEXT field and fitted to COSMIC v3.4 GRCh38 exome signatures (non-artefact subset) by non-negative least squares with backward elimination, ^49^ retaining one tumour per patient. G>T substitutions at guanines within GG/GGG runs were counted as a sequence-level proxy for hole-trapping oxidative damage. ^18,20,50^ Analyses were restricted to tumours with ≥90% callable MT-CYB sequence. Tumour purity used the consensus purity estimate and ABSOLUTE from Aran et al.; ^51^ cancer-cell heteroplasmy was approximated as VAF/purity capped at 1. ^52^

### Statistics

Enrichment used two-sided exact binomial or Poisson tests against spectrum-derived expectations, with binomial or Poisson confidence intervals. Somatic-versus-germline class shares used Fisher’s exact tests. The ET-specific component was tested in quasi-Poisson regression with spectrum offsets, adjusting for electrostatic shift, charge change, Kyte–Doolittle hydropathy change, side-chain volume change, burial and distance to the nearest redox centre, with the score orthogonalized to electrostatic shift and charge change. Signature associations used within-cancer-type permutation (10,000 permutations) with stratified bootstrap intervals; SBS18 presence also used a Cochran–Mantel–Haenszel test. Cross-cancer heterogeneity used a χ² test. Heteroplasmy distributions across functional groups were compared by two-sided Mann–Whitney U tests, and co-occurrence of complex I truncating mutations by Fisher’s exact test. Across the primary confirmatory tests—germline depletion and tumour enrichment for the haem-environment charge-change and tunnelling-perturbation classes, the per-class relaxed-constraint odds ratios, and the heteroplasmy-threshold contrast—P values were controlled for multiple testing by the Benjamini–Hochberg procedure; all remained significant at a 5% false-discovery rate (q ≤ 0.023).

## Ethics statement

This study is a secondary analysis of publicly available, de-identified data: somatic mtDNA variant calls from TCGA and PCAWG, aggregate germline variant frequencies from HelixMTdb, open-access TCGA somatic mutation files from the NCI Genomic Data Commons, and published tumour purity estimates. No new human participants were recruited and no individual-level identifiable data were accessed, so additional institutional review board approval and informed consent were not required. The original studies obtained informed consent and ethical approval as described in their publications.

## Data availability

Somatic mtDNA mutation calls for TCGA (10,132 exomes) and PCAWG (2,536 genomes) and HelixMTdb germline variant sites were obtained from Gorelick et al. ^5^ (https://github.com/reznik-lab/mtdna-mutations). Nuclear somatic single-nucleotide variants for TCGA were obtained as open-access ensemble-masked MAF files from the NCI Genomic Data Commons (https://portal.gdc.cancer.gov). Tumour purity estimates (consensus purity estimate and ABSOLUTE) were taken from Aran et al. ^51^ The cryo-EM structures of human respiratory complex III and complex IV are available from the Protein Data Bank under accession codes 9CG3 and 5Z62. The revised Cambridge Reference Sequence is available from NCBI under accession NC_012920. COSMIC v3.4 mutational signatures are available from https://cancer.sanger.ac.uk/signatures. MSK-IMPACT mtDNA variant calls were obtained from Gorelick et al. ^5^ Source data are provided with this paper.

## Code availability

All custom code used for pocket detection, Pathways coupling calculations, Marcus electron-transfer perturbation scoring, null-model construction, mutational signature fitting and statistical analyses is provided as Supplementary Software with this paper.

## Acknowledgements

We thank the TCGA Research Network, the ICGC/TCGA Pan-Cancer Analysis of Whole Genomes (PCAWG) Consortium and the Helix team for making their data publicly available, and the authors of Gorelick et al. for releasing their mtDNA mutation calls. This work was supported by a grant from the Korean ARPA-H Project through the Korea Health Industry Development Institute (KHIDI), funded by the Ministry of Health & Welfare, Republic of Korea (grant number RS-2025-25456722), and by the ANCHOR through the Seoul ANCHOR Center, funded by the Ministry of Education (MOE) and the Seoul Metropolitan Government (2026-ANCHOR-01-022-05), to J.-H.C.

## Author contributions

**J.-Y.S.** conceived the original study, designed and performed the structural, electron-tunnelling and selection analyses, generated the figures and drafted the manuscript. **J.-H.C.** conceived the original concept linking mitochondrial quantum mechanical bioenergetics to overarching biology and provided the biological framework for the study. **J.-Y.S.** and **J.-H.C.** supervised the study, contributed to the interpretation of the results and revised the manuscript. Both authors read and approved the final manuscript.

## Supplementary Figures

**Supplementary Fig. 1.**
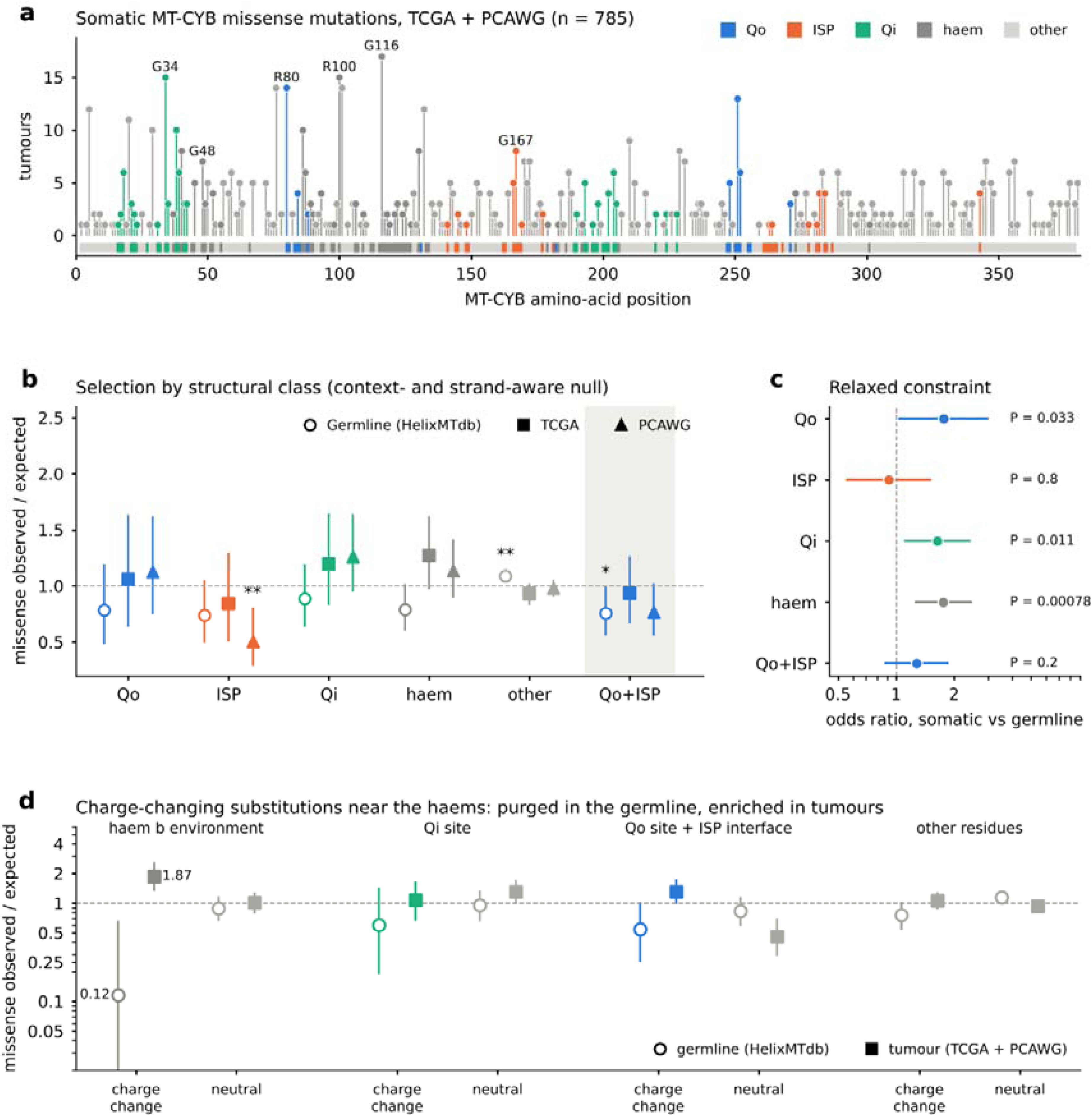
Selection by structural class without the population- polymorphism filter. As Fig. 2 but including all somatic MT-CYB missense calls (n = 785), without removing sites carried by more than ten HelixMTdb individuals. a, Somatic MT-CYB missense mutations coloured by structural class. b, Observed/expected ratio of missense mutations by class for germline sites (open circles), TCGA (squares) and PCAWG (triangles). c, Odds ratio for each class’s share of missense variants, somatic versus germline. d, Observed/expected split by class and by charge change (|Δq| ≥ 0.9); haem-environment charge-changing substitutions remained depleted in the germline (O/E = 0.12) and enriched in tumours (O/E = 1.87 pooled), confirming that the main-text result does not depend on the polymorphism filter. Error bars as in Fig. 2.

**Supplementary Fig. 2.**
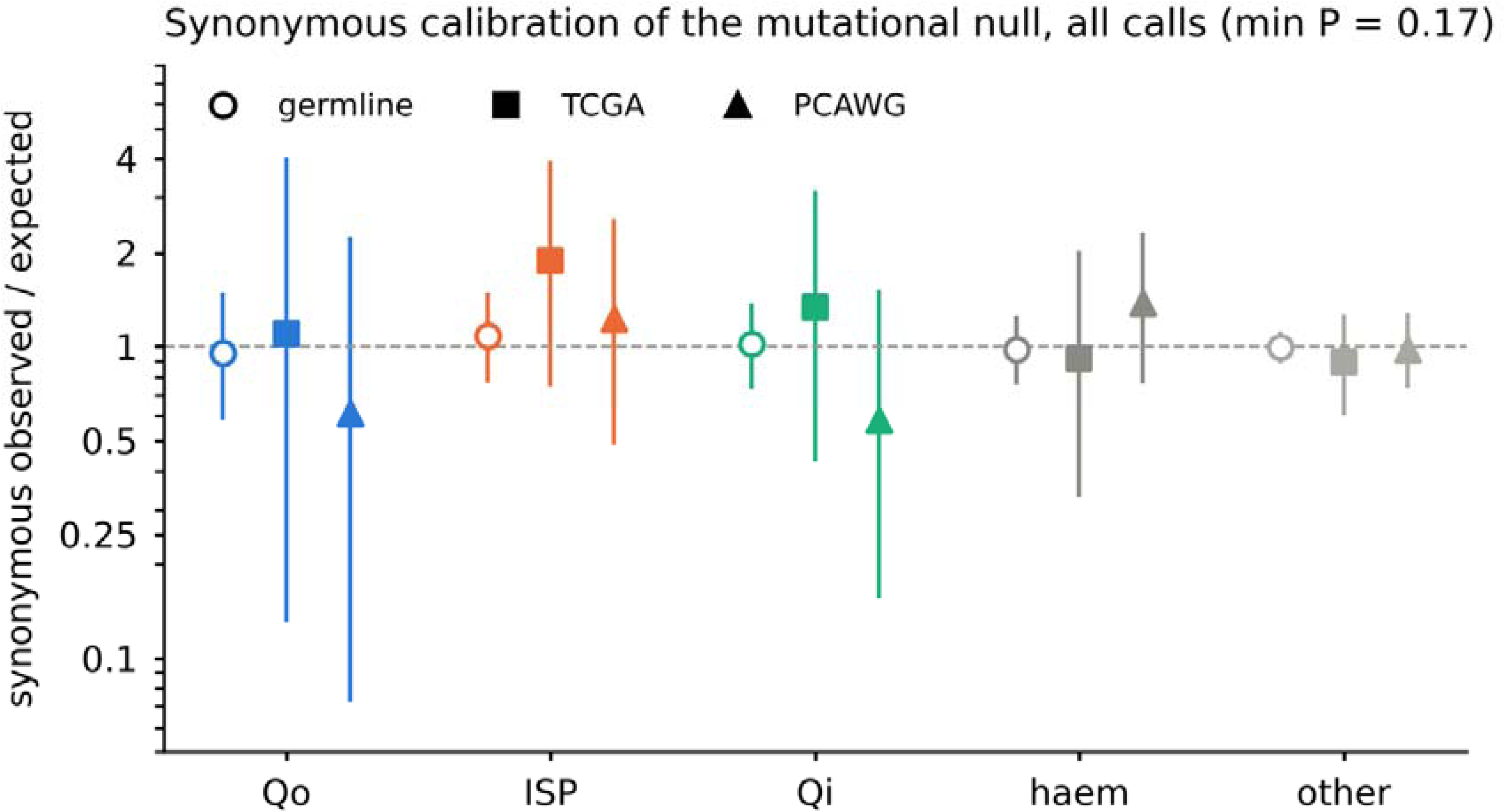
Synonymous calibration of the mutational null model. Observed/expected ratio of synonymous MT-CYB variants by structural class for germline sites (open circles), TCGA (squares) and PCAWG (triangles), using the same strand-aware trinucleotide null and site universe as the missense analysis. No class deviated from expectation in any cohort (minimum P = 0.17), indicating that the null model is well calibrated and does not itself generate the haem-environment signal. Error bars, 95% Poisson confidence intervals.

**Supplementary Fig. 3.**
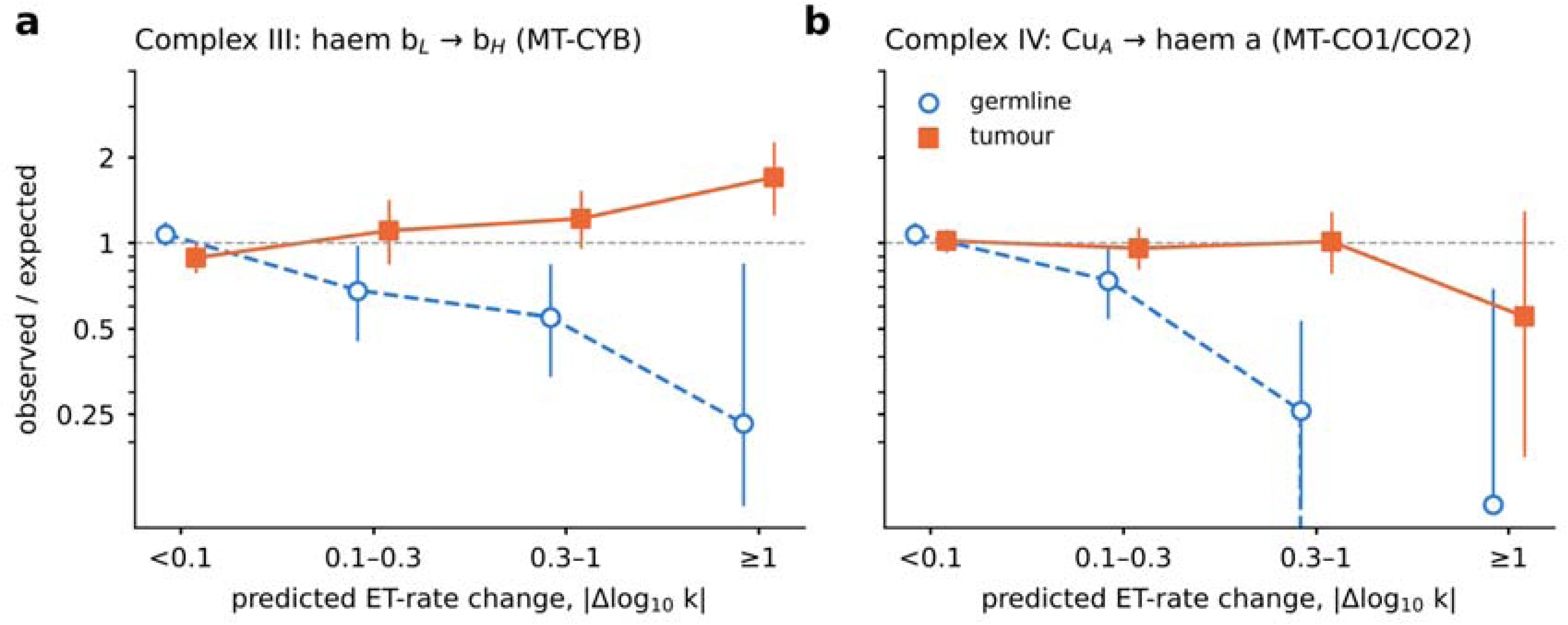
The electron-transfer selection signal of complex III is not reproduced in complex IV. Observed/expected ratio across bins of predicted electron-transfer rate change for germline sites (open circles, dashed) and pooled tumours (squares, solid). a, Complex III haem bL→bH step (MT-CYB), scored with the single-step bL→bH model to match the complex IV step (not the three-step Q-cycle score of Fig. 3b). b, Complex IV CuA→haem a step (MT-CO1, MT-CO2). The germline depletion of high-perturbation variants is present in both complexes, but the tumour enrichment is specific to complex III. Error bars, 95% Poisson confidence intervals.

**Supplementary Fig. 4.**
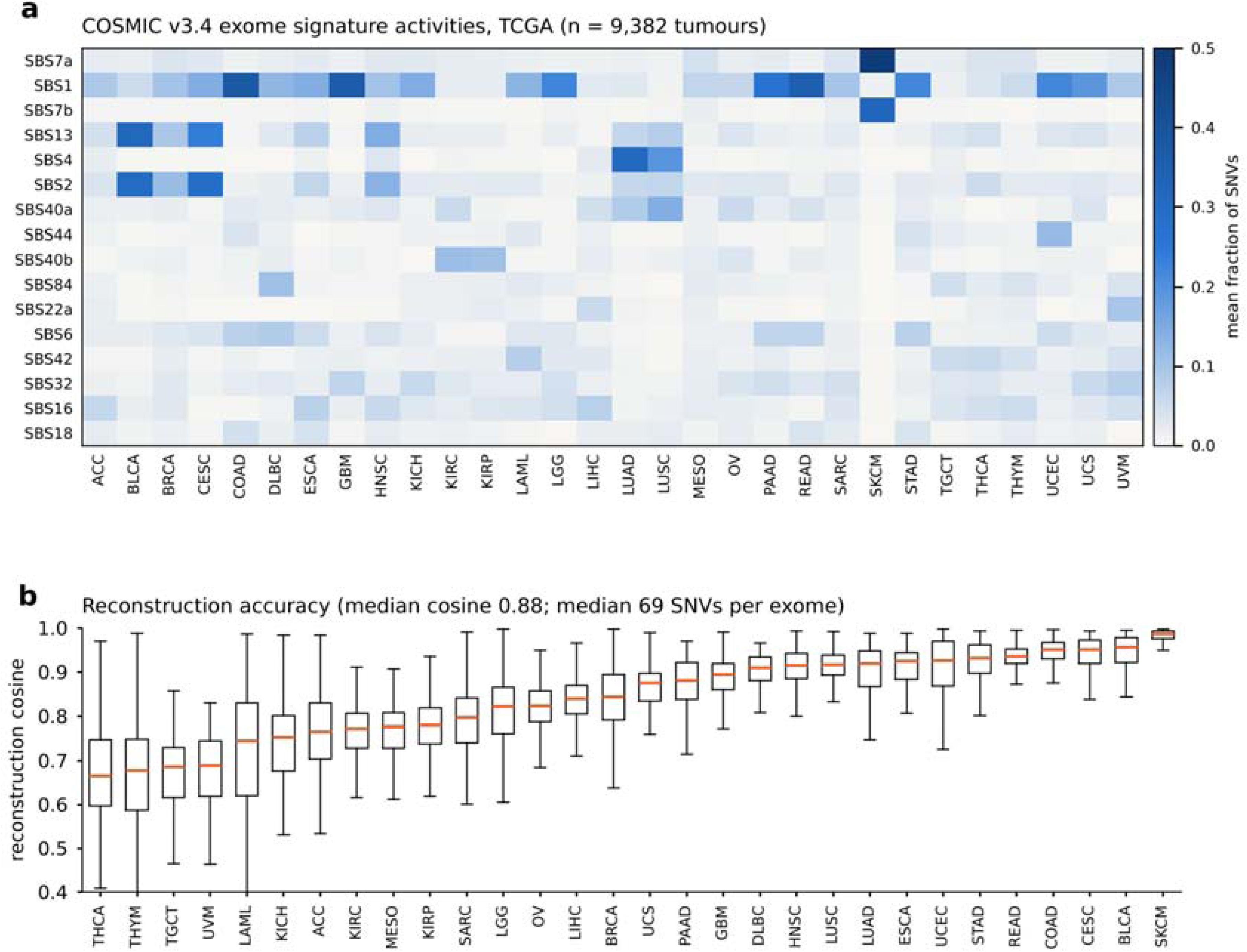
Nuclear single-base substitution signature fitting in TCGA. a, Mean per-tumour exposure (fraction of single-nucleotide variants) for the most active COSMIC v3.4 exome signatures across 9,382 TCGA tumours from 30 cancer types. Expected tissue patterns are recovered, including SBS7a/b in melanoma (SKCM), SBS4 in lung cancers (LUAD, LUSC) and the APOBEC signatures SBS2/SBS13 in bladder, breast and cervical cancers. b, Per-tumour reconstruction cosine similarity by cancer type (median 0.88; median 69 SNVs per exome). Boxes, interquartile range; whiskers, 1.5× IQR.

**Supplementary Fig. 5.**
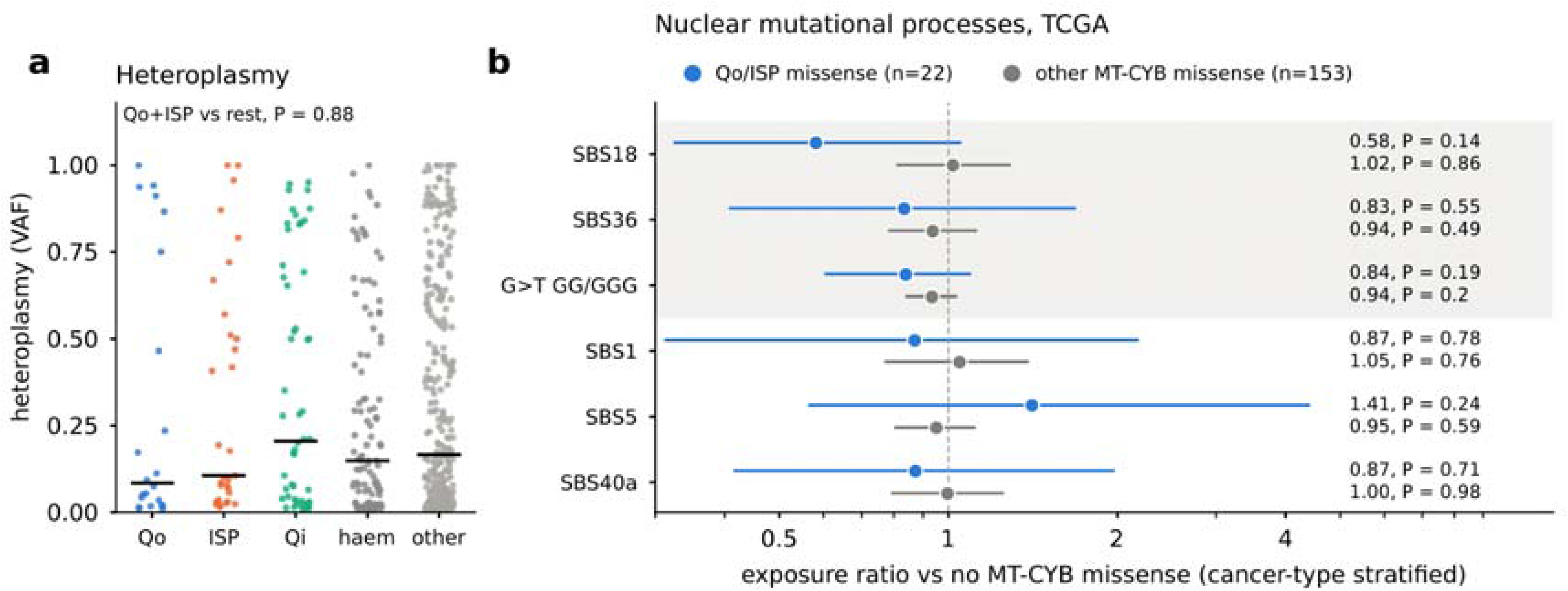
Heteroplasmy and nuclear mutational signatures in tumours with MT-CYB missense mutations. a, Variant allele fraction of somatic MT-CYB missense mutations by structural class. Horizontal lines, medians. Qo and ISP-interface mutations did not differ from other classes (two-sided Mann–Whitney U test, P = 0.88). b, Nuclear single-base substitution signature exposures in 3,744 TCGA tumours with ≥90% callable MT-CYB sequence, comparing tumours carrying Qo/ISP-interface missense mutations (n = 22) or other MT-CYB missense mutations (n = 153) with tumours lacking MT-CYB missense mutations. Exposure ratios are log-ratio differences relative to non-oxidative mutations, centred within cancer type. P values are from 10,000 within-cancer-type label permutations, and confidence intervals are from stratified bootstrap. Shading marks oxidative-damage readouts (SBS18, SBS36, G>T at GG/GGG). No oxidative signature was associated with MT-CYB mutation status.

**Supplementary Fig. 6.**
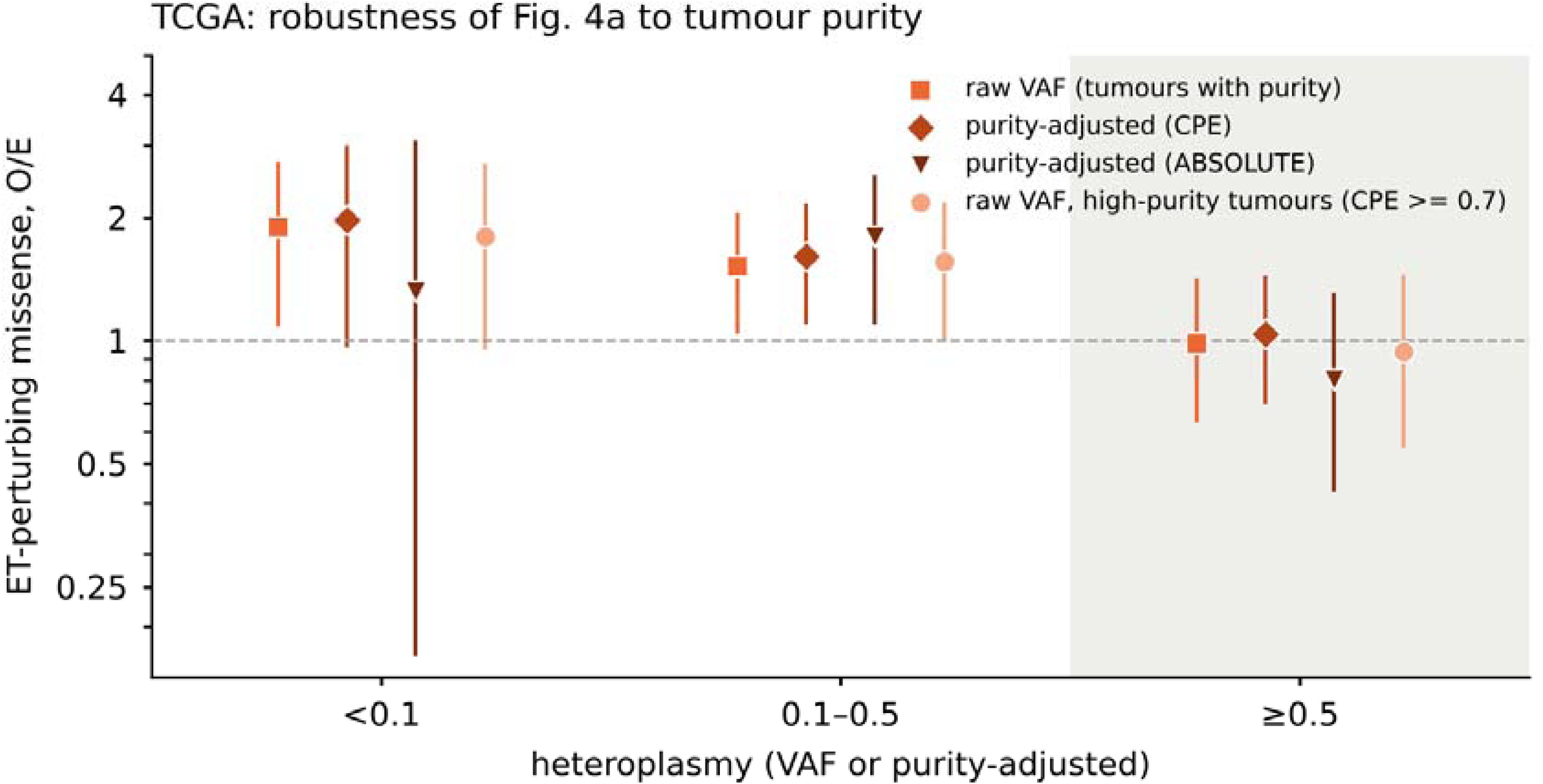
The heteroplasmy threshold is robust to tumour purity (TCGA). Observed/expected ratio of high-perturbation MT-CYB missense mutations stratified by heteroplasmy, computed with raw variant allele fraction, with VAF corrected for tumour purity using the consensus purity estimate (CPE) or ABSOLUTE, and in high-purity tumours (CPE ≥ 0.7). Enrichment below a variant allele fraction of 0.5 persists under every purity correction and is absent above 0.5 (shaded). Error bars, 95% Poisson confidence intervals.

