## Supplementary figures and images for "Relaxed evolutionary constraints on mitochondrial electron-transfer-associated variants in human cancers"

### SuppleFig1

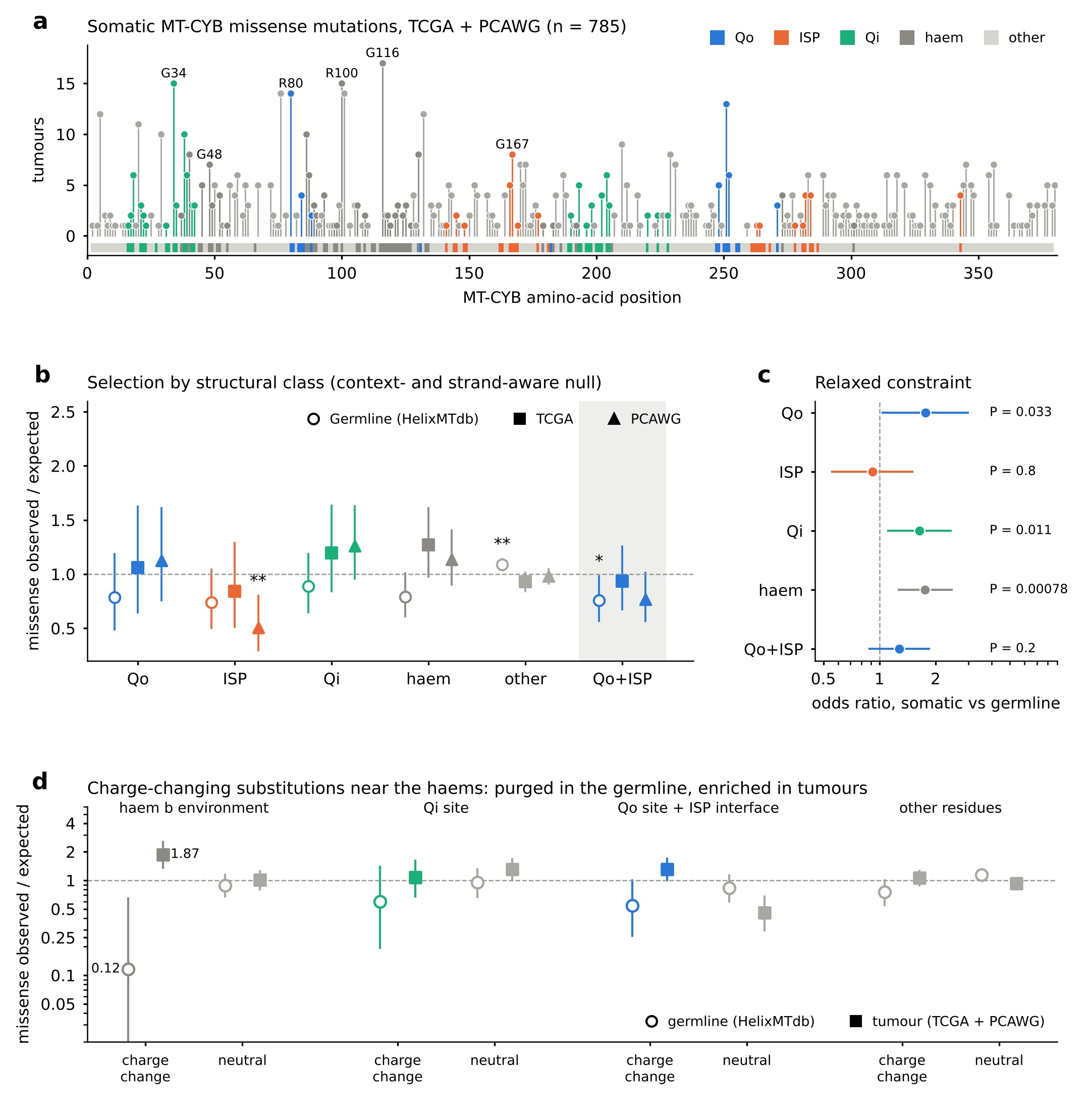

### SuppleFig2

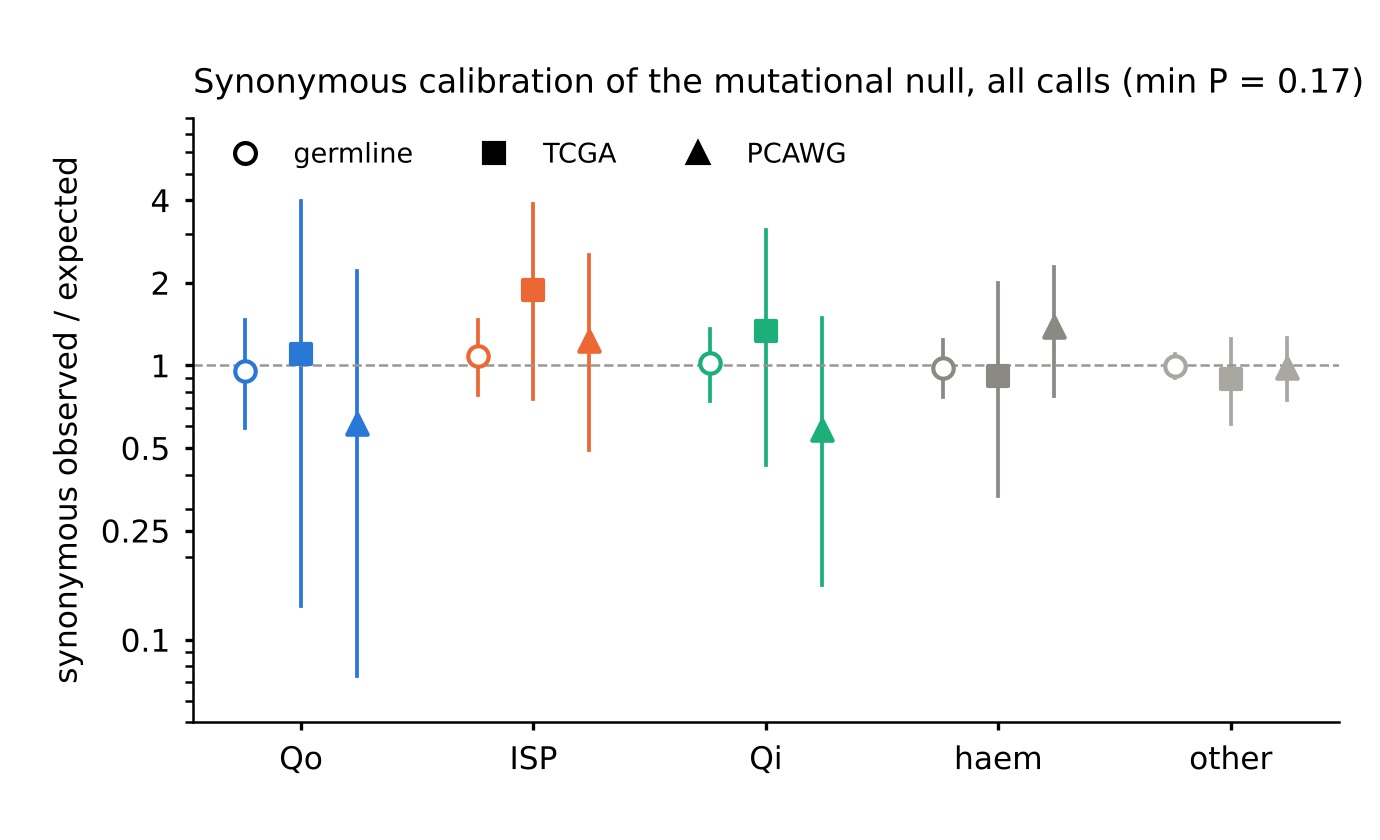

### SuppleFig3

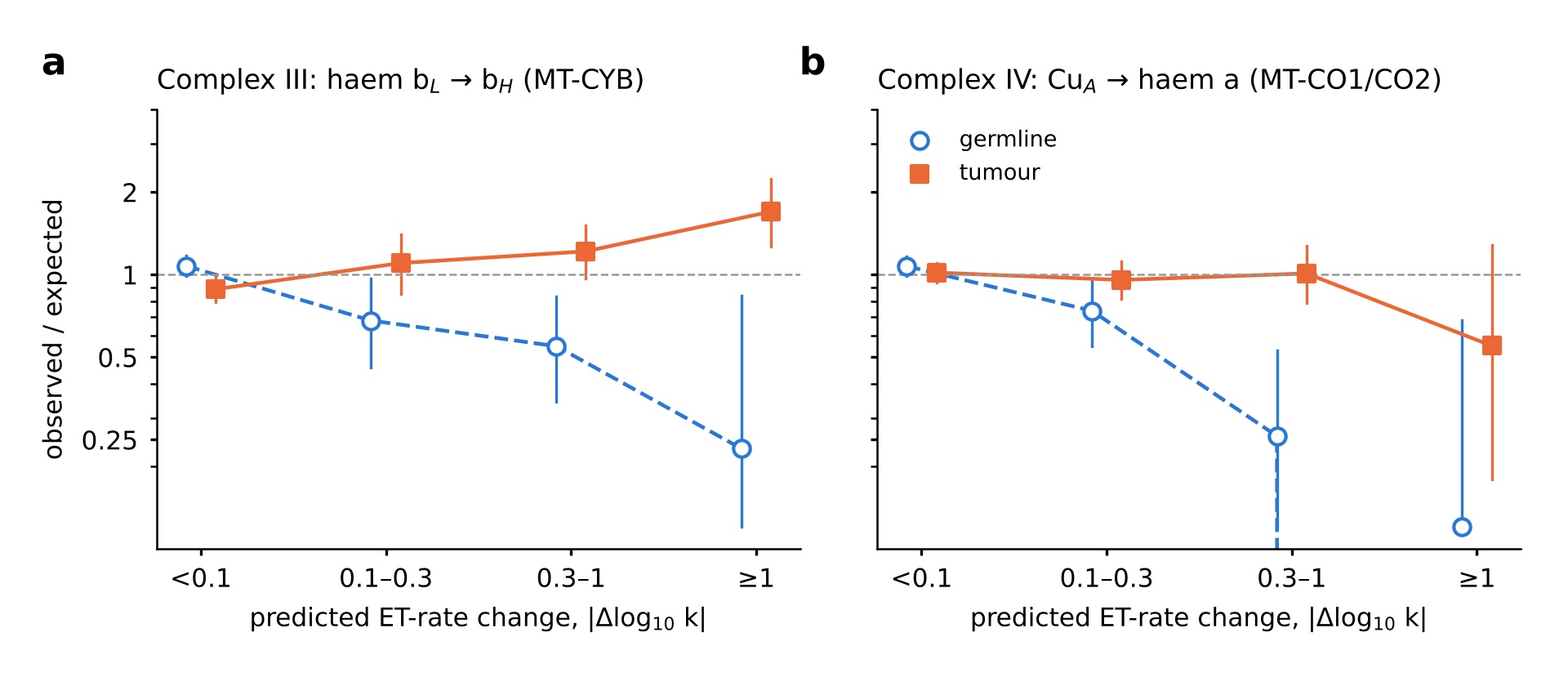

### SuppleFig4

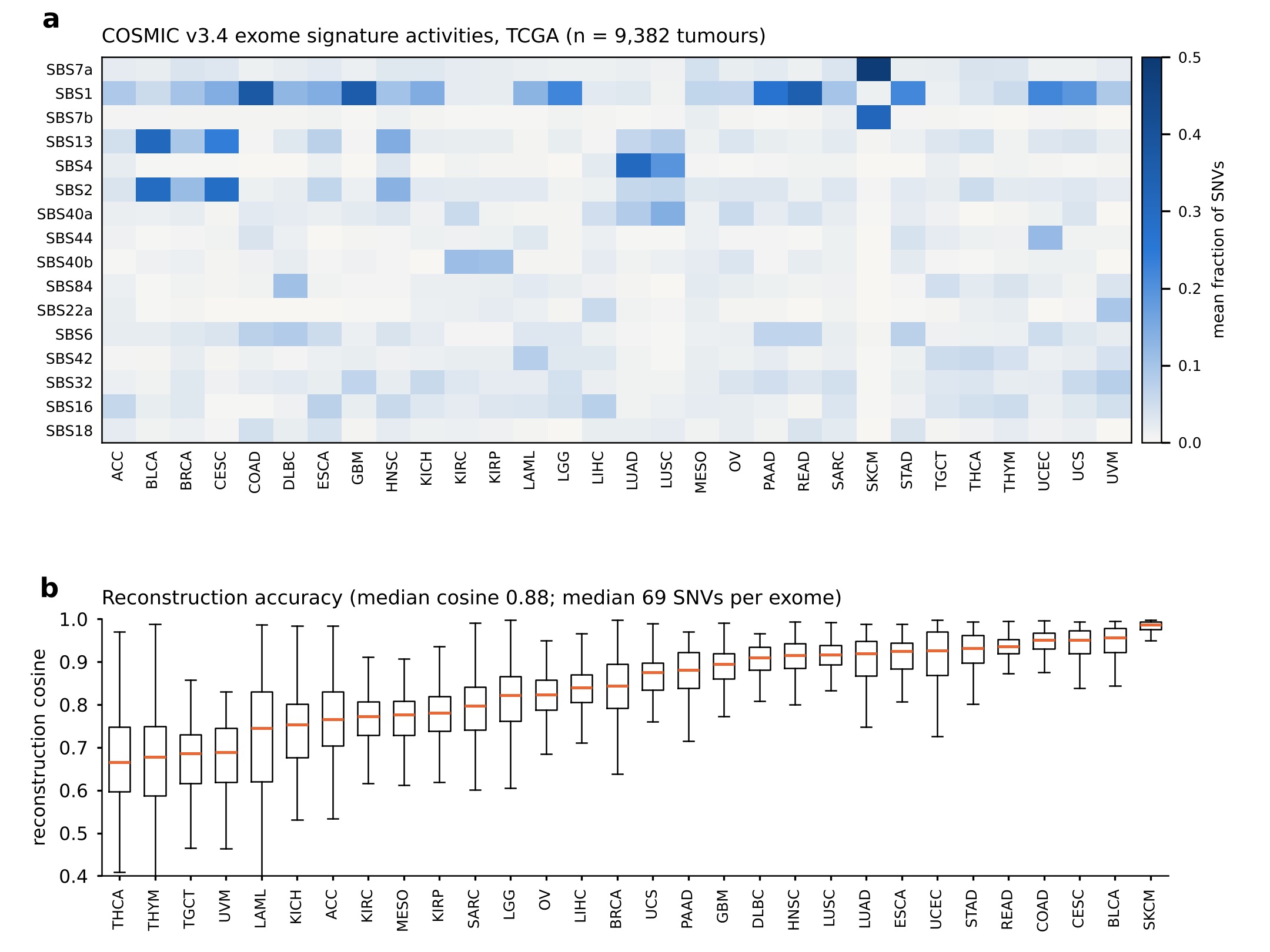

### SuppleFig5

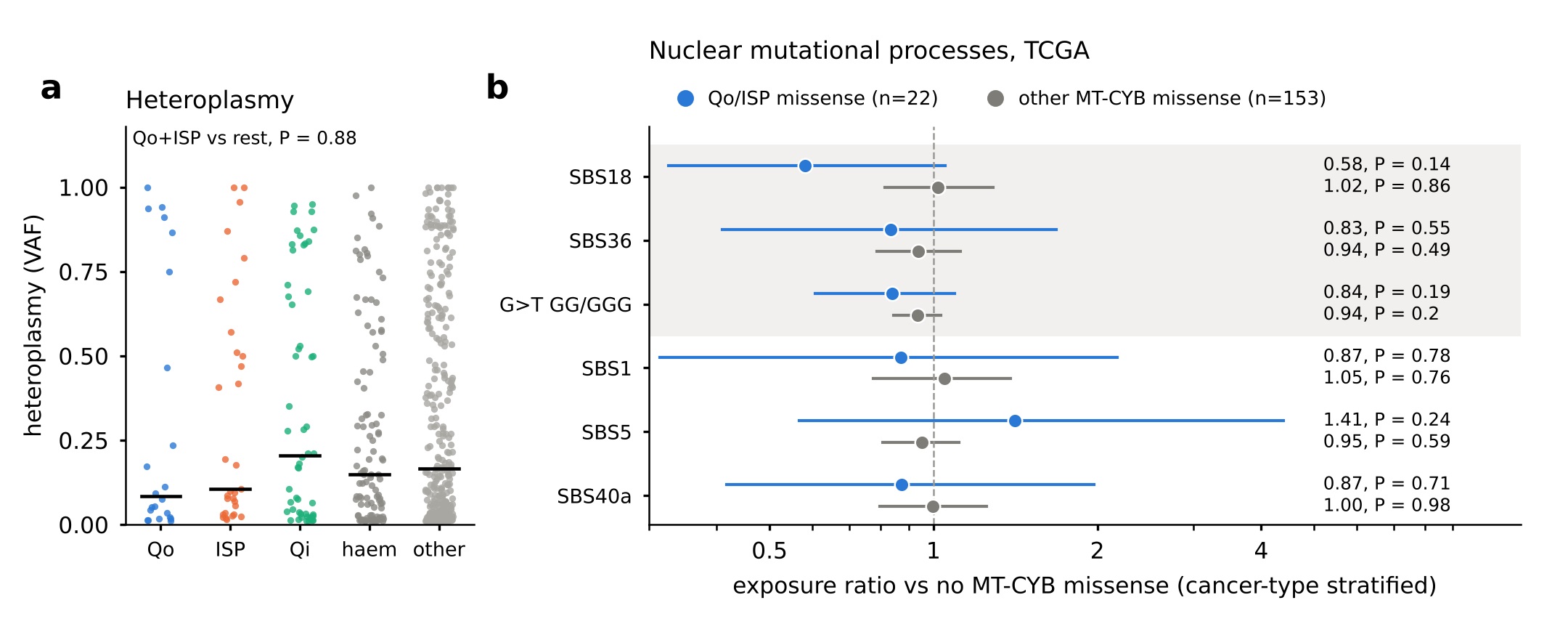

### SuppleFig6

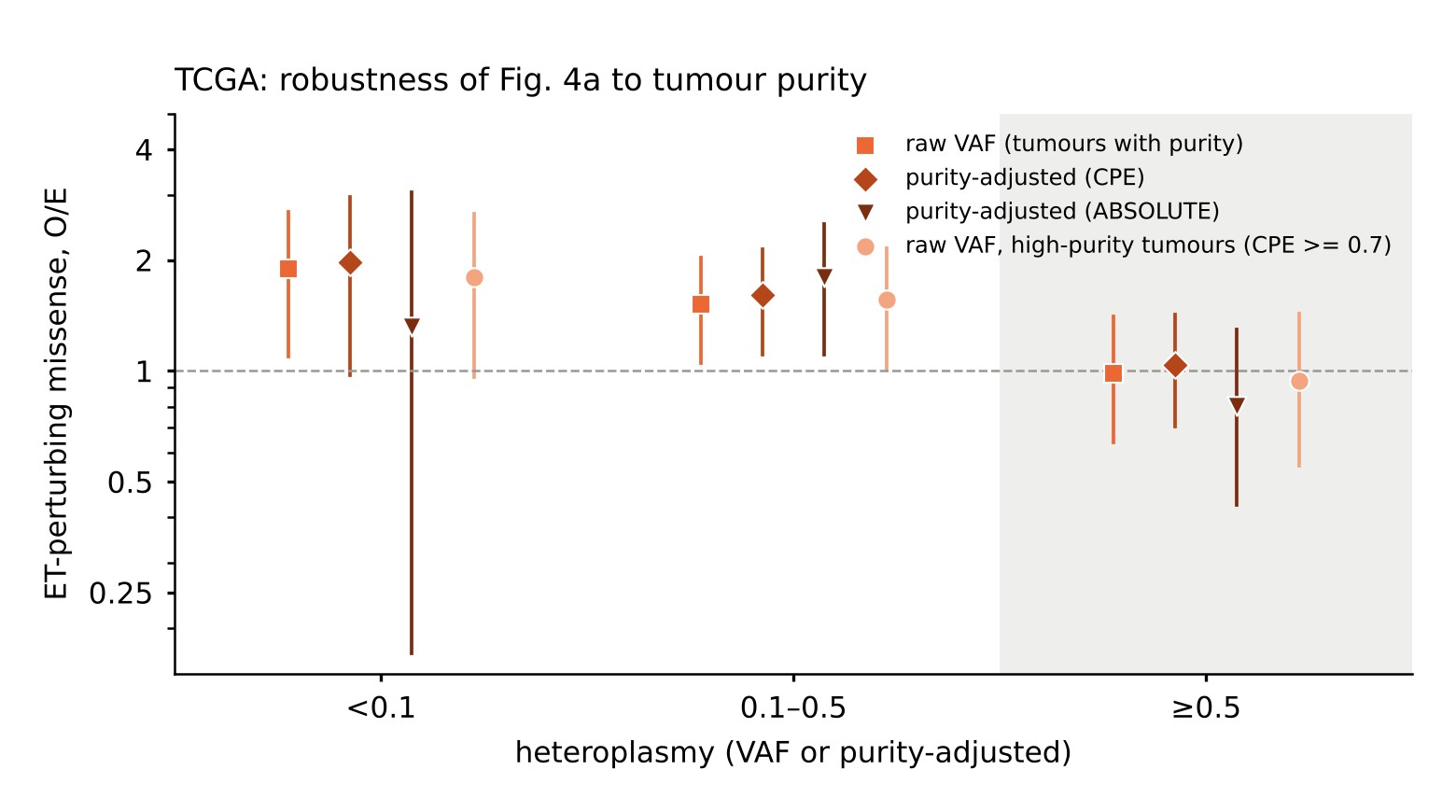
