## Supplementary code for "Relaxed evolutionary constraints on mitochondrial electron-transfer-associated variants in human cancers": README.md

# Exploratory analyses (not reported in the manuscript)

`survival_os.py` fits an overall-survival Cox model (strata = cancer type) and a log-rank test for
tumours carrying ET-perturbing MT-CYB missense mutations, using TCGA-CDR (`MTCYB_EXT`).

It was run while developing the study and is kept here for completeness. It is **not** part of the
manuscript: no survival result is reported, and nothing in the paper depends on it. It is excluded
from `run_all.sh` for that reason.

    export MTCYB_DATA=... MTCYB_EXT=... MTCYB_OUT=...
    python3 ../src/../exploratory/survival_os.py

Note the MITOMAP-overlap block printed by `src/functional_evidence.py` (section 3) is likewise
descriptive and is not reported in the manuscript.
