## Supplementary code for "Relaxed evolutionary constraints on mitochondrial electron-transfer-associated variants in human cancers": survival_os.py

```
"""Exploratory overall-survival analysis: do TCGA patients carrying a high-ET-perturbation MT-CYB
missense mutation differ in prognosis? Cox PH stratified by cancer type, adjusted for age; log-rank.
TCGA-CDR survival (Liu et al. 2018). Secondary contrast restricts to MT-CYB missense carriers to
remove the 'has any mtDNA mutation' confounder."""
import os as _os, sys as _sys
_sys.path.insert(0, _os.path.dirname(_os.path.abspath(__file__)))
from config import DATADIR, DATAROOT, EXTDIR, STRUCTDIR, ASSETDIR, OUTDIR

import sys
sys.path.insert(0, _os.path.dirname(_os.path.abspath(__file__)))
import numpy as np, pandas as pd
import statsmodels.api as sm
from statsmodels.duration.hazard_regression import PHReg
from statsmodels.duration.survfunc import survdiff

R = DATADIR
EXT = EXTDIR

cdr = pd.read_csv(EXT + 'TCGA-CDR-SupplementalTableS1.txt', sep='\t', low_memory=False)
cdr = cdr.rename(columns={'bcr_patient_barcode': 'PAT', 'OS.time': 'OStime',
                          'age_at_initial_pathologic_diagnosis': 'age'})
cdr = cdr[['PAT', 'type', 'age', 'OS', 'OStime']].copy()
for c in ('OS', 'OStime', 'age'):
    cdr[c] = pd.to_numeric(cdr[c], errors='coerce')
cdr = cdr.dropna(subset=['OS', 'OStime']).query('OStime > 0')

# screened TCGA patients (mtDNA dataset) = denominator
ts = pd.read_csv(R + 'data_samples_tcga.txt.gz', sep='\t', low_memory=False)
screened = set(ts.PATIENT_ID.astype(str))

# MT-CYB missense carriers with ET score
miss = pd.read_csv(OUTDIR + '/fig4_cyb_missense.tsv', sep='\t')
miss = miss[miss['class'] == 'somatic']
hi_pat = set(miss[miss.high].PATIENT_ID.astype(str))
anymiss_pat = set(miss.PATIENT_ID.astype(str))
lo_pat = anymiss_pat - hi_pat

cdr = cdr[cdr.PAT.isin(screened)].copy()
cdr['highET'] = cdr.PAT.isin(hi_pat).astype(int)
cdr['anymiss'] = cdr.PAT.isin(anymiss_pat).astype(int)
print(f'TCGA patients with survival + mtDNA screening: {len(cdr)}')
print(f'  high-ET MT-CYB missense carriers: {cdr.highET.sum()}')
print(f'  any MT-CYB missense carriers: {cdr.anymiss.sum()}')


def cox(df, cols, strata_col='type'):
    df = df.dropna(subset=cols + ['OS', 'OStime']).copy()
    X = df[cols].astype(float).values
    strata = df[strata_col].values
    mod = PHReg(df.OStime.values, X, status=df.OS.values, strata=strata)
    res = mod.fit()
    out = []
    for i, c in enumerate(cols):
        hr = np.exp(res.params[i]); lo, hi = np.exp(res.conf_int()[i])
        out.append((c, hr, lo, hi, res.pvalues[i]))
    return out, len(df)


print('\n=== Cox PH, OS ~ highET + age, stratified by cancer type (all screened TCGA) ===')
res, n = cox(cdr, ['highET', 'age'])
for c, hr, lo, hi, p in res:
    print(f'  {c:8s} HR={hr:.2f} [{lo:.2f}-{hi:.2f}] P={p:.3g}')
print(f'  n={n}')

print('\n=== Secondary: among MT-CYB missense carriers only, high-ET vs low-ET (adj age, strata type) ===')
carriers = cdr[cdr.anymiss == 1].copy()
res2, n2 = cox(carriers, ['highET', 'age'])
for c, hr, lo, hi, p in res2:
    print(f'  {c:8s} HR={hr:.2f} [{lo:.2f}-{hi:.2f}] P={p:.3g}')
print(f'  n={n2} (high-ET {carriers.highET.sum()}, low-ET {(carriers.highET==0).sum()})')

# crude log-rank high-ET vs rest
print('\n=== Log-rank (crude), high-ET vs all others ===')
chisq, p_lr = survdiff(cdr.OStime.values, cdr.OS.values, cdr.highET.values)
print(f'  chi2={chisq:.2f}, P={p_lr:.3g}')

# median OS days by group
for g, lab in [(1, 'high-ET'), (0, 'others')]:
    sub = cdr[cdr.highET == g]
    ev = sub[sub.OS == 1]
    print(f'  {lab}: n={len(sub)}, deaths={int(sub.OS.sum())}, median follow-up(d)={sub.OStime.median():.0f}')

pd.DataFrame([dict(analysis='all_screened', **dict(zip(['var', 'HR', 'lo', 'hi', 'P'], res[0]))),
              dict(analysis='carriers_only', **dict(zip(['var', 'HR', 'lo', 'hi', 'P'], res2[0]))),
              dict(analysis='logrank_crude', var='highET', HR=np.nan, lo=np.nan, hi=np.nan, P=p_lr)]
             ).to_csv(OUTDIR + '/survival_results.tsv', sep='\t', index=False)
print('\nsaved survival_results.tsv')
```
