## Supplementary code for "Relaxed evolutionary constraints on mitochondrial electron-transfer-associated variants in human cancers": detect_pockets.py

```
"""Grid-based detection of the Qo and Qi quinone pockets of cytochrome b (9CG3, monomer J).
No quinone/inhibitor is modelled in 9CG3, so pockets are found geometrically:
empty, buried grid points (probe >= 3.0 A from any heavy atom, many atoms within 8 A),
clustered by connectivity; the Qo cluster is the one adjacent to heme bL that is lined by
the PEWY Glu271, the Qi cluster the one adjacent to heme bH.
"""
import os as _os, sys as _sys
_sys.path.insert(0, _os.path.dirname(_os.path.abspath(__file__)))
from config import DATADIR, DATAROOT, EXTDIR, STRUCTDIR, ASSETDIR, OUTDIR

import json
import gemmi, numpy as np
from scipy.spatial import cKDTree
from scipy.sparse.csgraph import connected_components
from scipy.sparse import coo_matrix

st = gemmi.read_structure(STRUCTDIR + '/9CG3.cif'); st.remove_hydrogens(); st.remove_waters()
m = st[0]
recs, P = [], []
for c in m:
    for r in c:
        for a in r:
            recs.append((c.name, r.seqid.num, r.name, a.name)); P.append(a.pos.tolist())
P = np.array(P); tree = cKDTree(P)
fe = lambda ch, n: np.array([a.pos.tolist() for a in m[ch][str(n)][0] if a.name == 'FE'][0])
bL, bH = fe('J', 401), fe('J', 402)
fes = np.array([a.pos.tolist() for a in m['P']['301'][0] if a.element.name == 'Fe']).mean(0)

out = {}
for site, center in [('Qo', bL), ('Qi', bH)]:
    g = np.mgrid[-18:18.1:1.0, -18:18.1:1.0, -18:18.1:1.0].reshape(3, -1).T + center
    dmin, _ = tree.query(g)
    g = g[dmin >= 3.0]
    bur = np.array([len(x) for x in tree.query_ball_point(g, 8.0)])
    g = g[bur >= 60]
    pairs = cKDTree(g).query_pairs(1.01, output_type='ndarray')
    n = len(g)
    A = coo_matrix((np.ones(len(pairs)), (pairs[:, 0], pairs[:, 1])), shape=(n, n))
    k, lab = connected_components(A, directed=False)
    cands = []
    for c in range(k):
        pts = g[lab == c]
        if len(pts) < 8: continue
        d_heme = np.min(np.linalg.norm(pts - center, axis=1))
        lining = set()
        for idx in tree.query_ball_point(pts, 4.5):
            for i in idx:
                ch, num, rn, an = recs[i]
                lining.add((ch, num, rn))
        cyt = {(n_, r) for ch, n_, r in lining if ch == 'J'}
        cands.append(dict(n=len(pts), d_heme=d_heme, pts=pts, lining=lining, cyt=cyt,
                          glu271=(271, 'GLU') in cyt))
    if site == 'Qo':
        best = max([c for c in cands if c['glu271'] and c['d_heme'] < 12], key=lambda c: c['n'])
    else:
        best = max([c for c in cands if c['d_heme'] < 10 and not c['glu271']], key=lambda c: c['n'])
    cen = best['pts'].mean(0)
    print(f"{site}: {best['n']} grid pts (~{best['n']} A^3), min dist to heme Fe {best['d_heme']:.1f} A, "
          f"centroid-heme Fe {np.linalg.norm(cen-center):.1f} A, centroid-FeS {np.linalg.norm(cen-fes):.1f} A")
    cyt = sorted(best['cyt'])
    print("  cyt b lining:", ' '.join(f"{r}{n}" for n, r in cyt))
    other = sorted({(ch, n, r) for ch, n, r in best['lining'] if ch != 'J'})
    print("  other chains lining:", ' '.join(f"{ch}:{r}{n}" for ch, n, r in other))
    out[site] = dict(points=best['pts'].tolist(), lining=[list(x) for x in sorted(best['lining'])])
json.dump(out, open(ASSETDIR + '/pockets.json', 'w'))
```
