## Supplementary code for "Relaxed evolutionary constraints on mitochondrial electron-transfer-associated variants in human cancers": pathways_coupling.py

```
"""Beratan-Onuchic Pathways model on cryo-EM structures (heavy atoms only).

  covalent      eps_C = 0.6                       (incl. metal-ligand bonds)
  hydrogen bond eps_H = eps_C^2 * exp(-1.7(R-2.8)) (N/O pairs, 2.5 <= R <= 3.5 A)
  through space eps_S = 0.5*eps_C * exp(-1.7(R-1.4)) (R <= TS_CUT)
Edge weight = -ln(eps); best path = Dijkstra shortest path.
Donor/acceptor = metal + conjugated porphyrin core atoms (super-source/sink).
"""
import os as _os, sys as _sys
_sys.path.insert(0, _os.path.dirname(_os.path.abspath(__file__)))
from config import DATADIR, DATAROOT, EXTDIR, STRUCTDIR, ASSETDIR, OUTDIR

import re, sys, json, math
import gemmi, networkx as nx, numpy as np

TS_CUT = 6.0
EPS_C = 0.6
W_C = -math.log(EPS_C)
CORE_RE = re.compile(r'^(FE|N[A-D]|C[1-4][A-D]|CH[A-D])$')
BACKBONE = {'N', 'CA', 'C', 'O', 'OXT', 'CB'}
METALS = {'Fe', 'Cu', 'Zn', 'Mg'}

MT_GENES = {  # (pdb, chain) -> (gene, mtDNA start of CDS on rCRS, strand)
    ('9CG3', 'J'): ('MT-CYB', 14747), ('9CG3', 'V'): ('MT-CYB', 14747),
    ('5Z62', 'A'): ('MT-CO1', 5904), ('5Z62', 'B'): ('MT-CO2', 7586),
    ('5Z62', 'C'): ('MT-CO3', 9207),
}


def w_hb(r): return -math.log(EPS_C**2 * math.exp(-1.7 * (r - 2.8)))
def w_ts(r): return -math.log(0.5 * EPS_C * math.exp(-1.7 * (r - 1.4)))


def load_region(path, sites, pad=14.0):
    st = gemmi.read_structure(path)
    st.remove_hydrogens(); st.remove_waters()
    m = st[0]
    centers = []
    for ch, num in sites:
        for a in m[ch][str(num)][0]:
            centers.append(np.array(a.pos.tolist()))
    centers = np.array(centers)
    lo, hi = centers.min(0) - pad, centers.max(0) + pad
    atoms = []
    for c in m:
        for r in c:
            for a in r:
                p = np.array(a.pos.tolist())
                if np.all(p >= lo) and np.all(p <= hi):
                    atoms.append(dict(chain=c.name, resnum=r.seqid.num, resname=r.name,
                                      atom=a.name, el=a.element.name, pos=p))
    return atoms


def build_graph(atoms, exclude=frozenset()):
    idx = [i for i in range(len(atoms)) if i not in exclude]
    P = np.array([atoms[i]['pos'] for i in idx])
    from scipy.spatial import cKDTree
    tree = cKDTree(P)
    pairs = tree.query_pairs(TS_CUT, output_type='ndarray')
    G = nx.Graph()
    G.add_nodes_from(idx)
    d = np.linalg.norm(P[pairs[:, 0]] - P[pairs[:, 1]], axis=1)
    for (a, b), r in zip(pairs, d):
        i, j = idx[a], idx[b]
        ei, ej = atoms[i]['el'], atoms[j]['el']
        metal = ei in METALS or ej in METALS
        cov_cut = 2.7 if metal else (2.1 if 'S' in (ei, ej) else 1.9)
        if r <= cov_cut:
            w = W_C
        else:
            w = w_ts(r)
            if ei in ('N', 'O') and ej in ('N', 'O') and 2.5 <= r <= 3.5:
                w = min(w, w_hb(r))
        G.add_edge(i, j, weight=w)
    return G


def site_atoms(atoms, ch, num):
    out = []
    for i, a in enumerate(atoms):
        if a['chain'] == ch and a['resnum'] == num:
            if a['el'] in METALS or CORE_RE.match(a['atom']):
                out.append(i)
    return out


def with_terminals(G, D, A):
    H = G.copy()
    for i in D: H.add_edge('S', i, weight=0.0)
    for i in A: H.add_edge('T', i, weight=0.0)
    return H


def solve(atoms, donor, acceptor, label, pdb, knockout=True):
    D = site_atoms(atoms, *donor); A = site_atoms(atoms, *acceptor)
    G = build_graph(atoms)
    H = with_terminals(G, D, A)
    dS = nx.single_source_dijkstra_path_length(H, 'S')
    dT = nx.single_source_dijkstra_path_length(H, 'T')
    best = dS['T']
    path = nx.dijkstra_path(H, 'S', 'T')
    # describe path
    steps = []
    for u, v in zip(path[1:-2], path[2:-1]):
        r = float(np.linalg.norm(atoms[u]['pos'] - atoms[v]['pos']))
        w = H[u][v]['weight']
        kind = 'C' if abs(w - W_C) < 1e-9 else ('H' if atoms[u]['el'] in 'NO' and atoms[v]['el'] in 'NO' and 2.5 <= r <= 3.5 and abs(w - w_hb(r)) < 1e-9 else 'TS')
        steps.append(kind if kind == 'C' else f"{kind}({r:.2f}A)")
    path_res = []
    for i in path[1:-1]:
        a = atoms[i]; key = (a['chain'], a['resnum'], a['resname'])
        if not path_res or path_res[-1] != key: path_res.append(key)
    # residue tube score: best coupling of any path forced through the residue, relative to optimum
    res_best = {}
    for i, a in enumerate(atoms):
        if i in dS and i in dT:
            key = (a['chain'], a['resnum'], a['resname'])
            val = dS[i] + dT[i] - best
            res_best[key] = min(res_best.get(key, 1e9), val)
    # side-chain knockout (Ala-like truncation) for residues within 100x of optimum
    ko = {}
    if knockout:
        cand = [k for k, v in res_best.items() if v <= math.log(100) and k[2] not in ('HEM', 'HEA', 'HEC', 'CU', 'FES')]
        for key in cand:
            ex = [i for i, a in enumerate(atoms) if (a['chain'], a['resnum'], a['resname']) == key and a['atom'] not in BACKBONE]
            if not ex:
                ko[key] = 0.0; continue
            Hk = H.copy(); Hk.remove_nodes_from(ex)
            try:
                d = nx.dijkstra_path_length(Hk, 'S', 'T')
            except nx.NetworkXNoPath:
                d = float('inf')
            ko[key] = (d - best) / math.log(10)  # log10 drop in |T_DA|
    fe_d = None
    try:
        pa = [atoms[i]['pos'] for i in D if atoms[i]['el'] in METALS]
        pb = [atoms[i]['pos'] for i in A if atoms[i]['el'] in METALS]
        fe_d = min(np.linalg.norm(x - y) for x in pa for y in pb)
    except ValueError:
        pass
    return dict(label=label, pdb=pdb, log10_coupling=-best / math.log(10), metal_dist=fe_d,
                path_res=path_res, steps=steps, res_best=res_best, ko=ko)


STEPS = [
    ('9CG3', 'heme bL -> heme bH (cyt b, monomer 1)', ('J', 401), ('J', 402)),
    ('9CG3', 'heme bL -> heme bL\' (inter-monomer)', ('J', 401), ('V', 401)),
    ('5Z62', 'CuA -> heme a', ('B', 301), ('A', 603)),
    ('5Z62', 'heme a -> heme a3 (control)', ('A', 603), ('A', 604)),
]
FILES = {'9CG3': STRUCTDIR + '/9CG3.cif', '5Z62': STRUCTDIR + '/5Z62.cif'}

if __name__ == '__main__':
    results = []
    for pdb, label, don, acc in STEPS:
        sites = [don, acc]
        if don == ('B', 301): sites.append(('B', 302))
        atoms = load_region(FILES[pdb], sites)
        D = site_atoms(atoms, *don)
        if don == ('B', 301): pass
        r = solve(atoms, don, acc, label, pdb)
        # CuA is a dimer: allow either Cu as donor
        if don == ('B', 301):
            r2 = solve(atoms, ('B', 302), acc, label, pdb)
            if r2['log10_coupling'] > r['log10_coupling']: r = r2
        results.append(r)
        print(f"\n### {label} [{pdb}] metal-metal {r['metal_dist']:.1f} A, best log10|T| ~ {r['log10_coupling']:.2f}")
        print("  path:", ' -> '.join(f"{c}:{n}{s}" for c, s, n in r['path_res']))
        print("  steps:", ' '.join(r['steps']))
        tube = sorted([(v, k) for k, v in r['res_best'].items() if v <= math.log(10)])
        print(f"  residues within 10x of optimum: {len(tube)}")
        top = sorted(r['ko'].items(), key=lambda kv: -kv[1])[:12]
        print("  top side-chain knockout drops (log10):", ', '.join(f"{k[0]}:{k[2]}{k[1]}={v:.2f}" for k, v in top))
    out = []
    for r in results:
        for k, v in r['res_best'].items():
            if v <= math.log(100):
                out.append(dict(step=r['label'], pdb=r['pdb'], chain=k[0], resnum=k[1], resname=k[2],
                                on_best_path=k in r['path_res'],
                                through_path_penalty_log10=v / math.log(10),
                                knockout_drop_log10=r['ko'].get(k)))
    json.dump(out, open(ASSETDIR + '/pathway_residues.json', 'w'), indent=1)
```
