## Supplementary code for "Relaxed evolutionary constraints on mitochondrial electron-transfer-associated variants in human cancers": README.md

# Quantum electron tunnelling in mitochondrial cytochrome b — analysis code

Code for *"Relaxed evolutionary constraints on mitochondrial electron-transfer-associated variants in
human cancers"* (Sung & Cheong).

The pipeline predicts, for every possible missense substitution in mtDNA-encoded cytochrome b
(MT-CYB), its effect on the quantum-mechanical electron-tunnelling rates of the complex III Q-cycle
(Beratan–Onuchic Pathways couplings + Marcus non-adiabatic theory), and tests whether such variants
are under differential selection in the human germline versus tumours (TCGA, PCAWG, MSK-IMPACT).

## 1. Requirements

Python ≥ 3.11 and the packages in `requirements.txt`:

```bash
pip install -r requirements.txt
```

`gemmi` is used for structure parsing, `statsmodels` for the GLM/Cox models, `matplotlib` for figures.

## 2. Data

**Bundled (in `inputs/`)** — no download needed:
- `inputs/structures/9CG3.cif`, `5Z62.cif` — cryo-EM structures of human complex III and IV (RCSB PDB).
  9CG3 (human kidney respiratory complex III, 2.96 A; Zhang & Lyu) is an **unpublished deposition**: cite the
  PDB entry (https://doi.org/10.2210/pdb9cg3/pdb), not a journal article. Chain J models residues **2-379**;
  residues 1 and 380 are unresolved and are therefore absent from every structure-based score.
- `inputs/assets/rCRS.fasta` — revised Cambridge Reference Sequence (NCBI NC_012920).
- `inputs/assets/MT-CYB_residue_classes.tsv` — per-residue structural classification.
- `inputs/assets/pockets.json`, `pathway_residues.json`, `leak_summary.json` — derived geometric/Pathways
  assets (regenerate with `preprocessing/`; see §5).
- `inputs/assets/tcga_purity_aran2015.tsv` — tumour purity (CPE, ABSOLUTE) from Aran et al. 2015.

**External (obtain separately)** — public:
- Somatic/germline mtDNA calls: Gorelick et al., <https://github.com/reznik-lab/mtdna-mutations>
  (`data/original_data/`): `data_mutations_{tcga,pcawg,helix,impact}.txt.gz`, `data_samples_*.txt.gz`,
  and `data/processed_data/` (gene lengths). Point `MTCYB_DATA` at `data/original_data`.
- Overall-survival: TCGA-CDR (Liu et al. 2018), `TCGA-CDR-SupplementalTableS1.txt`, in the same repo's
  `data/ext/`. Point `MTCYB_EXT` at `data/ext`.
- Nuclear-signature analysis only (optional, see §6): GDC ensemble-masked WXS MAFs and the COSMIC v3.4
  GRCh38 exome signatures.

## 3. Configure paths

All paths live in `src/config.py`. Either edit the two defaults there, or set environment variables:

```bash
export MTCYB_DATA=/abs/path/to/reznik-lab/mtdna-mutations/data/original_data
export MTCYB_EXT=/abs/path/to/reznik-lab/mtdna-mutations/data/ext
# optional: export MTCYB_OUT=/abs/path/to/outputs   (default: ./outputs)
```

## 4. Reproduce everything

```bash
bash run_all.sh
```

Outputs (tables `*.tsv`, figures, and `Source_Data_MT-CYB_electron_tunnelling.xlsx`) are written to
`outputs/`. The `run_all.sh` header documents the exact order; scripts can also be run individually
from `src/`.

## 5. Method / script map

| Script (`src/`) | Produces | Notes |
|---|---|---|
| `et_variants.py` | core module: `features()`, `cohorts()`, `rates()`, `marcus()`, `poly_ok()` | Pathways + Marcus; 192-channel strand-aware spectrum null; population-rare site restriction. Its `__main__` fits a **supporting single-step** model (`et_validation.tsv`) **without** the `poly_ok()` restriction — it is not the Fig 3e source |
| `fig2_pipeline.py` | helper module (`load_rcrs`, `tri`, `spectrum`) | imported by the above |
| `qcycle_marcus.py` | multi-step Q-cycle ET score + **Fig 3e** source | `scores()` returns network / bL→bH / proximity; `qcycle_validation.tsv` holds the ET-specific component (`p_diff_only`) |
| `fig1_plot.py` | **Fig 1** | structural + tunnelling framework |
| `fig2_tables.py` | **Fig 2b/2c** source | class O/E + somatic-vs-germline odds ratios (`fig2_out[_poly10]/`) |
| `fig2_charge_split.py` | **Fig 2d** source | O/E by class × charge change |
| `fig2_plot.py` | **Fig 2** | |
| `fig3_tables.py` | **Fig 3a-c** source | ET-score bins, high-perturbation class, ε/λ sensitivity; per-variant table |
| `fig3_plot.py` | **Fig 3** | |
| `fig4_missense_table.py` | Fig 4 prep | per-mutation MT-CYB missense table with ET score/VAF (`fig4_cyb_missense.tsv`) |
| `fig4_tables.py` | **Fig 4a** source | high-perturbation O/E by heteroplasmy stratum |
| `fig4_helix_state.py` | **Fig 4b** source | germline heteroplasmic/homoplasmic split |
| `fig4_purity.py` | **Supp Fig 6** source | purity-adjusted heteroplasmy (CPE/ABSOLUTE) |
| `source_data_extras.py` | **Fig 4c**, **Supp Fig 1** source | per-tumour-type shares; no-filter counterpart of Fig 2 |
| `fig4_plot.py` | **Fig 4** | |
| `robustness_dedup.py` | TCGA/PCAWG patient de-duplication | confirms pooled endpoints are not inflated by overlap |
| `functional_evidence.py` | heteroplasmy vs truncating LoF; complex I co-occurrence (TCGA only) | see note below |
| `cooccurrence_matched.py` | complex I co-occurrence under one design across both cohorts | `ci_matched.tsv` |
| `impact_replication.py` | MSK-IMPACT independent cohort | reported as a non-replication / limitation |
| `multiple_testing_fdr.py` | Benjamini–Hochberg across the primary family | `fdr_primary.tsv` |
| `supp_figures.py` | **Supp Figs S2, S3, S4, S6** | S4 needs signatures (§6) |
| `nuclear_sbs.py` | nuclear SBS exposures | needs GDC MAFs + COSMIC (§6) |
| `build_source_data.py` | `Source_Data_*.xlsx` | assembles all tables above |

Preprocessing (`preprocessing/`, optional — outputs are bundled in `inputs/assets/`):
`detect_pockets.py` (geometric Qo/Qi pocket detection → `pockets.json`) and
`pathways_coupling.py` (Beratan–Onuchic couplings → `pathway_residues.json`, `leak_summary.json`).

Analyses that were run during development but are **not reported** in the manuscript live in
`exploratory/` (see `exploratory/README.md`); they are excluded from `run_all.sh`.

## 6. Signature analysis (optional)

`nuclear_sbs.py` and the S4/S5 panels of `supp_figures.py` require inputs too large to bundle: the GDC
ensemble-masked WXS MAFs (per TCGA cancer type) and the COSMIC v3.4 GRCh38 exome signature matrix.
Set their locations at the top of `nuclear_sbs.py`, run it to produce the SBS exposure/QC tables, then
the signature-dependent panels. The rest of the pipeline does not depend on this step.

## 7. Key parameters (defaults)

Screened Coulomb ΔEm = 14.4·Δq/(ε·r) eV (ε = 8, range 4–16, r floored at 3 Å); T = 310 K.
Marcus per-step reference values (ΔG₀ / λ, eV): Qo→bL −0.20 / 0.80; bL→bH −0.10 / 0.70; bH→Qi −0.02 / 0.70;
CuA→haem a (complex IV) −0.05 / 0.40. λ is scaled by 0.7×, 1× and 1.4× in the sensitivity grid.
Per-step and composite |Δlog₁₀ k| scores are capped at 3 log units. charges D,E = −1, K,R = +1, H = +0.1; high-perturbation = |Δlog₁₀ k| ≥ 0.3 in ≥1 Q-cycle step;
population-rare site filter = HelixMTdb carriers ≤ 10, with somatic expectations computed over the **same**
restricted site universe (omitting this inflates enrichment by ~50%).

## 8. Corrections applied in this version

- `fig2_pipeline.expected_fractions` enumerated sites over `range(CYB_START, CYB_END - 2)` while
  `fig2_tables.annotate` counted observed mutations over the full `CYB_START..CYB_END` span. MT-CYB
  is 1,141 nt (380 codons + a single-nucleotide incomplete stop codon completed by polyadenylation),
  so m.15887 was translated as a non-existent "codon 381" by reading two nucleotides into MT-TT, and
  four somatic calls there — annotated `p.*381=` (Silent) in the source data — were counted as
  missense. Numerator and denominator now share the 380-codon span (nt 14,747–15,886).
  Somatic MT-CYB missense: **528 → 524**; germline sites: **603 → 602**. The structure-based
  analyses (Figs 1, 3, 4) were already on the correct set and are unchanged at 524 / 597, the
  germline difference being the five codon-380 variants that have no coordinates in 9CG3.
- Source Data: `qcycle_validation.tsv` is now included as the Fig 3e sheet; the purity sheet is
  relabelled Supplementary Fig. 6; Fig 4b, Fig 4c and Supplementary Fig. 1 sheets were added.

## 9. License

Code: MIT (`LICENSE`). Bundled third-party data retain their original terms (PDB; NCBI; Aran et al. 2015;
Gorelick et al.; TCGA-CDR).
