## Supplementary code for "Relaxed evolutionary constraints on mitochondrial electron-transfer-associated variants in human cancers": run_all.sh

```
#!/usr/bin/env bash
# Reproduce all tables, figures and the Source Data workbook.
#
# Usage:
#   export MTCYB_DATA=/abs/path/to/reznik-lab/mtdna-mutations/data/original_data
#   export MTCYB_EXT=/abs/path/to/reznik-lab/mtdna-mutations/data/ext
#   bash run_all.sh
#
# Structures (9CG3.cif, 5Z62.cif), rCRS and the derived assets are bundled under inputs/.
# Outputs are written to outputs/ (override with MTCYB_OUT).
set -euo pipefail
cd "$(dirname "$0")/src"

: "${MTCYB_DATA:?set MTCYB_DATA to the original_data directory}"
: "${MTCYB_EXT:?set MTCYB_EXT to the data/ext directory}"
: "${MTCYB_OUT:=$(cd .. && pwd)/outputs}"
export MTCYB_OUT

run () { echo "### $1"; python3 "$1"; }

# --- core selection pipeline (runs from bundled structures + public mutation data) ---
run et_variants.py           # per-step ET features + single-step supporting model (et_validation.tsv)
run fig2_tables.py           # Fig 2b/2c  (writes fig2_out_poly10/)
run fig2_charge_split.py     # Fig 2d     (fig2_charge_split.tsv)
run fig3_tables.py           # Fig 3a-c   (fig3_oe.tsv, fig3_variants.tsv)
run qcycle_marcus.py         # Fig 3e     (qcycle_validation.tsv) -- the ET-specific component
run fig4_missense_table.py   # Fig 4 prep (fig4_cyb_missense.tsv; needs fig3_variants.tsv)
run fig4_tables.py           # Fig 4a     (fig4_vaf_oe.tsv)
run fig4_helix_state.py      # Fig 4b     (fig4_helix_state.tsv)
run fig4_purity.py           # Supp Fig 6 (fig4_purity_oe.tsv)

# --- robustness and supporting analyses ---
run robustness_dedup.py        # TCGA/PCAWG patient de-duplication (dedup_results.tsv)
run functional_evidence.py     # heteroplasmy vs truncating LoF; TCGA-only co-occurrence (func_*.tsv)
run cooccurrence_matched.py    # complex I co-occurrence, one design across both cohorts (ci_matched.tsv)
run impact_replication.py      # MSK-IMPACT independent cohort (impact_repl.tsv)
run multiple_testing_fdr.py    # Benjamini-Hochberg across the primary family (fdr_primary.tsv)
run supp_figures.py            # Supp Figs S2, S3 (S4 needs signatures, see below)

# --- Supplementary Fig. 1: the same analysis without the population-polymorphism filter ---
echo "### Supplementary Fig. 1 (no polymorphism filter)"
POLY_MAX=1000000000 python3 fig2_tables.py > /dev/null
ET_NOFILTER=1 CHARGE_OUT="$MTCYB_OUT/fig2_charge_split_nofilter.tsv" python3 fig2_charge_split.py > /dev/null
run source_data_extras.py      # Fig 4c and Supp Fig 1 source tables
# Supp Fig 1 is Fig 2 re-rendered from the unfiltered tables (same layout, different data):
FIG2_IN="$MTCYB_OUT/fig2_out/" FIG2_SUFFIX=_tmp_nofilter \
  FIG2_CHARGE="$MTCYB_OUT/fig2_charge_split_nofilter.tsv" python3 fig2_plot.py
# fig2_plot.py always writes "Figure2<suffix>"; rename so the file says which figure it is.
for e in png pdf; do mv -f "$MTCYB_OUT/Figure2_tmp_nofilter.$e" "$MTCYB_OUT/SuppFig_S1_nofilter.$e"; done

# --- figures (require matplotlib) ---
run fig1_plot.py
run fig2_plot.py
run fig3_plot.py
run fig4_plot.py

# --- nuclear mutational signatures (OPTIONAL: extra external inputs) ---
# nuclear_sbs.py and the S4/S5 panels of supp_figures.py need the GDC ensemble-masked MAFs and the
# COSMIC v3.4 GRCh38 exome signatures; see README "Signature analysis". They are not required by any
# main-text figure.
#   run nuclear_sbs.py

# --- assemble the Source Data workbook (needs the tables above) ---
run build_source_data.py

echo "Done. See $MTCYB_OUT."
```
