## Supplementary code for "Relaxed evolutionary constraints on mitochondrial electron-transfer-associated variants in human cancers": build_source_data.py

```
"""Assemble the Nature Communications Source Data workbook for the MT-CYB electron-tunnelling study."""
import os as _os, sys as _sys
_sys.path.insert(0, _os.path.dirname(_os.path.abspath(__file__)))
from config import DATADIR, DATAROOT, EXTDIR, STRUCTDIR, ASSETDIR, OUTDIR

import pandas as pd, numpy as np
from openpyxl import Workbook
from openpyxl.styles import Font, PatternFill, Alignment, Border, Side
from openpyxl.utils import get_column_letter

R = OUTDIR + '/'
HEAD_FILL = PatternFill('solid', fgColor='1F3864')
HEAD_FONT = Font(name='Arial', bold=True, color='FFFFFF', size=10)
TITLE_FONT = Font(name='Arial', bold=True, size=13, color='1F3864')
SUB_FONT = Font(name='Arial', italic=True, size=10, color='555555')
CELL_FONT = Font(name='Arial', size=10)
BOLD = Font(name='Arial', size=10, bold=True)
THIN = Side(style='thin', color='BFBFBF')
BORDER = Border(left=THIN, right=THIN, top=THIN, bottom=THIN)

wb = Workbook()

# ---- sheet definitions: (sheet, figure title, caption, source tsv, rename/round) ----
def load(fn, **kw):
    return pd.read_csv(R + fn, sep='\t', **kw)

# Fig2 obs/exp real: order cohorts germline, TCGA, PCAWG
f2 = load('fig2_out_poly10/fig2_obs_exp_real.tsv')
f2or = load('fig2_out_poly10/fig2_somatic_vs_germline.tsv')
f2cs = load('fig2_charge_split.tsv')
f3 = load('fig3_oe.tsv')
f3v = load('fig3_variants.tsv')
f4v = load('fig4_vaf_oe.tsv')
f4p = load('fig4_purity_oe.tsv')
s2 = load('supp_S2_synonymous.tsv')
s3 = load('supp_S3_complexIV.tsv')
rescls = pd.read_csv(ASSETDIR + '/MT-CYB_residue_classes.tsv', sep='\t')  # bundled asset
feat = load('et_features_CIII.tsv')
dedup = load('dedup_results.tsv')
fdr = load('fdr_primary.tsv')
fvaf = load('func_vaf_groups.tsv')
fco = load('func_cooccur.tsv')
etval = load('et_validation.tsv')
impact = load('impact_sourcedata.tsv')
qcyc = load('qcycle_validation.tsv')
helix = load('fig4_helix_state.tsv')
f4ct = load('fig4_cancertype.tsv')
s1nf = load('suppfig1_nofilter.tsv')
cimatch = load('ci_matched.tsv')

# Fig1 source: per-residue structural/ET descriptors
f1 = rescls.rename(columns={
    'aa_pos': 'MT-CYB residue', 'residue': 'amino acid', 'mtDNA_codon': 'mtDNA codon',
    'category': 'structural class', 'qo_lining': 'Qo-pocket lining',
    'bL_Qo_tube10': 'in bL-Qo coupling tube (<10A)', 'bL_Qo_knockout_drop_log10': 'Pathways knockout Dlog10|T_DA|',
    'min_dist_ISP': 'min dist to Rieske ISP (A)', 'min_dist_bL': 'min dist to haem bL (A)',
    'min_dist_bH': 'min dist to haem bH (A)'})

# Fig3 per-variant master
f3v = f3v.rename(columns={'pos': 'mtDNA position', 'alt': 'alt allele', 'aa_pos': 'MT-CYB residue',
                          'aa_ref': 'ref aa', 'aa_alt': 'alt aa', 'dq': 'charge change (dq)',
                          'network': 'Q-cycle ET score |Dlog10 k|', 'prox_eV': 'proximity control (|dEm|, eV)',
                          'tumours': 'tumour occurrences (TCGA+PCAWG)', 'germline': 'germline present (HelixMTdb)'})
f3v['high-ET perturbation (>=0.3)'] = (f3v['Q-cycle ET score |Dlog10 k|'] >= 0.3)

SHEETS = [
    ('Contents', None, None, None),
    ('Fig1_structure', 'Figure 1',
     'Structural and electron-tunnelling descriptors for every MT-CYB residue (cryo-EM 9CG3, monomer J). '
     'Distances are side-chain centroid to redox centre; the Pathways knockout column is the per-residue drop in '
     'log10|T_DA| for the haem bL->bH coupling on side-chain removal.', f1),
    ('Fig2b_class_OE', 'Figure 2b',
     'Observed/expected ratio of MT-CYB missense variants by structural class, per cohort. Spectrum-aware '
     '192-channel strand-aware null; somatic cohorts restricted to population-rare sites (HelixMTdb carriers <=10); '
     'expectation computed over the same restricted site universe. 95% binomial CIs. P, two-sided binomial test.', f2),
    ('Fig2c_oddsratio', 'Figure 2c',
     'Relaxed constraint: odds ratio for each class\'s share of missense variants, pooled somatic (TCGA+PCAWG) '
     'versus germline. P, two-sided Fisher exact test.', f2or),
    ('Fig2d_charge_split', 'Figure 2d',
     'Observed/expected of MT-CYB missense split by structural class and by charge change (|dq| >= 0.9), per cohort. '
     '95% binomial CIs; P, two-sided binomial test.', f2cs),
    ('Fig3b_score_bins', 'Figure 3b,c',
     'Observed/expected across bins of the Q-cycle Marcus ET-perturbation score, and the high-perturbation class '
     '(score >= 0.3), per cohort and pooled. Poisson exact 95% CIs; P, two-sided Poisson test. eps/lam_scale rows '
     'are the parameter-sensitivity grid.', f3),
    ('Fig3_variant_scores', 'Figure 3 (per variant)',
     'All possible MT-CYB missense SNVs with their Q-cycle ET-perturbation score, proximity control, charge change, '
     'and observed germline/tumour occurrences. This is the variant-level source for Fig. 3 and the discriminating-'
     'variant list.', f3v),
    ('Fig3e_ET_specific', 'Figure 3e',
     'THE source for Fig. 3e. Quasi-Poisson GLM with spectrum offsets on the three-step Q-cycle score. '
     'p_diff_only is the ET-specific component: the Q-cycle score orthogonalised to the electrostatic shift '
     '(prox) and the charge change, entered alongside prox and the structural/chemical controls. '
     'r_network_prox documents the collinearity (0.94 at the reference setting). Rows span the eps/lambda grid; '
     'the reference setting is eps=8, lam_scale=1.0.', qcyc),
    ('Fig4a_heteroplasmy', 'Figure 4a',
     'Observed/expected of high-ET-perturbation MT-CYB missense stratified by heteroplasmy (variant allele '
     'fraction), per cohort. trend rows give the Cochran-Armitage-style slope and its P. Poisson 95% CIs.', f4v),
    ('Fig4b_germline_state', 'Figure 4b',
     'Germline MT-CYB missense sites in HelixMTdb split by whether the site is seen only in the heteroplasmic '
     'state or is homoplasmic in at least one carrier, and whether it is ET-perturbing (score >= 0.3). '
     'Two-sided Fisher exact test (odds ratio 1.63, P = 0.19).', helix),
    ('Fig4c_cancertype', 'Figure 4c',
     'Share of somatic MT-CYB missense mutations that are ET-perturbing (score >= 0.3), per tumour type with at '
     'least eight mutations (TCGA and PCAWG project codes). p_vs_pancancer is a two-sided binomial test against '
     'the pan-cancer share; heterogeneity across types was assessed by a chi-squared test '
     '(chi2 = 27.79, d.f. = 19, P = 0.088).', f4ct),
    ('SuppFig1_nofilter', 'Supplementary Fig. 1',
     'As Fig. 2 but without the population-polymorphism filter (all somatic MT-CYB missense calls, n = 785). '
     'Rows are stacked by analysis: class O/E, somatic-versus-germline odds ratios, and the class x charge split. '
     'The haem-environment charge-change result is unchanged (germline O/E = 0.12; pooled somatic O/E = 1.87).', s1nf),
    ('SuppFig2_synonymous', 'Supplementary Fig. 2',
     'Synonymous calibration: observed/expected of synonymous MT-CYB variants by structural class, per cohort. '
     'Same null and site universe as Fig. 2. 95% binomial CIs; P, two-sided binomial test.', s2),
    ('SuppFig3_complexIV', 'Supplementary Fig. 3',
     'Complex III (MT-CYB, haem bL->bH) versus Complex IV (MT-CO1/CO2, CuA->haem a) observed/expected across '
     'ET-perturbation-score bins, germline and pooled tumour. 95% Poisson CIs.', s3),
    ('SuppFig6_purity', 'Supplementary Fig. 6',
     'Robustness of the heteroplasmy threshold to tumour purity (TCGA): raw VAF, VAF corrected by CPE and by '
     'ABSOLUTE, and high-purity tumours (CPE >= 0.7). Poisson 95% CIs; P, two-sided Poisson test. Note the '
     'ABSOLUTE-corrected <0.1 stratum contains only 6 tumours and is underpowered (O/E 1.31, P = 0.65).', f4p),
    ('Stats_dedup', 'Statistics: patient dedup',
     'Robustness of the two headline pooled endpoints to TCGA/PCAWG patient overlap. DEDUP drops PCAWG samples '
     'whose patient barcode is already in TCGA. The two cohorts share 789 patients; 628 of the corresponding '
     'PCAWG samples carry mtDNA calls and were removed (1,975 PCAWG SNV rows).', dedup),
    ('Stats_FDR', 'Statistics: multiple testing',
     'Benjamini-Hochberg FDR across the primary confirmatory test family. All tests remain significant at q<0.05.', fdr),
    ('IMPACT_replication', 'Independent cohort (non-replication)',
     'MSK-IMPACT deep targeted-panel cohort (539 MT-CYB missense). Neither headline endpoint replicated '
     '(haem-env charge-change O/E=1.10, P=0.64; high-ET O/E=0.94, P=0.54). The cohort\'s heteroplasmy '
     'distribution (median VAF 0.5) under-samples the low-VAF range carrying the signal; reported as a '
     'limitation in the Discussion. Poisson tests; blank CI for the VAF-stratified rows.', impact),
    ('Func_heteroplasmy', 'Functional evidence 1',
     'Heteroplasmy (VAF) distribution of TCGA MT-CYB variants by functional group. High-ET missense resemble '
     'truncating loss-of-function (Mann-Whitney P=0.44) and differ from low-ET missense (P=0.0059). This is '
     'consistent with, but does not establish, heteroplasmy-dependent purifying selection.', fvaf),
    ('Func_cooccurrence', 'Functional evidence 2',
     'Complex I (ND genes) truncating co-occurrence in tumours carrying high-ET MT-CYB missense versus others '
     '(TCGA, patient-level). Fisher OR=0.84, P=0.74. Note this test draws the exposure from somatic calls only '
     'but the ND-truncating outcome from all classes; see Func_cooccurrence_matched for a consistently filtered, '
     'cohort-matched analysis. No conclusion about complex III specificity is drawn from this test alone.', fco),
    ('Func_cooccurrence_matched', 'Functional evidence 2 (matched)',
     'Complex I (ND) truncating co-occurrence under one design applicable to both cohorts: sample-level, '
     'Variant_Classification-based truncating, population-rare filter, denominator = tumours with >=1 mtDNA call. '
     'TCGA null (CMH OR 0.56, P = 0.20); PCAWG nominally positive (1.67, P = 0.023) but not after removing samples '
     'shared with TCGA (1.52, P = 0.10); pooled de-duplicated null (1.17, P = 0.48).', cimatch),
    ('Supp_regression', 'Supplementary regression',
     'Quasi-Poisson GLM coefficients for the Marcus ET-perturbation term (marcus): alone, in the full model with '
     'structural/chemical controls, and the proximity control in the full model, across eps and lambda settings and '
     'cohorts. NOTE: this is a supporting single-step (bL->bH) model and is NOT the source for Fig. 3e; the '
     'Fig. 3e statistics come from the three-step Q-cycle model in sheet Fig3e_ET_specific. This table also '
     'omits the population-rare site restriction applied elsewhere, so its somatic rows are not directly '
     'comparable with Figs 2-4.', etval),
]


def round_df(df):
    out = df.copy()
    for c in out.columns:
        if pd.api.types.is_float_dtype(out[c]):
            if out[c].abs().dropna().lt(1e-3).any() or out[c].name.lower().startswith(('p', 'q')) or 'p_' in str(c).lower() or str(c).lower() in ('p', 'q_bh', 'p_value'):
                out[c] = out[c].map(lambda x: f'{x:.3e}' if pd.notnull(x) and (abs(x) < 1e-3 and x != 0) else (round(x, 4) if pd.notnull(x) else x))
            else:
                out[c] = out[c].round(4)
    return out


def write_table(ws, df, start_row):
    df = round_df(df)
    # header
    for j, col in enumerate(df.columns, start=1):
        c = ws.cell(row=start_row, column=j, value=str(col))
        c.fill = HEAD_FILL; c.font = HEAD_FONT; c.border = BORDER
        c.alignment = Alignment(horizontal='center', vertical='center', wrap_text=True)
    for i, (_, row) in enumerate(df.iterrows(), start=start_row + 1):
        for j, col in enumerate(df.columns, start=1):
            v = row[col]
            if isinstance(v, (np.integer,)): v = int(v)
            elif isinstance(v, (np.floating,)): v = float(v)
            elif isinstance(v, (np.bool_, bool)): v = bool(v)
            c = ws.cell(row=i, column=j, value=v)
            c.font = CELL_FONT; c.border = BORDER
            c.alignment = Alignment(horizontal='center' if not isinstance(v, str) or len(str(v)) < 24 else 'left', vertical='center')
    # column widths
    for j, col in enumerate(df.columns, start=1):
        w = max(len(str(col)), *(len(str(x)) for x in df[col].astype(str))) if len(df) else len(str(col))
        ws.column_dimensions[get_column_letter(j)].width = min(max(w + 2, 10), 42)
    ws.freeze_panes = ws.cell(row=start_row + 1, column=1)


# ---- Contents sheet ----
ws = wb.active; ws.title = 'Contents'
ws['A1'] = 'Source Data'; ws['A1'].font = TITLE_FONT
ws['A2'] = ('Relaxed evolutionary constraints on mitochondrial electron-transfer-associated variants '
            'in human cancers')
ws['A2'].font = Font(name='Arial', bold=True, size=11)
ws['A3'] = ('Sung J-Y, Cheong J-H.  Each sheet lists the figure/analysis it supports and the exact test used. '
            'Supplementary Figs 4 and 5 depend on the nuclear mutational-signature step and are not included here.')
ws['A3'].font = SUB_FONT
ws['A5'] = 'Sheet'; ws['B5'] = 'Figure / analysis'; ws['C5'] = 'Description'
for col in ('A5', 'B5', 'C5'):
    ws[col].fill = HEAD_FILL; ws[col].font = HEAD_FONT; ws[col].border = BORDER
    ws[col].alignment = Alignment(horizontal='left', vertical='center')
r = 6
for name, fig, cap, df in SHEETS:
    if name == 'Contents': continue
    ws.cell(row=r, column=1, value=name).font = BOLD
    ws.cell(row=r, column=2, value=fig).font = CELL_FONT
    ws.cell(row=r, column=3, value=cap).font = CELL_FONT
    for cc in (1, 2, 3):
        ws.cell(row=r, column=cc).alignment = Alignment(vertical='top', wrap_text=(cc == 3))
        ws.cell(row=r, column=cc).border = BORDER
    r += 1
ws.column_dimensions['A'].width = 22; ws.column_dimensions['B'].width = 24; ws.column_dimensions['C'].width = 95
ws.freeze_panes = 'A6'

# ---- data sheets ----
for name, fig, cap, df in SHEETS:
    if name == 'Contents': continue
    ws = wb.create_sheet(name)
    ws['A1'] = f'{fig} — Source Data'; ws['A1'].font = TITLE_FONT
    ws['A2'] = cap; ws['A2'].font = SUB_FONT
    ws.merge_cells(start_row=2, start_column=1, end_row=2, end_column=min(len(df.columns), 10))
    ws.row_dimensions[2].height = 42
    ws['A2'].alignment = Alignment(wrap_text=True, vertical='top')
    write_table(ws, df, start_row=4)

out = OUTDIR + '/Source_Data_MT-CYB_electron_tunnelling.xlsx'
wb.save(out)
print('saved', out, '|', len(wb.sheetnames), 'sheets:', wb.sheetnames)
```
