## Supplementary code for "Relaxed evolutionary constraints on mitochondrial electron-transfer-associated variants in human cancers": config.py

```
"""Central path configuration for the MT-CYB electron-tunnelling analysis.

Every script imports the directory constants defined here, so you set paths once. Override any with
an environment variable (shown in brackets) or by editing the defaults below.

Expected layout:
  DATADIR  [MTCYB_DATA]   public mtDNA calls (Gorelick et al., reznik-lab/mtdna-mutations/data/
                          original_data): data_mutations_{tcga,pcawg,helix,impact}.txt.gz, data_samples_*.txt.gz
  DATAROOT               parent of DATADIR (…/data); holds original_data/ and processed_data/
  EXTDIR   [MTCYB_EXT]    external clinical tables (…/data/ext): TCGA-CDR-SupplementalTableS1.txt
  STRUCTDIR[MTCYB_STRUCT] cryo-EM structures (bundled): 9CG3.cif, 5Z62.cif
  ASSETDIR               bundled derived assets: rCRS.fasta, MT-CYB_residue_classes.tsv, pockets.json,
                          pathway_residues.json, leak_summary.json, tcga_purity_aran2015.tsv
  OUTDIR   [MTCYB_OUT]    all generated tables and figures are written here
"""
import os

_ROOT = os.path.dirname(os.path.dirname(os.path.abspath(__file__)))  # repo root


def _dir(env, default, trailing=False):
    p = os.environ.get(env, default)
    if trailing and not p.endswith('/'):
        p += '/'
    return p


# --- EDIT to point at your copy of the public mutation data (or set the env vars) ---
DATADIR = _dir('MTCYB_DATA', '/path/to/reznik-lab/mtdna-mutations/data/original_data', trailing=True)
EXTDIR = _dir('MTCYB_EXT',  '/path/to/reznik-lab/mtdna-mutations/data/ext', trailing=True)

# parent of original_data (…/data/), for scripts that also read processed_data/
DATAROOT = os.path.dirname(DATADIR.rstrip('/')) + '/'

# bundled with the repository
STRUCTDIR = _dir('MTCYB_STRUCT', os.path.join(_ROOT, 'inputs', 'structures'))
ASSETDIR = os.path.join(_ROOT, 'inputs', 'assets')

# outputs
OUTDIR = _dir('MTCYB_OUT', os.path.join(_ROOT, 'outputs'))
os.makedirs(OUTDIR, exist_ok=True)
```
