## Supplementary code for "Relaxed evolutionary constraints on mitochondrial electron-transfer-associated variants in human cancers": cooccurrence_matched.py

```
"""Complex I (ND) truncating co-occurrence under a single design applicable to BOTH cohorts.

The co-occurrence test in functional_evidence.py is TCGA-only: it is patient-level and uses the
'truncating' flag, neither of which exists in the PCAWG table, and it draws the exposure from
somatic calls while taking the ND-truncating outcome from all classes. This script applies one
consistent design to both cohorts:

  unit        tumour sample (Tumor_Sample_Barcode; the only ID present in both cohorts)
  exposure    carries >=1 high-ET MT-CYB missense (Q-cycle score >= 0.3), population-rare filter
  outcome     carries >=1 ND truncating call (Variant_Classification based)
  denominator tumours with >=1 mtDNA SNV call
  tests       unstratified Fisher, and Cochran-Mantel-Haenszel stratified by cancer type

PCAWG samples whose normalised barcode is already a TCGA patient are flagged so the analysis can be
repeated with the cohort overlap removed, as elsewhere in the paper (robustness_dedup.py).

Writes ci_matched.tsv (source data for the revised Results statement).
"""
import os as _os, sys as _sys
_sys.path.insert(0, _os.path.dirname(_os.path.abspath(__file__)))
from config import DATADIR, OUTDIR

import re
import numpy as np, pandas as pd
from scipy import stats
from statsmodels.stats.contingency_tables import StratifiedTable

R = DATADIR
TRUNC = ['Nonsense_Mutation', 'Frame_Shift_Del', 'Frame_Shift_Ins']
ND = {'MT-ND1', 'MT-ND2', 'MT-ND3', 'MT-ND4', 'MT-ND4L', 'MT-ND5', 'MT-ND6'}

V = pd.read_csv(OUTDIR + '/fig3_variants.tsv', sep='\t').set_index(['pos', 'alt'])
hc = pd.read_csv(R + 'data_mutations_helix.txt.gz', sep='\t', low_memory=False)
hc = hc[hc.Variant_Type == 'SNP']
poly = hc.groupby(['Start_Position', 'Tumor_Seq_Allele2']).TOTAL_COUNT.sum().to_dict()


def load(p, som, ctcol, name):
    d = pd.read_csv(R + p, sep='\t', low_memory=False)
    if som: d = d[d['class'] == 'somatic']
    d = d.rename(columns={ctcol: 'ct'}); d['cohort'] = name
    return d


T = load('data_mutations_tcga.txt.gz', True, 'CANCER_TYPE', 'TCGA')
P = load('data_mutations_pcawg.txt.gz', False, 'project', 'PCAWG')
P['ct'] = P.ct.str.split('-').str[0]
D = pd.concat([T, P], ignore_index=True)
D['poly'] = [poly.get((a, b), 0) for a, b in zip(D.Start_Position, D.Tumor_Seq_Allele2)]


def norm(x):
    m = re.match(r'(TCGA-[0-9A-Za-z]+-[0-9A-Za-z]+)', str(x)); return m.group(1) if m else str(x)


tcga_pat = set(norm(x) for x in T.PATIENT_ID.astype(str))
D['is_dup'] = (D.cohort == 'PCAWG') & D.Tumor_Sample_Barcode.map(lambda s: norm(s) in tcga_pat)

snv = D[(D.Variant_Type == 'SNP') & (D.poly <= 10)]
cyb = snv[(snv.Hugo_Symbol == 'MT-CYB') & (snv.Variant_Classification == 'Missense_Mutation')].copy()
cyb['score'] = [V.network.get((p, a), np.nan) for p, a in zip(cyb.Start_Position, cyb.Tumor_Seq_Allele2)]
cyb['high'] = cyb.score >= 0.3
ci = D[D.Hugo_Symbol.isin(ND) & D.Variant_Classification.isin(TRUNC)]


def run(label, mask):
    sub = D[mask]
    allS = set(sub.Tumor_Sample_Barcode)
    hiS = set(cyb.loc[cyb.high, 'Tumor_Sample_Barcode']) & allS
    ciS = set(ci.Tumor_Sample_Barcode) & allS
    a = len(hiS & ciS); b = len(hiS - ciS); c = len(ciS - hiS); d = len(allS - hiS - ciS)
    orf, pf = stats.fisher_exact([[a, b], [c, d]])
    ctmap = sub.drop_duplicates('Tumor_Sample_Barcode').set_index('Tumor_Sample_Barcode').ct
    tabs = []
    for _, grp in ctmap.groupby(ctmap):
        s = set(grp.index)
        aa = len(s & hiS & ciS); bb = len((s & hiS) - ciS)
        cc = len((s & ciS) - hiS); dd = len(s - hiS - ciS)
        if (aa + bb) and (aa + cc): tabs.append(np.array([[aa, bb], [cc, dd]]) + 0.0)
    if tabs:
        st = StratifiedTable(tabs)
        orc, pc, ns = st.oddsratio_pooled, st.test_null_odds().pvalue, len(tabs)
    else:
        orc, pc, ns = np.nan, np.nan, 0
    print('%-22s n=%5d exposed=%4d ND-trunc=%4d both=%3d | Fisher OR %5.2f P %.4g | CMH OR %5.2f P %.4g (strata %d)'
          % (label, len(allS), len(hiS), len(ciS), a, orf, pf, orc, pc, ns))
    return dict(analysis=label, n=len(allS), exposed=len(hiS), nd_truncating=len(ciS), both=a,
                fisher_OR=orf, fisher_P=pf, CMH_OR=orc, CMH_P=pc, n_strata=ns)


print('=== Complex I truncating co-occurrence: matched design, both cohorts ===')
rows = [run('TCGA', D.cohort == 'TCGA'),
        run('PCAWG (all)', D.cohort == 'PCAWG'),
        run('PCAWG (dedup)', (D.cohort == 'PCAWG') & (~D.is_dup)),
        run('Pooled (dedup)', (D.cohort == 'TCGA') | ((D.cohort == 'PCAWG') & (~D.is_dup)))]
pd.DataFrame(rows).to_csv(OUTDIR + '/ci_matched.tsv', sep='\t', index=False)
print('saved', OUTDIR + '/ci_matched.tsv')
```
