## Supplementary code for "Relaxed evolutionary constraints on mitochondrial electron-transfer-associated variants in human cancers": et_variants.py

```
"""Does a Marcus–Pathways ET-perturbation score explain somatic/germline missense patterns beyond
simpler structural and chemical predictors?

Unit of analysis: every possible missense SNV in the gene(s) of an ET step.
Outcome: number of observed variants (somatic: tumours; germline: HelixMTdb site present).
Offset: log of the cohort's own 192-channel strand-aware mutation rate for that SNV, so the
mutational spectrum is absorbed. Quasi-Poisson GLM (Pearson-scaled SE).
Predictors (standardised):
  marcus   |log10 k_mut/k_wt| for the ET step (Marcus driving-force term + Pathways |T_DA| term)
  prox     screened Coulomb magnitude of the charge change at the nearer redox centre (no
           donor/acceptor differential) -- "charge near a cofactor" control
  dcharge  |net charge change|
  dKD      |Kyte–Doolittle hydropathy change|;  dvol |side-chain volume change|
  burial   heavy atoms within 10 A of the side chain;  mind  min distance to either redox centre
Key test: marcus coefficient in the full model, and marcus x somatic interaction in a stacked model.
"""
import os as _os, sys as _sys
_sys.path.insert(0, _os.path.dirname(_os.path.abspath(__file__)))
from config import DATADIR, DATAROOT, EXTDIR, STRUCTDIR, ASSETDIR, OUTDIR

import sys, json, math
import numpy as np, pandas as pd, gemmi
import statsmodels.api as sm
sys.path.insert(0, _os.path.dirname(_os.path.abspath(__file__)))
import fig2_pipeline as fp

R = DATADIR
seq = fp.load_rcrs(ASSETDIR + '/rCRS.fasta')
kT = 8.617e-5 * 310
CHG = {'D': -1, 'E': -1, 'K': 1, 'R': 1, 'H': 0.1}
KD = dict(A=1.8, R=-4.5, N=-3.5, D=-3.5, C=2.5, Q=-3.5, E=-3.5, G=-0.4, H=-3.2, I=4.5, L=3.8, K=-3.9, M=1.9, F=2.8, P=-1.6, S=-0.8, T=-0.7, W=-0.9, Y=-1.3, V=4.2)
VOL = dict(A=88.6, R=173.4, N=114.1, D=111.1, C=108.5, Q=143.8, E=138.4, G=60.1, H=153.2, I=166.7, L=166.7, K=168.6, M=162.9, F=189.9, P=112.7, S=89.0, T=116.1, W=227.8, Y=193.6, V=140.0)

STEPS = {
    'CIII bL->bH': dict(pdb=STRUCTDIR + '/9CG3.cif', genes={'MT-CYB': (14747, 15887, 'J')},
                        donor=('J', 401, 'FE'), acceptor=('J', 402, 'FE'), dG0=-0.10, lam=0.70,
                        ko_step='heme bL -> heme bH (cyt b, monomer 1)'),
    'CIV CuA->heme a': dict(pdb=STRUCTDIR + '/5Z62.cif', genes={'MT-CO1': (5904, 7445, 'A'), 'MT-CO2': (7586, 8269, 'B')},
                            donor=('B', (301, 302), 'CU'), acceptor=('A', 603, 'FE'), dG0=-0.05, lam=0.40,
                            ko_step='CuA -> heme a'),
}
B = 'TCAG'; AAS = 'FFLLSSSSYY**CCWWLLLLPPPPHHQQRRRRIIMMTTTTNNKKSSSSVVVVAAAADDEEGGGG'
CODE = {x + y + z: AAS[i] for i, (x, y, z) in enumerate((x, y, z) for x in B for y in B for z in B)}
CODE.update({'AGA': '*', 'AGG': '*', 'ATA': 'M', 'TGA': 'W'})


def annotate(pos, alt, start):
    k = (pos - start) // 3 + 1; off = (pos - start) % 3; s = start + 3 * (k - 1)
    c = seq[s - 1:s + 2]; mc = c[:off] + alt + c[off + 1:]
    a, b = CODE[c], CODE[mc]
    return k, a, b, ('missense' if a != b and '*' not in (a, b) else 'other')


def centre(m, spec):
    ch, num, name = spec
    nums = num if isinstance(num, tuple) else (num,)
    return np.mean([[a.pos.x, a.pos.y, a.pos.z] for n in nums for a in m[ch][str(n)][0] if a.name == name], axis=0)


def features(step):
    S = STEPS[step]
    st = gemmi.read_structure(S['pdb']); st.remove_hydrogens(); m = st[0]
    D, A = centre(m, S['donor']), centre(m, S['acceptor'])
    allxyz = np.array([[a.pos.x, a.pos.y, a.pos.z] for c in m for r in c for a in r])
    from scipy.spatial import cKDTree
    tree = cKDTree(allxyz)
    ko = {(r['chain'], r['resnum']): r['knockout_drop_log10'] or 0.0
          for r in json.load(open(ASSETDIR + '/pathway_residues.json')) if r['step'] == S['ko_step']}
    rows = []
    for gene, (start, end, ch) in S['genes'].items():
        geo = {}
        for r in m[ch]:
            if r.het_flag == 'H': continue
            sc = [a for a in r if a.name not in ('N', 'CA', 'C', 'O')] or [a for a in r if a.name == 'CA']
            x = np.mean([[a.pos.x, a.pos.y, a.pos.z] for a in sc], axis=0)
            geo[r.seqid.num] = (np.linalg.norm(x - D), np.linalg.norm(x - A), len(tree.query_ball_point(x, 10.0)))
        for pos in range(start, end - 2):
            t = fp.tri(seq, pos)
            for alt in 'ACGT':
                if alt == t[1]: continue
                k, a, b, eff = annotate(pos, alt, start)
                if eff != 'missense' or k not in geo: continue
                rD, rA, bur = geo[k]
                rows.append(dict(step=step, gene=gene, pos=pos, alt=alt, tri=t, aa_pos=k, aa_ref=a, aa_alt=b,
                                 rD=rD, rA=rA, burial=bur, dq=CHG.get(b, 0) - CHG.get(a, 0),
                                 dKD=abs(KD[b] - KD[a]), dvol=abs(VOL[b] - VOL[a]) / 100,
                                 ko=ko.get((ch, k), 0.0)))
    return pd.DataFrame(rows)


def marcus(df, dG0, lam, eps):
    dEmD = 14.4 * df.dq / (eps * df.rD.clip(lower=3)); dEmA = 14.4 * df.dq / (eps * df.rA.clip(lower=3))
    ddG = -(dEmA - dEmD)
    f = lambda g: -((g + lam) ** 2) / (4 * lam * kT) / math.log(10)
    return (f(dG0 + ddG) - f(dG0) - 2 * df.ko).abs().clip(upper=3), ddG


def cohorts():
    hc = pd.read_csv(R + 'data_mutations_helix.txt.gz', sep='\t', low_memory=False); hc = hc[hc.Variant_Type == 'SNP']
    poly = hc.groupby(['Start_Position', 'Tumor_Seq_Allele2']).TOTAL_COUNT.sum().to_dict()
    def load(p, som=False, filt=False):
        d = pd.read_csv(R + p, sep='\t', low_memory=False)
        if som: d = d[d['class'] == 'somatic']
        d = d[d.Variant_Type == 'SNP']
        o = pd.DataFrame(dict(sample=d.Tumor_Sample_Barcode.values, pos=d.Start_Position.astype(int).values, alt=d.Tumor_Seq_Allele2.str.upper().values))
        if filt: o = o[[poly.get((a, b), 0) <= 10 for a, b in zip(o.pos, o.alt)]]
        return o
    H = load('data_mutations_helix.txt.gz'); H['sample'] = 'h' + H.pos.astype(str) + H.alt
    import os
    filt = os.environ.get('ET_NOFILTER') is None
    return {'germline': H, 'TCGA': load('data_mutations_tcga.txt.gz', True, filt), 'PCAWG': load('data_mutations_pcawg.txt.gz', False, filt)}


def rates(d):
    # exclude all protein-coding nonsynonymous calls from rate estimation would need every gene;
    # exclude only the genes analysed here (their missense) -- conservative and simple
    ex = set()
    for S in STEPS.values():
        for gene, (start, end, ch) in S['genes'].items():
            sel = d[(d.pos >= start) & (d.pos <= end)]
            for r in sel.itertuples():
                if annotate(r.pos, r.alt, start)[3] == 'missense': ex.add((r.sample, r.pos, r.alt))
    return fp.spectrum(d, seq, ex)


PRED = ['marcus', 'prox', 'dcharge', 'dKD', 'dvol', 'burial', 'mind']


def design(F, eps):
    X = pd.DataFrame(index=F.index)
    X['marcus'] = F.marcus
    X['prox'] = (14.4 * F.dq.abs() / (eps * np.minimum(F.rD, F.rA).clip(lower=3)))
    X['dcharge'] = F.dq.abs(); X['dKD'] = F.dKD; X['dvol'] = F.dvol
    X['burial'] = F.burial; X['mind'] = np.minimum(F.rD, F.rA)
    return (X - X.mean()) / X.std()


def fit(y, X, off, cols):
    mod = sm.GLM(y, sm.add_constant(X[cols]), family=sm.families.Poisson(), offset=off)
    r = mod.fit(scale='X2')
    return r


if __name__ == '__main__':
    import itertools
    C = cohorts(); RT = {k: rates(v) for k, v in C.items()}
    out = []
    for step, S in STEPS.items():
        F = features(step)
        for eps, lam in [(8, S['lam']), (4, S['lam']), (16, S['lam']), (8, S['lam'] * 0.7), (8, S['lam'] * 1.4)]:
            F['marcus'], F['ddG'] = marcus(F, S['dG0'], lam, eps)
            X = design(F, eps)
            stacked = []
            for name, d in C.items():
                key = d.groupby(['pos', 'alt']).size()
                y = np.array([key.get((p, a), 0) for p, a in zip(F.pos, F.alt)])
                if name == 'germline': y = (y > 0).astype(int)
                off = np.log([RT[name][(t, a)] for t, a in zip(F.tri, F.alt)])
                r0 = fit(y, X, off, ['marcus']); r1 = fit(y, X, off, PRED); rc = fit(y, X, off, [p for p in PRED if p != 'marcus'])
                out.append(dict(step=step, eps=eps, lam=round(lam, 2), cohort=name, n_obs=int(y.sum()),
                                beta_alone=r0.params['marcus'], p_alone=r0.pvalues['marcus'],
                                beta_full=r1.params['marcus'], p_full=r1.pvalues['marcus'],
                                beta_prox_full=r1.params['prox'], p_prox_full=r1.pvalues['prox'],
                                dev_gain=rc.deviance - r1.deviance))
                stacked.append(pd.DataFrame(dict(y=y, off=off, som=int(name != 'germline'), **{c: X[c].values for c in PRED})))
            # stacked interaction model: all predictors x somatic; test marcus:som
            Sd = pd.concat(stacked, ignore_index=True)
            Z = Sd[PRED].copy(); Z['som'] = Sd.som
            for c in PRED: Z[c + ':som'] = Sd[c] * Sd.som
            r = sm.GLM(Sd.y, sm.add_constant(Z), family=sm.families.Poisson(), offset=Sd.off).fit(scale='X2')
            out.append(dict(step=step, eps=eps, lam=round(lam, 2), cohort='somatic vs germline (interaction)', n_obs=int(Sd.y.sum()),
                            beta_full=r.params['marcus:som'], p_full=r.pvalues['marcus:som'],
                            beta_prox_full=r.params['prox:som'], p_prox_full=r.pvalues['prox:som']))
        F.to_csv(f"{OUTDIR}/et_features_{step.split()[0]}.tsv", sep='\t', index=False)
    res = pd.DataFrame(out); res.to_csv(OUTDIR + '/et_validation.tsv', sep='\t', index=False)
    pd.set_option('display.width', 250)
    print(res.round(4).to_string(index=False))


def poly_ok(F, max_carriers=10):
    """True for (pos, alt) seen in <= max_carriers HelixMTdb individuals. When somatic calls are filtered on
    population frequency, the expectation must be restricted to the same sites (avoids circularity)."""
    import os
    if os.environ.get('ET_NOFILTER') is not None: return np.ones(len(F), bool)
    hc = pd.read_csv(R + 'data_mutations_helix.txt.gz', sep='\t', low_memory=False); hc = hc[hc.Variant_Type == 'SNP']
    poly = hc.groupby(['Start_Position', 'Tumor_Seq_Allele2']).TOTAL_COUNT.sum().to_dict()
    return np.array([poly.get((p, a), 0) <= max_carriers for p, a in zip(F.pos, F.alt)])
```
