## Supplementary code for "Relaxed evolutionary constraints on mitochondrial electron-transfer-associated variants in human cancers": fig1_plot.py

```
import os as _os, sys as _sys
_sys.path.insert(0, _os.path.dirname(_os.path.abspath(__file__)))
from config import DATADIR, DATAROOT, EXTDIR, STRUCTDIR, ASSETDIR, OUTDIR
import json, csv
import numpy as np, gemmi
import matplotlib
matplotlib.use('Agg')
import matplotlib.pyplot as plt
from matplotlib.patches import FancyBboxPatch, FancyArrowPatch, Rectangle

plt.rcParams.update({'font.family': 'DejaVu Sans', 'font.size': 8, 'axes.linewidth': 0.6,
                     'axes.edgecolor': '#000000', 'xtick.color': '#000000', 'ytick.color': '#000000',
                     'axes.labelcolor': '#000000', 'text.color': '#000000'})
INK, INK2, MUTED, GRID = '#000000', '#000000', '#9a9893', '#e4e3df'
C_QO, C_ISP, C_QI = '#2a78d6', '#eb6834', '#1baf7a'
C_HEME, C_OTHER = '#8a8984', '#d6d5d0'
CAT = {'Qo site / bL leak path': C_QO, 'ISP interface': C_ISP, 'Qi site': C_QI,
       'heme b environment': C_HEME, 'other': C_OTHER}

rows = list(csv.DictReader(open(ASSETDIR + '/MT-CYB_residue_classes.tsv'), delimiter='\t'))
cat = {int(r['aa_pos']): r['category'] for r in rows}
pk = json.load(open(ASSETDIR + '/pockets.json'))
leak = json.load(open(ASSETDIR + '/leak_summary.json'))['leak']
st = gemmi.read_structure(STRUCTDIR + '/9CG3.cif'); st.remove_hydrogens(); m = st[0]
fe = lambda ch, n: np.array([a.pos.tolist() for a in m[ch][str(n)][0] if a.name == 'FE'][0])
ring = lambda ch, n: np.array([a.pos.tolist() for a in m[ch][str(n)][0] if a.name[0] in 'NC' and len(a.name) <= 3 and not a.name.startswith(('CG', 'CB', 'CAA', 'CAD', 'CBA', 'CBD', 'CMA', 'CMB', 'CMC', 'CMD'))])
bL, bH = fe('J', 401), fe('J', 402)
fesP = np.array([a.pos.tolist() for a in m['P']['301'][0] if a.element.name == 'Fe']).mean(0)
edge = lambda A, B: float(np.min(np.linalg.norm(A[:, None] - B[None], axis=2)))
e_blbh = edge(ring('J', 401), ring('J', 402)); e_blbl = edge(ring('J', 401), ring('V', 401))

fig = plt.figure(figsize=(7.2, 7.6), dpi=300)
gs = fig.add_gridspec(3, 2, height_ratios=[0.42, 1.85, 0.8], width_ratios=[1.15, 1], hspace=0.42, wspace=0.28,
                      left=0.1, right=0.98, top=0.97, bottom=0.06)
gr = gs[1, 1].subgridspec(2, 1, height_ratios=[1, 1.05], hspace=0.62)
lab = lambda ax, s, x=-0.04, y=1.02: ax.text(x, y, s, transform=ax.transAxes, fontsize=11, fontweight='bold', va='bottom', ha='right')

# ---------- (a) hypothesis schematic ----------
ax = fig.add_subplot(gs[0, :]); ax.set_xlim(0, 100); ax.set_ylim(0, 22); ax.axis('off'); lab(ax, 'a', 0.0)
boxes = [
    (1, 'MT-CYB missense\nnear a haem\n(charge change $\\Delta q$)'),
    (21, 'Haem redox potential\nshift $\\Delta E_m$\n(protein electrostatics)'),
    (41, 'Driving force $\\Delta G$\nof b$_L$ → b$_H$\nelectron transfer'),
    (61, 'Tunnelling rate\n(Marcus theory)\n$k \\propto |T_{DA}|^2 e^{-(\\Delta G+\\lambda)^2/4\\lambda k_BT}$'),
    (81, 'Selection on\nET-perturbing variants\ngermline vs tumour'),
]
for x, t in boxes:
    ax.add_patch(FancyBboxPatch((x, 4), 17, 14, boxstyle='round,pad=0.4,rounding_size=1.2', fc='white', ec=INK2, lw=0.9))
    ax.text(x + 8.5, 11, t, ha='center', va='center', fontsize=6.6 if x != 61 else 6.3, linespacing=1.3)
for x in (18.6, 38.6, 58.6, 78.6):
    ax.add_patch(FancyArrowPatch((x, 11), (x + 2.6, 11), arrowstyle='-|>', mutation_scale=8, color=INK2, lw=0.8))

# ---------- (b) structure projection ----------
ax = fig.add_subplot(gs[1, 0]); lab(ax, 'b', 0.0)
nrm = (bL - bH) / np.linalg.norm(bL - bH)      # membrane normal, positive toward IMS (P side)
ca = {r.seqid.num: np.array(r['CA'][0].pos.tolist()) for r in m['J'] if r.seqid.num < 400 and r.find_atom('CA', '*')}
X = np.array(list(ca.values())); cen = X.mean(0)
Xp = X - cen; Xp -= np.outer(Xp @ nrm, nrm)
u = np.linalg.svd(Xp, full_matrices=False)[2][0]; u -= (u @ nrm) * nrm; u /= np.linalg.norm(u)
proj = lambda p: np.array([(np.atleast_2d(p) - cen) @ u, (np.atleast_2d(p) - cen) @ nrm]).T
nums = list(ca.keys()); P2 = proj(X)
ax.plot(P2[:, 0], P2[:, 1], color=GRID, lw=0.6, zorder=1)
order = ['other', 'heme b environment', 'Qi site', 'ISP interface', 'Qo site / bL leak path']
for c in order:
    idx = [i for i, n in enumerate(nums) if cat.get(n) == c]
    ax.scatter(P2[idx, 0], P2[idx, 1], s=9 if c != 'other' else 5, color=CAT[c], edgecolors='white', linewidths=0.4, zorder=3)
for site, col in [('Qo', C_QO), ('Qi', C_QI)]:
    q = proj(np.array(pk[site]['points']))
    ax.scatter(q[:, 0], q[:, 1], s=16, color=col, alpha=0.18, lw=0, zorder=2)
    off = (-2, 9) if site == 'Qo' else (8, -2)
    ax.text(q[:, 0].mean() + off[0], q[:, 1].mean() + off[1], f'{site} pocket', color=INK, fontsize=7, va='center', ha='center' if site == 'Qo' else 'left')
for p, t in [(bL, 'heme b$_L$'), (bH, 'heme b$_H$')]:
    q = proj(p)[0]; ax.scatter(*q, marker='D', s=42, color='#7a1f1f', edgecolors='white', lw=0.8, zorder=5)
    ax.text(q[0] + (2.6 if 'L' in t else -2.6), q[1] + (-1.5 if 'L' in t else 0), t, ha='left' if 'L' in t else 'right', va='center', fontsize=7)
q = proj(fesP)[0]; ax.scatter(*q, marker='s', s=30, color='#b8860b', edgecolors='white', lw=0.8, zorder=5)
ax.text(q[0] + 2, q[1] + 3.0, 'Rieske [2Fe-2S]\n(c$_1$ position)', fontsize=6.5, color=INK2, ha='left', va='bottom', bbox=dict(fc='white', ec='none', alpha=0.85, pad=0.5))
a1, a2 = proj(bL)[0], proj(bH)[0]
qo = proj(np.array(pk['Qo']['points'])).mean(0); qi = proj(np.array(pk['Qi']['points'])).mean(0)
for p0, p1, sa, sb in [(qo, a1, 3, 5), (a1, a2, 5, 5), (a2, qi, 5, 3)]:
    ax.add_patch(FancyArrowPatch(p0, p1, arrowstyle='-|>', mutation_scale=8, color=INK, lw=1.0, zorder=4, shrinkA=sa, shrinkB=sb))
ax.set_xlabel('in-membrane axis (Å)'); ax.set_ylabel('membrane normal (Å)\n← matrix (N)      IMS (P) →')
ax.set_aspect('equal', adjustable='datalim'); ax.tick_params(length=2)
for s in ('top', 'right'): ax.spines[s].set_visible(False)
ax.set_title('Cytochrome b (MT-CYB), human CIII 9CG3', fontsize=8, loc='left', color=INK2)

# ---------- (c) tunneling couplings ----------
ax = fig.add_subplot(gr[0]); lab(ax, 'c', -0.02)
data = [
    ('Qo pocket ↔ haem b$_L$', leak['bL -> Qo pocket']['log10T'], leak['bL -> Qo pocket']['edge'], 'quinol-oxidation site'),
    ('haem b$_L$ → b$_H$', -5.72, e_blbh, 'transmembrane step'),
    ('haem b$_H$ → Qi pocket', leak['bH -> Qi pocket']['log10T'], leak['bH -> Qi pocket']['edge'], 'quinone-reduction site'),
    ('haem b$_L$ → b$_L$′', -6.65, e_blbl, 'inter-monomer'),
]
y = np.arange(len(data))[::-1]
for yi, (name, v, e, note) in zip(y, data):
    ax.barh(yi, -v, height=0.34, color='#8a8984', edgecolor='none')
    ax.text(-v + 0.12, yi, f'{v:.1f}', va='center', fontsize=6.6, color=INK)
    ax.text(0.05, yi + 0.24, f'{name}  ·  {e:.1f} Å', fontsize=6.6, va='bottom', color=INK)
    ax.text(7.85, yi + 0.24, note, fontsize=5.9, va='bottom', ha='right', color=INK2)
ax.set_yticks([]); ax.set_xlim(0, 7.9); ax.set_ylim(-0.4, 3.75)
ax.set_xlabel('−log$_{10}$ |T$_{DA}$| (Pathways)', fontsize=7)
ax.xaxis.grid(True, color=GRID, lw=0.5); ax.set_axisbelow(True)
for s_ in ('top', 'right', 'left'): ax.spines[s_].set_visible(False)
ax.tick_params(length=2, labelsize=6.5)
ax.set_title('Electron tunnelling coupling', fontsize=8, loc='left', color=INK2)

# ---------- (d) Marcus framework ----------
ax = fig.add_subplot(gr[1]); lab(ax, 'd', -0.02)
kT = 8.617e-5 * 310; lam = 0.70; g0 = -0.10
fcm = lambda g: -((g + lam) ** 2) / (4 * lam * kT) / np.log(10)
g = np.linspace(-1.05, 0.35, 300)
ax.plot(g, fcm(g) - fcm(g0), color=INK2, lw=1.2)
ax.axvline(-lam, color=MUTED, lw=0.5, ls=':'); ax.text(-lam, 2.45, '−λ', ha='center', fontsize=6.3, color=INK2)
pts = [('WT', g0, INK), ('G48E', g0 - 0.285, '#2a78d6'), ('G116D', g0 + 0.253, '#eb6834')]
for name, gg, c in pts:
    yy = fcm(gg) - fcm(g0)
    ax.scatter(gg, yy, s=26, color=c, edgecolors='white', lw=0.7, zorder=3)
    dx, dy, ha, va = {'WT': (0.04, 0.2, 'left', 'bottom'), 'G48E': (0.04, 0.25, 'left', 'bottom'), 'G116D': (-0.06, -0.05, 'right', 'top')}[name]
    ax.text(gg + dx, yy + dy, f'{name} ({yy:+.1f})' if name != 'WT' else 'WT', fontsize=6.2, ha=ha, va=va, color=INK)
ax.annotate('', (g0 + 0.253, -3.35), (g0, -3.35), arrowprops=dict(arrowstyle='->', color='#eb6834', lw=0.8))
ax.annotate('', (g0 - 0.285, -3.35), (g0, -3.35), arrowprops=dict(arrowstyle='->', color='#2a78d6', lw=0.8))
ax.text(g0, -3.6, '$\\Delta\\Delta G$ from $\\Delta E_m$', ha='center', va='top', fontsize=6, color=INK2)
ax.set_xlim(-1.05, 0.35); ax.set_ylim(-4.3, 2.9)
ax.set_xlabel('$\\Delta G$ of b$_L$ → b$_H$ (eV)', fontsize=7); ax.set_ylabel('log$_{10}$ k / k$_{WT}$', fontsize=7)
ax.tick_params(length=2, labelsize=6.5)
for s_ in ('top', 'right'): ax.spines[s_].set_visible(False)
ax.set_title('Marcus tunnelling rate', fontsize=8, loc='left', color=INK2)

# ---------- (e) residue track ----------
ax = fig.add_subplot(gs[2, :]); lab(ax, 'e', 0.0, 1.08)
for r in rows:
    n = int(r['aa_pos']); c = r['category']
    ax.add_patch(Rectangle((n - 0.5, 0.0 if c == 'other' else -0.25), 1, 0.5 if c == 'other' else 1.0,
                           fc=CAT[c], ec='none'))
ax.set_xlim(0, 381); ax.set_ylim(-1.7, 2.6); ax.set_yticks([])
ax.set_xlabel('MT-CYB amino-acid position')
for s in ('top', 'right', 'left'): ax.spines[s].set_visible(False)
ax.tick_params(length=2)
top = ax.secondary_xaxis('top', functions=(lambda a: 14747 + 3 * (a - 1), lambda p: (p - 14747) / 3 + 1))
top.set_xlabel('mtDNA position (rCRS, m.)', fontsize=7, color=INK2); top.tick_params(labelsize=6.5, length=2)
marks = {83: 'H83', 97: 'H97', 131: 'Y131', 182: 'H182', 196: 'H196', 201: 'H201', 228: 'D228', 271: 'E271\n(PEWY)', 35: 'S35'}
for n, t in marks.items():
    hy = 1.75 if n in (196, 97) else 1.25
    ax.annotate(t, (n, 0.78), (n, hy), ha='center', fontsize=6, color=INK, arrowprops=dict(arrowstyle='-', color=MUTED, lw=0.5))
from collections import Counter
cnt = Counter(r['category'] for r in rows)
for xs, c in zip([0, 64, 145, 210, 305], ['Qo site / bL leak path', 'ISP interface', 'Qi site', 'heme b environment', 'other']):
    ax.add_patch(Rectangle((xs, -1.5), 6, 0.45, fc=CAT[c], ec='none'))
    ax.text(xs + 8, -1.27, f"{'Qo site' if c.startswith('Qo') else c} (n={cnt[c]})", va='center', fontsize=6.8, color=INK)
fig.savefig(OUTDIR + '/Figure1.png'); fig.savefig(OUTDIR + '/Figure1.pdf')
print('edges', round(e_blbh, 1), round(e_blbl, 1))
```
