## Supplementary code for "Relaxed evolutionary constraints on mitochondrial electron-transfer-associated variants in human cancers": fig2_charge_split.py

```
"""Spectrum-aware O/E of MT-CYB missense split by structural class x charge change (|dq| >= 0.9)."""
import os as _os, sys as _sys
_sys.path.insert(0, _os.path.dirname(_os.path.abspath(__file__)))
from config import DATADIR, DATAROOT, EXTDIR, STRUCTDIR, ASSETDIR, OUTDIR

import sys; sys.path.insert(0, _os.path.dirname(_os.path.abspath(__file__)))
import numpy as np, pandas as pd
from scipy import stats
import et_variants as ev
F = ev.features('CIII bL->bH')
cls = pd.read_csv(ASSETDIR + '/MT-CYB_residue_classes.tsv', sep='\t'); cm = dict(zip(cls.aa_pos, cls.category))
grp = {'Qo site / bL leak path': 'Qo/ISP', 'ISP interface': 'Qo/ISP', 'Qi site': 'Qi', 'heme b environment': 'haem', 'other': 'other'}
F['cls'] = F.aa_pos.map(cm).fillna('other').map(grp)
F['chg'] = np.where(F.dq.abs() >= 0.9, 'charge', 'neutral')
F['cell'] = F.cls + ' · ' + F.chg
C = ev.cohorts(); out = []
for name, d in C.items():
    rate = ev.rates(d)
    w = np.array([rate[(t, a)] for t, a in zip(F.tri, F.alt)]) * (1 if name == 'germline' else ev.poly_ok(F))
    key = d.groupby(['pos', 'alt']).size()
    y = np.array([key.get((p, a), 0) for p, a in zip(F.pos, F.alt)])
    if name == 'germline': y = (y > 0).astype(int)
    N = y.sum()
    for cell in sorted(F.cell.unique()):
        m = (F.cell == cell).values; f = w[m].sum() / w.sum(); n = int(y[m].sum())
        bt = stats.binomtest(n, N, f); ci = bt.proportion_ci()
        out.append(dict(cohort=name, cell=cell, observed=n, expected=N * f, oe=n / (N * f), lo=ci[0] / f, hi=ci[1] / f, p=bt.pvalue))
r = pd.DataFrame(out); import os
r.to_csv(os.environ.get('CHARGE_OUT', OUTDIR + '/fig2_charge_split.tsv'), sep='\t', index=False)
pd.set_option('display.width', 200); print(r.round(3).to_string(index=False))
# somatic (TCGA+PCAWG pooled) vs germline contrast within haem: charge vs neutral
for cl in ['haem', 'Qi', 'Qo/ISP', 'other']:
    s = r[(r.cohort != 'germline') & r.cell.str.startswith(cl + ' ·')].groupby('cell')[['observed', 'expected']].sum()
    g = r[(r.cohort == 'germline') & r.cell.str.startswith(cl + ' ·')].set_index('cell')[['observed', 'expected']]
    print(cl, 'somatic O/E', (s.observed / s.expected).round(2).to_dict(), '| germline O/E', (g.observed / g.expected).round(2).to_dict())
```
