## Supplementary code for "Relaxed evolutionary constraints on mitochondrial electron-transfer-associated variants in human cancers": fig2_pipeline.py

```
"""Figure 2 pipeline: somatic MT-CYB missense mutations by structural class and
nuclear oxidative mutational signatures.

Inputs
  --mut      somatic mtDNA SNVs (tsv/csv). Column names set with --mut-cols
             sample,pos,ref,alt,vaf[,cancer_type]   (pos = rCRS 1-based)
  --sbs      per-sample SBS signature activities, wide format (PCAWG SigProfiler style:
             one row per sample, one column per signature, counts). --sbs-sample-col names the id column,
             --sbs-type-col the cancer-type column (optional if cancer_type is in --mut).
  --hole     optional per-sample nuclear hole-hotspot burden (tsv: sample, value)
  --classes  MT-CYB_residue_classes.tsv (Figure 1d)
  --rcrs     rCRS fasta
Outputs (in --out): Figure2.png/.pdf, fig2_mtcyb_missense.tsv, fig2_obs_exp.tsv,
                    fig2_associations.tsv, fig2_log.txt
"""
import os as _os, sys as _sys
_sys.path.insert(0, _os.path.dirname(_os.path.abspath(__file__)))
from config import DATADIR, DATAROOT, EXTDIR, STRUCTDIR, ASSETDIR, OUTDIR

import argparse, os, sys, math, json
import numpy as np, pandas as pd
from scipy import stats

CYB_START, CYB_END = 14747, 15887
CLASSES = ['Qo site / bL leak path', 'ISP interface', 'Qi site', 'heme b environment', 'other']
COL = {'Qo site / bL leak path': '#2a78d6', 'ISP interface': '#eb6834', 'Qi site': '#1baf7a',
       'heme b environment': '#8a8984', 'other': '#d6d5d0'}
LEAK = {'Qo site / bL leak path', 'ISP interface'}
B = 'TCAG'; AA = 'FFLLSSSSYY**CCWWLLLLPPPPHHQQRRRRIIMMTTTTNNKKSSSSVVVVAAAADDEEGGGG'
CODE = {x + y + z: AA[i] for i, (x, y, z) in enumerate((x, y, z) for x in B for y in B for z in B)}
CODE.update({'AGA': '*', 'AGG': '*', 'ATA': 'M', 'TGA': 'W'})   # vertebrate mitochondrial code


def read_table(p):
    return pd.read_csv(p, sep=None, engine='python')


def load_rcrs(p):
    s = ''.join(l.strip() for l in open(p) if not l.startswith('>')).upper()
    assert len(s) == 16569, f'rCRS length {len(s)}'
    return s


def tri(seq, pos):  # circular trinucleotide on reference (L) strand, pos 1-based
    n = len(seq)
    return seq[(pos - 2) % n] + seq[pos - 1] + seq[pos % n]


def annotate_cyb(pos, alt, seq):
    k = (pos - CYB_START) // 3 + 1
    off = (pos - CYB_START) % 3
    s = CYB_START + 3 * (k - 1)
    codon = seq[s - 1:s + 2]
    mcod = codon[:off] + alt + codon[off + 1:]
    a, b = CODE[codon], CODE[mcod]
    eff = 'synonymous' if a == b else ('nonsense' if b == '*' else ('stop_lost' if a == '*' else 'missense'))
    return k, a, b, eff


def spectrum(snv, seq, exclude):
    """192-channel strand-aware rate: count(context,alt)/occurrences(context) on rCRS."""
    occ = {}
    for p in range(1, len(seq) + 1):
        occ[tri(seq, p)] = occ.get(tri(seq, p), 0) + 1
    cnt = {}
    for r in snv.itertuples():
        if (r.sample, r.pos, r.alt) in exclude: continue
        key = (tri(seq, r.pos), r.alt)
        cnt[key] = cnt.get(key, 0) + 1
    rate = {}
    for t, o in occ.items():
        for alt in 'ACGT':
            if alt != t[1]:
                rate[(t, alt)] = (cnt.get((t, alt), 0) + 0.5) / o
    return rate


def expected_fractions(seq, rate, cls, skip=None):
    w = {c: 0.0 for c in CLASSES}
    for pos in range(CYB_START, CYB_END):   # all 380 codons; CYB_END (15887) is the incomplete stop base
        t = tri(seq, pos)
        for alt in 'ACGT':
            if alt == t[1]: continue
            k, a, b, eff = annotate_cyb(pos, alt, seq)
            if eff == 'missense' and not (skip and (pos, alt) in skip):
                w[cls.get(k, 'other')] += rate[(t, alt)]
    tot = sum(w.values())
    return {c: v / tot for c, v in w.items()}


def strat_perm(y, g, strata, n=5000, rng=None):
    """Mean difference of within-stratum-centred y between g==1 and g==0; labels permuted within strata."""
    rng = rng or np.random.default_rng(0)
    y = np.asarray(y, float); g = np.asarray(g, int); strata = np.asarray(strata)
    yc = y.copy()
    for s in np.unique(strata):
        m = strata == s; yc[m] -= y[m].mean()
    obs = yc[g == 1].mean() - yc[g == 0].mean()
    idx = [np.where(strata == s)[0] for s in np.unique(strata)]
    null = np.empty(n)
    for i in range(n):
        gp = g.copy()
        for ix in idx: gp[ix] = rng.permutation(g[ix])
        null[i] = yc[gp == 1].mean() - yc[gp == 0].mean()
    p = (1 + np.sum(np.abs(null) >= abs(obs))) / (n + 1)
    # stratified bootstrap CI
    bs = np.empty(2000)
    for i in range(2000):
        sel = np.concatenate([rng.choice(ix, len(ix)) for ix in idx])
        yy, gg = yc[sel], g[sel]
        bs[i] = (yy[gg == 1].mean() - yy[gg == 0].mean()) if gg.sum() and (gg == 0).sum() else np.nan
    lo, hi = np.nanpercentile(bs, [2.5, 97.5])
    return obs, lo, hi, p


def main(a):
    os.makedirs(a.out, exist_ok=True)
    log = open(os.path.join(a.out, 'fig2_log.txt'), 'w')
    say = lambda *x: (print(*x), print(*x, file=log))
    seq = load_rcrs(a.rcrs)
    cls_tab = pd.read_csv(a.classes, sep='\t')
    cls = dict(zip(cls_tab.aa_pos, cls_tab.category))

    # ---------- mutations ----------
    m = read_table(a.mut)
    names = a.mut_cols.split(',')
    std = ['sample', 'pos', 'ref', 'alt', 'vaf', 'cancer_type'][:len(names)]
    m = m.rename(columns=dict(zip(names, std)))[std]
    m['ref'] = m.ref.astype(str).str.upper(); m['alt'] = m.alt.astype(str).str.upper()
    snv = m[(m.ref.str.len() == 1) & (m.alt.str.len() == 1) & m.ref.isin(list('ACGT')) & m.alt.isin(list('ACGT'))].copy()
    snv['pos'] = snv.pos.astype(int)
    mism = (snv.ref != snv.pos.map(lambda p: seq[p - 1])).sum()
    say(f'SNVs: {len(snv)} in {snv["sample"].nunique()} samples; rCRS ref mismatches: {mism}')
    snv = snv[snv.ref == snv.pos.map(lambda p: seq[p - 1])]

    cyb = snv[(snv.pos >= CYB_START) & (snv.pos <= CYB_END)].copy()
    ann = cyb.apply(lambda r: annotate_cyb(r.pos, r.alt, seq), axis=1, result_type='expand')
    cyb[['aa_pos', 'aa_ref', 'aa_alt', 'effect']] = ann
    cyb['category'] = cyb.aa_pos.map(cls).fillna('other')
    mis = cyb[cyb.effect == 'missense'].copy()
    mis.to_csv(os.path.join(a.out, 'fig2_mtcyb_missense.tsv'), sep='\t', index=False)
    say(f'MT-CYB SNVs: {len(cyb)}; missense {len(mis)}; synonymous {(cyb.effect=="synonymous").sum()}; nonsense {(cyb.effect=="nonsense").sum()}')

    # ---------- observed / expected by class ----------
    excl = set(zip(mis['sample'], mis.pos, mis.alt))
    rate = spectrum(snv, seq, excl)
    f = expected_fractions(seq, rate, cls)
    N = len(mis); rows = []
    for c in CLASSES:
        n = int((mis.category == c).sum()); e = N * f[c]
        ci = stats.binomtest(n, N, f[c]).proportion_ci() if N else (np.nan, np.nan)
        p = stats.binomtest(n, N, f[c]).pvalue if N else np.nan
        rows.append(dict(category=c, observed=n, expected=round(e, 2), oe=n / e if e else np.nan,
                         oe_lo=ci[0] / f[c], oe_hi=ci[1] / f[c], p=p))
    oe = pd.DataFrame(rows); oe.to_csv(os.path.join(a.out, 'fig2_obs_exp.tsv'), sep='\t', index=False)
    say(oe.round(3).to_string(index=False))
    leak_n = int(mis.category.isin(LEAK).sum()); leak_f = sum(f[c] for c in LEAK)
    say(f'Qo+ISP pooled: obs {leak_n}, exp {N*leak_f:.1f}, O/E {leak_n/(N*leak_f) if N else float("nan"):.2f}, '
        f'p={stats.binomtest(leak_n, N, leak_f).pvalue if N else float("nan"):.3g}')

    # ---------- tumour groups & signatures ----------
    sbs = read_table(a.sbs)
    sbs = sbs.rename(columns={a.sbs_sample_col: 'sample'})
    if a.sbs_type_col and a.sbs_type_col in sbs: sbs = sbs.rename(columns={a.sbs_type_col: 'cancer_type_sbs'})
    sig_cols = [c for c in sbs.columns if c.upper().startswith('SBS')]
    sbs['total'] = sbs[sig_cols].sum(1)
    if 'cancer_type_sbs' in sbs: sbs['stratum'] = sbs['cancer_type_sbs']
    elif 'cancer_type' in m: sbs['stratum'] = sbs['sample'].map(m.drop_duplicates('sample').set_index('sample').cancer_type)
    else: sbs['stratum'] = 'all'
    sbs = sbs[sbs.total > 0].dropna(subset=['stratum'])
    hit = mis[mis.vaf >= a.vaf_min]
    leak_s = set(hit[hit.category.isin(LEAK)]['sample'])
    other_s = set(hit['sample']) - leak_s
    sbs['group'] = np.where(sbs['sample'].isin(leak_s), 'Qo/ISP missense',
                            np.where(sbs['sample'].isin(other_s), 'other MT-CYB missense', 'no MT-CYB missense'))
    say(f"tumours matched to signatures: {len(sbs)}; groups: {sbs.group.value_counts().to_dict()}")
    if a.hole:
        h = read_table(a.hole); h.columns = ['sample', 'hole_burden']
        sbs = sbs.merge(h, on='sample', how='left')

    outcomes = [s for s in a.signatures.split(',') if s in sbs] + (['hole_burden'] if a.hole else [])
    res = []
    rng = np.random.default_rng(1)
    for comp, gsel in [('Qo/ISP vs no MT-CYB', ['Qo/ISP missense', 'no MT-CYB missense']),
                       ('other MT-CYB vs no MT-CYB', ['other MT-CYB missense', 'no MT-CYB missense'])]:
        d = sbs[sbs.group.isin(gsel)]
        # keep strata that contain at least one case
        cases = d.group == gsel[0]
        keep = d.stratum.isin(d.loc[cases, 'stratum'].unique())
        d = d[keep]
        if (d.group == gsel[0]).sum() < 2: continue
        for o in outcomes:
            dd = d.dropna(subset=[o])
            ox = [c for c in ('SBS18', 'SBS36') if c in dd]
            denom = dd.total - dd[ox].sum(axis=1) - (dd[o] if o not in ox and o != 'hole_burden' else 0)
            y = np.log((dd[o] + 0.5) / (denom.clip(lower=0) + 1)) if o != 'hole_burden' else np.log(dd[o] + 1e-3)
            g = (dd.group == gsel[0]).astype(int).values
            eff, lo, hi, p = strat_perm(y.values, g, dd.stratum.values, n=a.perms, rng=rng)
            frac1 = (dd.loc[g == 1, o] > 0).mean(); frac0 = (dd.loc[g == 0, o] > 0).mean()
            res.append(dict(comparison=comp, outcome=o, n_case=int(g.sum()), n_ctrl=int((1 - g).sum()),
                            ratio=math.exp(eff), lo=math.exp(lo), hi=math.exp(hi), p_perm=p,
                            frac_present_case=frac1, frac_present_ctrl=frac0))
    ass = pd.DataFrame(res); ass.to_csv(os.path.join(a.out, 'fig2_associations.tsv'), sep='\t', index=False)
    say(ass.round(3).to_string(index=False) if len(ass) else 'no associations computed (too few cases)')

    # ---------- figure ----------
    make_figure(mis, oe, sbs, ass, a)
    log.close()


def make_figure(mis, oe, sbs, ass, a):
    import matplotlib; matplotlib.use('Agg')
    import matplotlib.pyplot as plt
    from matplotlib.patches import Rectangle
    plt.rcParams.update({'font.family': 'DejaVu Sans', 'font.size': 7.5, 'axes.linewidth': 0.6,
                         'axes.edgecolor': '#52514e', 'xtick.color': '#52514e', 'ytick.color': '#52514e'})
    INK2, GRID = '#52514e', '#e4e3df'
    fig = plt.figure(figsize=(7.2, 7.0), dpi=300)
    gs = fig.add_gridspec(3, 3, height_ratios=[0.9, 1, 1.1], hspace=0.55, wspace=0.45,
                          left=0.08, right=0.98, top=0.95, bottom=0.07)
    lab = lambda ax, s: ax.text(-0.08, 1.04, s, transform=ax.transAxes, fontsize=11, fontweight='bold', ha='right', va='bottom')
    def clean(ax, left=True):
        for s in ('top', 'right') + (() if left else ('left',)): ax.spines[s].set_visible(False)
        ax.tick_params(length=2)

    # a lollipop
    ax = fig.add_subplot(gs[0, :]); lab(ax, 'a')
    cnt = mis.groupby(['aa_pos', 'category']).size().reset_index(name='n')
    for r in cnt.itertuples():
        c = COL[r.category] if r.category != 'other' else '#a8a7a2'
        ax.plot([r.aa_pos, r.aa_pos], [0, r.n], color=c, lw=0.8)
        ax.scatter(r.aa_pos, r.n, s=14, color=c, edgecolors='white', linewidths=0.5, zorder=3)
    cls_tab = pd.read_csv(a.classes, sep='\t')
    for r in cls_tab.itertuples():
        ax.add_patch(Rectangle((r.aa_pos - 0.5, -0.9), 1, 0.6, fc=COL[r.category], ec='none'))
    ax.set_xlim(0, 381); ax.set_ylim(-1.1, max(cnt.n.max() if len(cnt) else 1, 2) + 0.8)
    ax.set_xlabel('MT-CYB amino-acid position'); ax.set_ylabel('tumours'); clean(ax)
    ax.set_title(f'Somatic MT-CYB missense mutations (n = {len(mis)})', loc='left', fontsize=8, color=INK2)

    # b O/E
    ax = fig.add_subplot(gs[1, 0]); lab(ax, 'b')
    x = np.arange(len(oe))
    for i, r in oe.iterrows():
        ax.plot([i, i], [r.oe_lo, r.oe_hi], color=INK2, lw=0.8)
        ax.scatter(i, r.oe, s=28, color=COL[r.category], edgecolors='white', linewidths=0.6, zorder=3)
        ax.text(i, -0.05, f'{r.observed}', ha='center', va='top', fontsize=6, color=INK2, transform=ax.get_xaxis_transform())
    ax.axhline(1, color='#9a9893', lw=0.6, ls='--')
    ax.set_xticks(x); ax.set_xticklabels(['Qo', 'ISP', 'Qi', 'haem', 'other'])
    ax.tick_params(axis='x', pad=9)
    ax.set_ylabel('observed / expected'); clean(ax)
    ax.set_title('Missense by class', loc='left', fontsize=8, color=INK2)

    # c VAF
    ax = fig.add_subplot(gs[1, 1]); lab(ax, 'c')
    rng = np.random.default_rng(0)
    for i, c in enumerate(CLASSES):
        v = mis.loc[mis.category == c, 'vaf'].values
        ax.scatter(i + rng.uniform(-0.18, 0.18, len(v)), v, s=8, color=COL[c] if c != 'other' else '#a8a7a2', lw=0)
        if len(v): ax.plot([i - 0.25, i + 0.25], [np.median(v)] * 2, color='#0b0b0b', lw=1)
    ax.set_xticks(range(5)); ax.set_xticklabels(['Qo', 'ISP', 'Qi', 'haem', 'other'])
    ax.set_ylim(0, 1.02); ax.set_ylabel('heteroplasmy (VAF)'); clean(ax)
    ax.set_title('Heteroplasmy', loc='left', fontsize=8, color=INK2)

    # d SBS18 distribution by group
    ax = fig.add_subplot(gs[1, 2]); lab(ax, 'd')
    groups = ['no MT-CYB missense', 'other MT-CYB missense', 'Qo/ISP missense']
    gcol = ['#b5b4ae', '#7d7c77', '#2a78d6']
    if 'SBS18' in sbs:
        for i, (g, c) in enumerate(zip(groups, gcol)):
            v = (sbs.loc[sbs.group == g, 'SBS18'] / sbs.loc[sbs.group == g, 'total']).values * 100
            if len(v) > 200: ax.violinplot(v, [i], widths=0.7, showextrema=False)['bodies'][0].set(fc=c, alpha=0.5, ec='none')
            else: ax.scatter(i + rng.uniform(-0.15, 0.15, len(v)), v, s=9, color=c, lw=0)
            if len(v): ax.plot([i - 0.25, i + 0.25], [np.median(v)] * 2, color='#0b0b0b', lw=1)
            ax.text(i, -0.05, f'n={len(v)}', transform=ax.get_xaxis_transform(), ha='center', va='top', fontsize=6, color=INK2)
    ax.set_xticks(range(3)); ax.set_xticklabels(['none', 'other', 'Qo/ISP']); ax.tick_params(axis='x', pad=9)
    ax.set_ylabel('SBS18 (% of SBS)'); clean(ax)
    ax.set_title('SBS18 by MT-CYB status', loc='left', fontsize=8, color=INK2)

    # e forest
    ax = fig.add_subplot(gs[2, :2]); lab(ax, 'e')
    if len(ass):
        d = ass.iloc[::-1].reset_index(drop=True)
        for i, r in d.iterrows():
            c = '#2a78d6' if r.comparison.startswith('Qo') else '#7d7c77'
            ax.plot([r.lo, r.hi], [i, i], color=c, lw=1)
            ax.scatter(r.ratio, i, s=24, color=c, edgecolors='white', linewidths=0.6, zorder=3)
            ax.text(1.02, i, f'{r.ratio:.2f} ({r.lo:.2f}–{r.hi:.2f})  P = {r.p_perm:.2g}', transform=ax.get_yaxis_transform(),
                    va='center', fontsize=6.3, color=INK2)
        ax.set_yticks(range(len(d)))
        ax.set_yticklabels([f"{r.outcome} · {'Qo/ISP' if r.comparison.startswith('Qo') else 'other'} (n={r.n_case})" for r in d.itertuples()], fontsize=6.5)
        ax.set_xscale('log'); ax.axvline(1, color='#9a9893', lw=0.6, ls='--')
    ax.set_xlabel('exposure ratio vs tumours without MT-CYB missense (stratified by cancer type)'); clean(ax)
    ax.set_title('Association with nuclear mutational processes', loc='left', fontsize=8, color=INK2)

    # f SBS18 vs VAF in Qo/ISP mutants
    ax = fig.add_subplot(gs[2, 2]); lab(ax, 'f')
    hit = mis[mis.category.isin(LEAK)].groupby('sample').vaf.max()
    if 'SBS18' in sbs and len(hit):
        d = sbs.set_index('sample').reindex(hit.index).dropna(subset=['SBS18'])
        x = hit.loc[d.index].values; y = (d.SBS18 / d.total).values * 100
        ax.scatter(x, y, s=14, color='#2a78d6', edgecolors='white', linewidths=0.5)
        if len(x) >= 4:
            rho, p = stats.spearmanr(x, y)
            ax.text(0.03, 0.95, f'ρ = {rho:.2f}, P = {p:.2g}', transform=ax.transAxes, va='top', fontsize=6.5, color=INK2)
    ax.set_xlabel('max Qo/ISP VAF'); ax.set_ylabel('SBS18 (% of SBS)'); clean(ax)
    ax.set_title('Dose–response', loc='left', fontsize=8, color=INK2)

    fig.savefig(os.path.join(a.out, 'Figure2.png')); fig.savefig(os.path.join(a.out, 'Figure2.pdf'))


if __name__ == '__main__':
    ap = argparse.ArgumentParser()
    ap.add_argument('--mut', required=True)
    ap.add_argument('--mut-cols', default='sample,pos,ref,alt,vaf,cancer_type')
    ap.add_argument('--sbs', required=True)
    ap.add_argument('--sbs-sample-col', default='Sample Names')
    ap.add_argument('--sbs-type-col', default='Cancer Types')
    ap.add_argument('--hole')
    ap.add_argument('--classes', default=ASSETDIR + '/MT-CYB_residue_classes.tsv')
    ap.add_argument('--rcrs', default=ASSETDIR + '/rCRS.fasta')
    ap.add_argument('--signatures', default='SBS18,SBS36,SBS1,SBS5,SBS40')
    ap.add_argument('--vaf-min', type=float, default=0.05)
    ap.add_argument('--perms', type=int, default=5000)
    ap.add_argument('--out', default=OUTDIR + '/fig2_out')
    main(ap.parse_args())
```
