## Supplementary code for "Relaxed evolutionary constraints on mitochondrial electron-transfer-associated variants in human cancers": fig2_plot.py

```
import os as _os, sys as _sys
_sys.path.insert(0, _os.path.dirname(_os.path.abspath(__file__)))
from config import DATADIR, DATAROOT, EXTDIR, STRUCTDIR, ASSETDIR, OUTDIR
import json
import numpy as np, pandas as pd
import matplotlib; matplotlib.use('Agg')
import matplotlib.pyplot as plt
import matplotlib.ticker
from matplotlib.patches import Rectangle
from matplotlib.lines import Line2D
import sys; sys.path.insert(0, _os.path.dirname(_os.path.abspath(__file__)))
import fig2_pipeline as fp

import os
O = os.environ.get('FIG2_IN', OUTDIR + '/fig2_out_poly10/')
SUFFIX = os.environ.get('FIG2_SUFFIX', '')
plt.rcParams.update({'font.family': 'DejaVu Sans', 'font.size': 7.5, 'axes.linewidth': 0.6, 'axes.edgecolor': '#000000',
                     'xtick.color': '#000000', 'ytick.color': '#000000', 'text.color': '#000000'})
INK, INK2, MUTED, GRID = '#000000', '#000000', '#9a9893', '#e4e3df'
COL = dict(fp.COL); COL['other'] = '#a8a7a2'
SH = {'Qo site / bL leak path': 'Qo', 'ISP interface': 'ISP', 'Qi site': 'Qi', 'heme b environment': 'haem', 'other': 'other', 'Qo+ISP': 'Qo+ISP'}
oe = pd.read_csv(O + 'fig2_obs_exp_real.tsv', sep='\t')
som = pd.read_csv(O + 'fig2_mtcyb_missense_somatic.tsv', sep='\t')
con = pd.read_csv(O + 'fig2_somatic_vs_germline.tsv', sep='\t')
# (the nuclear-signature tables are not used by any Fig 2 panel and are no longer loaded here)
cls = pd.read_csv(ASSETDIR + '/MT-CYB_residue_classes.tsv', sep='\t')

fig = plt.figure(figsize=(7.2, 7.4), dpi=300)
gs = fig.add_gridspec(3, 3, height_ratios=[0.85, 1.05, 0.95], hspace=0.62, wspace=0.5, left=0.08, right=0.97, top=0.96, bottom=0.07)
def lab(ax, s, x=-0.1): ax.text(x, 1.05, s, transform=ax.transAxes, fontsize=11, fontweight='bold', ha='right', va='bottom')
def clean(ax):
    for s in ('top', 'right'): ax.spines[s].set_visible(False)
    ax.tick_params(length=2)
def title(ax, t): ax.set_title(t, loc='left', fontsize=8, color=INK2)

# a  lollipop
ax = fig.add_subplot(gs[0, :]); lab(ax, 'a', -0.04)
cnt = som.groupby(['aa_pos', 'category', 'cohort']).size().unstack(fill_value=0)
tot = cnt.sum(axis=1)
for (pos, cat), n in tot.items():
    ax.plot([pos, pos], [0, n], color=COL[cat], lw=0.7)
    ax.scatter(pos, n, s=12, color=COL[cat], edgecolors='white', linewidths=0.4, zorder=3)
for r in cls.itertuples():
    ax.add_patch(Rectangle((r.aa_pos - 0.5, -1.6), 1, 0.9, fc=fp.COL[r.category], ec='none'))
top = [(k, n) for k, n in tot.items() if k[0] in (34, 48, 80, 100, 116, 167)]
for (pos, cat), n in top:
    aa = som[som.aa_pos == pos].iloc[0]
    ax.text(pos, n + 0.3, f'{aa.aa_ref}{pos}', ha='center', va='bottom', fontsize=6, color=INK)
ax.set_xlim(0, 381); ax.set_ylim(-1.9, tot.max() + 1.8)
ax.set_xlabel('MT-CYB amino-acid position'); ax.set_ylabel('tumours'); clean(ax)
title(ax, f'Somatic MT-CYB missense mutations, TCGA + PCAWG (n = {len(som)})')
hand = [Line2D([], [], marker='s', ls='', color=fp.COL[c], markersize=6, label=SH[c]) for c in fp.CLASSES]
ax.legend(handles=hand, ncol=5, frameon=False, fontsize=6.5, loc='upper right', bbox_to_anchor=(1, 1.12), handletextpad=0.3, columnspacing=1.0)

# b  observed / expected
ax = fig.add_subplot(gs[1, :2]); lab(ax, 'b', -0.06)
cats = fp.CLASSES + ['Qo+ISP']
coh = [('Germline (HelixMTdb)', 'o', 'white'), ('Somatic TCGA', 's', None), ('Somatic PCAWG', '^', None)]
for i, c in enumerate(cats):
    if c == 'Qo+ISP': ax.axvspan(i - 0.45, i + 0.45, color=GRID, alpha=0.6, lw=0)
    for j, (name, mk, fc) in enumerate(coh):
        r = oe[(oe.cohort == name) & (oe.category == c)].iloc[0]
        x = i + (j - 1) * 0.24; col = COL.get(c, '#2a78d6' if c == 'Qo+ISP' else INK2)
        if c == 'Qo+ISP': col = '#2a78d6'
        ax.plot([x, x], [r.lo, r.hi], color=col, lw=0.9)
        ax.scatter(x, r.oe, marker=mk, s=26, facecolor=fc or col, edgecolor=col, linewidths=1.0, zorder=3)
        if r.p < 0.05: ax.text(x, r.hi + 0.05, '*' if r.p >= 0.01 else '**', ha='center', va='bottom', fontsize=8, color=INK)
ax.axhline(1, color=MUTED, lw=0.6, ls='--')
ax.set_xticks(range(len(cats))); ax.set_xticklabels([SH[c] for c in cats]); ax.set_ylim(0.2, 2.6)
ax.set_ylabel('missense observed / expected'); clean(ax)
title(ax, 'Selection by structural class (context- and strand-aware null)')
ax.legend(handles=[Line2D([], [], marker=m, ls='', mfc=fc or INK2, mec=INK2, color=INK2, markersize=5, label=n.replace('Somatic ', ''))
                   for n, m, fc in coh], frameon=False, fontsize=6.5, loc='upper right', ncol=3, handletextpad=0.2)

# c  somatic vs germline share
ax = fig.add_subplot(gs[1, 2]); lab(ax, 'c', -0.18)
d = con.iloc[::-1].reset_index(drop=True)
for i, r in d.iterrows():
    tab = np.array([[r.somatic_in, r.somatic_out], [r.germline_in, r.germline_out]], float)
    se = np.sqrt((1 / tab).sum()); lo, hi = np.exp(np.log(r.OR_somatic_vs_germline) + np.array([-1.96, 1.96]) * se)
    col = '#2a78d6' if r.category == 'Qo+ISP' else COL[r.category]
    ax.plot([lo, hi], [i, i], color=col, lw=1); ax.scatter(r.OR_somatic_vs_germline, i, s=24, color=col, edgecolors='white', lw=0.6, zorder=3)
    ax.text(3.9, i, f'P = {r.p:.2g}', va='center', ha='left', fontsize=6, color=INK2)
ax.set_yticks(range(len(d))); ax.set_yticklabels([SH[c] for c in d.category])
ax.set_xscale('log'); ax.set_xlim(0.45, 9); ax.set_xticks([0.5, 1, 2]); ax.set_xticklabels(['0.5', '1', '2']); ax.xaxis.set_minor_formatter(matplotlib.ticker.NullFormatter())
ax.axvline(1, color=MUTED, lw=0.6, ls='--'); ax.set_xlabel('odds ratio, somatic vs germline'); clean(ax)
title(ax, 'Relaxed constraint')

# d  charge change x class
ax = fig.add_subplot(gs[2, :]); lab(ax, 'd', -0.04)
cs = pd.read_csv(os.environ.get('FIG2_CHARGE', OUTDIR + '/fig2_charge_split.tsv'), sep='\t')
from scipy.stats import chi2
def pois_ci(n, e):
    lo = chi2.ppf(0.025, 2 * n) / 2 / e if n > 0 else 0.0
    return lo, chi2.ppf(0.975, 2 * n + 2) / 2 / e
groups = [('haem', 'heme b environment'), ('Qi', 'Qi site'), ('Qo/ISP', 'Qo site / bL leak path'), ('other', 'other')]
xt, xl = [], []
for gi, (g, ckey) in enumerate(groups):
    for ci_, ch in enumerate(['charge', 'neutral']):
        x0 = gi * 2.6 + ci_ * 1.0
        cell = f'{g} · {ch}'
        for k, (lab_, sel, mk, fcol) in enumerate([('germline', cs.cohort == 'germline', 'o', 'white'),
                                                  ('tumour', cs.cohort != 'germline', 's', None)]):
            d = cs[sel & (cs.cell == cell)]; n, e = int(d.observed.sum()), d.expected.sum()
            lo, hi = pois_ci(n, e); x = x0 + (k - 0.5) * 0.36
            col = COL[ckey] if ch == 'charge' else '#a8a7a2'
            ax.plot([x, x], [lo, hi], color=col, lw=1.0)
            ax.scatter(x, n / e, marker=mk, s=30, facecolor=fcol or col, edgecolor=col, linewidths=1.1, zorder=3)
            if g == 'haem' and ch == 'charge':
                ax.text(x + (0.08 if k else -0.08), n / e, f'{n / e:.2f}', ha='left' if k else 'right', va='center', fontsize=6.2, color=INK)
        xt.append(x0); xl.append('charge\nchange' if ch == 'charge' else 'neutral\n ')
    ax.text(gi * 2.6 + 0.5, 4.75, {'haem': 'haem b environment', 'Qi': 'Qi site', 'Qo/ISP': 'Qo site + ISP interface', 'other': 'other residues'}[g],
            ha='center', va='bottom', fontsize=6.8, color=INK2)
ax.axhline(1, color=MUTED, lw=0.6, ls='--')
ax.set_xticks(xt); ax.set_xticklabels(xl, fontsize=6.5); ax.set_xlim(-0.6, 3 * 2.6 + 1.6)
ax.set_yscale('log'); ax.set_ylim(0.02, 6); ax.set_yticks([0.05, 0.1, 0.25, 0.5, 1, 2, 4]); ax.set_yticklabels(['0.05', '0.1', '0.25', '0.5', '1', '2', '4'])
ax.yaxis.set_minor_formatter(matplotlib.ticker.NullFormatter())
ax.set_ylabel('missense observed / expected'); clean(ax)
title(ax, 'Charge-changing substitutions near the haems: purged in the germline, enriched in tumours')
ax.legend(handles=[Line2D([], [], marker='o', ls='', mfc='white', mec=INK2, markersize=5, label='germline (HelixMTdb)'),
                   Line2D([], [], marker='s', ls='', color=INK2, markersize=5, label='tumour (TCGA + PCAWG)')],
          frameon=False, fontsize=6.5, loc='lower right', ncol=2, handletextpad=0.2)
fig.savefig(f'{OUTDIR}/Figure2{SUFFIX}.png'); fig.savefig(f'{OUTDIR}/Figure2{SUFFIX}.pdf')
```
