## Supplementary code for "Relaxed evolutionary constraints on mitochondrial electron-transfer-associated variants in human cancers": fig2_tables.py

```
"""Figure 2 on real data.
Somatic mtDNA SNVs: TCGA (WES off-target) and PCAWG (WGS) calls from Gorelick et al. 2021
(github.com/reznik-lab/mtdna-mutations). Germline: HelixMTdb variant sites (same repository).
Nuclear SBS exposures: tcga_sbs_exposures.csv (nuclear_sbs.py, COSMIC v3.4 exome fit).
"""
import os as _os, sys as _sys
_sys.path.insert(0, _os.path.dirname(_os.path.abspath(__file__)))
from config import DATADIR, DATAROOT, EXTDIR, STRUCTDIR, ASSETDIR, OUTDIR

import sys, math, json
import numpy as np, pandas as pd
from scipy import stats
sys.path.insert(0, _os.path.dirname(_os.path.abspath(__file__)))
import fig2_pipeline as fp

R = DATAROOT
OUT = OUTDIR + '/fig2_out'
import os; os.makedirs(OUT, exist_ok=True)
log = open(f'{OUT}/fig2_real_log.txt', 'w')
def say(*x): print(*x); print(*x, file=log)

seq = fp.load_rcrs(ASSETDIR + '/rCRS.fasta')
ct = pd.read_csv(ASSETDIR + '/MT-CYB_residue_classes.tsv', sep='\t'); cls = dict(zip(ct.aa_pos, ct.category))
SHORT = {'Qo site / bL leak path': 'Qo', 'ISP interface': 'ISP', 'Qi site': 'Qi', 'heme b environment': 'haem', 'other': 'other'}


def load(path, sample_col='Tumor_Sample_Barcode', somatic_only=False):
    d = pd.read_csv(path, sep='\t', low_memory=False)
    if somatic_only: d = d[d['class'] == 'somatic']
    d = d[d.Variant_Type == 'SNP']
    o = pd.DataFrame(dict(sample=d[sample_col].values, pos=d.Start_Position.astype(int).values,
                          ref=d.Reference_Allele.str.upper().values, alt=d.Tumor_Seq_Allele2.str.upper().values,
                          vaf=d['TumorVAF'].values if 'TumorVAF' in d else np.nan,
                          hgvsp=d.HGVSp_Short.values,
                          cancer=d['CANCER_TYPE'].values if 'CANCER_TYPE' in d else (d['project'].values if 'project' in d else 'NA'),
                          patient=d['PATIENT_ID'].values if 'PATIENT_ID' in d else d[sample_col].values))
    bad = (o.ref != o.pos.map(lambda p: seq[p - 1])).sum()
    o = o[o.ref == o.pos.map(lambda p: seq[p - 1])]
    return o, bad


def annotate(o):
    c = o[(o.pos >= fp.CYB_START) & (o.pos <= fp.CYB_END - 1)].copy()
    ann = c.apply(lambda r: fp.annotate_cyb(r.pos, r.alt, seq), axis=1, result_type='expand')
    c[['aa_pos', 'aa_ref', 'aa_alt', 'effect']] = ann
    c['category'] = c.aa_pos.map(cls).fillna('other')
    # cross-check with published HGVSp
    hg = c.hgvsp.astype(str).str.extract(r'p\.([A-Z\*])(\d+)([A-Z\*])')
    ok = hg[1].notna()
    agree = ((hg.loc[ok, 1].astype(int) == c.loc[ok, 'aa_pos']) & (hg.loc[ok, 0] == c.loc[ok, 'aa_ref'])).mean()
    return c, agree


def obs_exp(snv, mis, skip=None):
    excl = set(zip(mis['sample'], mis.pos, mis.alt))
    f = fp.expected_fractions(seq, fp.spectrum(snv, seq, excl), cls, skip)
    N = len(mis); rows = []
    for c in fp.CLASSES:
        n = int((mis.category == c).sum())
        bt = stats.binomtest(n, N, f[c]); ci = bt.proportion_ci()
        rows.append(dict(category=c, observed=n, expected=N * f[c], oe=n / (N * f[c]), lo=ci[0] / f[c], hi=ci[1] / f[c], p=bt.pvalue))
    lk = int(mis.category.isin(fp.LEAK).sum()); lf = sum(f[c] for c in fp.LEAK); bt = stats.binomtest(lk, N, lf); ci = bt.proportion_ci()
    rows.append(dict(category='Qo+ISP', observed=lk, expected=N * lf, oe=lk / (N * lf), lo=ci[0] / lf, hi=ci[1] / lf, p=bt.pvalue))
    return pd.DataFrame(rows)


POLY_MAX = int(os.environ.get('POLY_MAX', '10'))  # default: population-rare filter (manuscript main); set 1000000000 for unfiltered (Supp Fig 1)
cohorts = {}
t, bt_ = load(R + 'original_data/data_mutations_tcga.txt.gz', somatic_only=True)
p, bp_ = load(R + 'original_data/data_mutations_pcawg.txt.gz')
h, bh_ = load(R + 'original_data/data_mutations_helix.txt.gz')
h['sample'] = 'helix_' + h.pos.astype(str) + h.alt     # site-level: each distinct variant once
say(f'ref mismatches removed: TCGA {bt_}, PCAWG {bp_}, Helix {bh_}')
_hc = pd.read_csv(R + 'original_data/data_mutations_helix.txt.gz', sep='\t', low_memory=False)
_hc = _hc[_hc.Variant_Type == 'SNP'].groupby(['Start_Position', 'Tumor_Seq_Allele2']).TOTAL_COUNT.sum().to_dict()
if POLY_MAX < 10**9:
    for _n, _d in [('t', t), ('p', p)]:
        keep = [(_hc.get((a, b), 0) <= POLY_MAX) for a, b in zip(_d.pos, _d.alt)]
        say(f'polymorphism filter (Helix carriers <= {POLY_MAX}): {_n} removed {len(_d) - sum(keep)}')
    t = t[[(_hc.get((a, b), 0) <= POLY_MAX) for a, b in zip(t.pos, t.alt)]]
    p = p[[(_hc.get((a, b), 0) <= POLY_MAX) for a, b in zip(p.pos, p.alt)]]
    OUT = OUT + f'_poly{POLY_MAX}'; os.makedirs(OUT, exist_ok=True)
POLYSITES = {k for k, v in _hc.items() if v > POLY_MAX} if POLY_MAX < 10**9 else set()
OE = []
for name, d in [('Germline (HelixMTdb)', h), ('Somatic TCGA', t), ('Somatic PCAWG', p)]:
    c, agree = annotate(d); mis = c[c.effect == 'missense']
    say(f'{name}: SNVs {len(d)}, MT-CYB missense {len(mis)}, HGVSp agreement {agree:.3f}')
    skip = POLYSITES if (POLY_MAX < 10**9 and name != 'Germline (HelixMTdb)') else None
    oe = obs_exp(d, mis, skip); oe['cohort'] = name; OE.append(oe); cohorts[name] = mis
    say(oe.round(3).to_string(index=False))
# pooled somatic
som = pd.concat([cohorts['Somatic TCGA'].assign(cohort='TCGA'), cohorts['Somatic PCAWG'].assign(cohort='PCAWG')])
OE = pd.concat(OE); OE.to_csv(f'{OUT}/fig2_obs_exp_real.tsv', sep='\t', index=False)
som.to_csv(f'{OUT}/fig2_mtcyb_missense_somatic.tsv', sep='\t', index=False)

# germline vs somatic contrast for Qo+ISP and for each class (Fisher on class vs other, germline sites vs somatic)
con = []
gm = cohorts['Germline (HelixMTdb)']
if POLY_MAX < 10**9: gm = gm[[(a, b) not in POLYSITES for a, b in zip(gm.pos, gm.alt)]]   # same site universe as somatic
for c in fp.CLASSES[:4] + ['Qo+ISP']:
    sel = (lambda d: d.category.isin(fp.LEAK)) if c == 'Qo+ISP' else (lambda d, c=c: d.category == c)
    a1, a0 = int(sel(som).sum()), int((~sel(som)).sum()); b1, b0 = int(sel(gm).sum()), int((~sel(gm)).sum())
    orr, pv = stats.fisher_exact([[a1, a0], [b1, b0]])
    con.append(dict(category=c, somatic_in=a1, somatic_out=a0, germline_in=b1, germline_out=b0, OR_somatic_vs_germline=orr, p=pv))
con = pd.DataFrame(con); con.to_csv(f'{OUT}/fig2_somatic_vs_germline.tsv', sep='\t', index=False)
say('\nSomatic vs germline class share (Fisher):'); say(con.round(4).to_string(index=False))

# VAF by class (somatic, TCGA+PCAWG)
v_leak = som.loc[som.category.isin(fp.LEAK), 'vaf']; v_oth = som.loc[~som.category.isin(fp.LEAK), 'vaf']
mw = stats.mannwhitneyu(v_leak, v_oth)
say(f'\nVAF Qo/ISP median {v_leak.median():.3f} (n={len(v_leak)}) vs other classes {v_oth.median():.3f} (n={len(v_oth)}); MWU p={mw.pvalue:.3g}')

# ---------- nuclear signatures (TCGA) ----------
if not os.path.exists(OUTDIR + '/tcga_sbs_exposures.csv'):
    say('\n[skip] nuclear-signature association: tcga_sbs_exposures.csv not found (run nuclear_sbs.py first)')
    sys.exit(0)
e = pd.read_csv(OUTDIR + '/tcga_sbs_exposures.csv'); q = pd.read_csv(OUTDIR + '/tcga_sbs_qc.tsv', sep='\t')
e = e.merge(q[['sample', 'n_snv', 'gg_CtoA']], left_on='Sample Names', right_on='sample')
L = pd.read_csv(R + 'processed_data/tcga_sample_gene_lengths_precalculated.txt.gz', sep='\t')
L = L[L.sample_type == 'tumor+normal']; callable_p = set(L.loc[L['MT-CYB'] >= 0.9 * 1141, 'Tumor_Sample_Barcode'].str[:12])
e = e[e.Patient.isin(callable_p)].copy()
sig = [c for c in e.columns if c.startswith('SBS')]
e['total'] = e[sig].sum(axis=1)
tm = cohorts['Somatic TCGA']
leak_p = set(tm.loc[tm.category.isin(fp.LEAK), 'patient']); any_p = set(tm.patient)
e['group'] = np.where(e.Patient.isin(leak_p), 'Qo/ISP missense', np.where(e.Patient.isin(any_p), 'other MT-CYB missense', 'no MT-CYB missense'))
e['stratum'] = e['Cancer Types']
say(f"\nTCGA tumours with nuclear SBS and callable MT-CYB: {len(e)}; groups {e.group.value_counts().to_dict()}")
e.to_csv(f'{OUT}/fig2_tcga_groups.tsv', sep='\t', index=False)
outs = ['SBS18', 'SBS36', 'gg_CtoA', 'SBS1', 'SBS5', 'SBS40a']
res = []; rng = np.random.default_rng(1)
for label, sub in [('all', e), ('>=50 SNVs', e[e.n_snv >= 50])]:
    for comp, gsel in [('Qo/ISP', ['Qo/ISP missense', 'no MT-CYB missense']), ('other MT-CYB', ['other MT-CYB missense', 'no MT-CYB missense'])]:
        d = sub[sub.group.isin(gsel)]
        d = d[d.stratum.isin(d.loc[d.group == gsel[0], 'stratum'].unique())]
        for o in outs:
            ox = ['SBS18', 'SBS36']
            denom = d.total - d[ox].sum(axis=1) - (d[o] if o not in ox else 0)
            if o == 'gg_CtoA': denom = d.n_snv - d[o]
            y = np.log((d[o] + 0.5) / (denom.clip(lower=0) + 1))
            g = (d.group == gsel[0]).astype(int).values
            eff, lo, hi, pv = fp.strat_perm(y.values, g, d.stratum.values, n=10000, rng=rng)
            res.append(dict(set=label, comparison=comp, outcome=o, n_case=int(g.sum()), n_ctrl=int((1 - g).sum()),
                            ratio=math.exp(eff), lo=math.exp(lo), hi=math.exp(hi), p=pv,
                            present_case=(d.loc[g == 1, o] > 0).mean(), present_ctrl=(d.loc[g == 0, o] > 0).mean()))
res = pd.DataFrame(res); res.to_csv(f'{OUT}/fig2_associations_real.tsv', sep='\t', index=False)
say(res.round(3).to_string(index=False))
# SBS18 presence: stratified (CMH) test Qo/ISP vs none
from statsmodels.stats.contingency_tables import StratifiedTable
d = e[e.group.isin(['Qo/ISP missense', 'no MT-CYB missense'])]
tabs = []
for s, dd in d.groupby('stratum'):
    a = ((dd.group == 'Qo/ISP missense') & (dd.SBS18 > 0)).sum(); b = ((dd.group == 'Qo/ISP missense') & (dd.SBS18 == 0)).sum()
    c = ((dd.group != 'Qo/ISP missense') & (dd.SBS18 > 0)).sum(); dd_ = ((dd.group != 'Qo/ISP missense') & (dd.SBS18 == 0)).sum()
    if a + b > 0: tabs.append(np.array([[a, b], [c, dd_]]) + 0.0)
st = StratifiedTable(tabs)
say(f'SBS18 presence Qo/ISP vs none, CMH OR {st.oddsratio_pooled:.2f}, p={st.test_null_odds().pvalue:.3g}')
json.dump(dict(cmh_or=st.oddsratio_pooled, cmh_p=st.test_null_odds().pvalue, vaf_mwu_p=mw.pvalue,
               vaf_leak_median=v_leak.median(), vaf_other_median=v_oth.median()), open(f'{OUT}/fig2_stats.json', 'w'))
```
