## Supplementary code for "Relaxed evolutionary constraints on mitochondrial electron-transfer-associated variants in human cancers": fig3_plot.py

```
import os as _os, sys as _sys
_sys.path.insert(0, _os.path.dirname(_os.path.abspath(__file__)))
from config import DATADIR, DATAROOT, EXTDIR, STRUCTDIR, ASSETDIR, OUTDIR
import numpy as np, pandas as pd
import matplotlib; matplotlib.use('Agg')
import matplotlib.pyplot as plt, matplotlib.ticker
from matplotlib.lines import Line2D
from matplotlib.colors import TwoSlopeNorm, LinearSegmentedColormap
from scipy import stats

plt.rcParams.update({'font.family': 'DejaVu Sans', 'font.size': 7.5, 'axes.linewidth': 0.6, 'axes.edgecolor': '#000000',
                     'xtick.color': '#000000', 'ytick.color': '#000000', 'text.color': '#000000'})
INK, INK2, MUTED, GRID = '#000000', '#000000', '#9a9893', '#e4e3df'
TUM, GER = '#eb6834', '#2a78d6'
R = pd.read_csv(OUTDIR + '/fig3_oe.tsv', sep='\t')
V = pd.read_csv(OUTDIR + '/fig3_variants.tsv', sep='\t')
Q = pd.read_csv(OUTDIR + '/qcycle_validation.tsv', sep='\t')

fig = plt.figure(figsize=(7.2, 6.0), dpi=300)
gs = fig.add_gridspec(2, 3, height_ratios=[1, 1.15], width_ratios=[0.8, 1.15, 1.05], hspace=0.5, wspace=0.55,
                      left=0.08, right=0.97, top=0.91, bottom=0.09)
def lab(ax, s, x=-0.22): ax.text(x, 1.06, s, transform=ax.transAxes, fontsize=11, fontweight='bold', ha='right', va='bottom')
def clean(ax):
    for s in ('top', 'right'): ax.spines[s].set_visible(False)
    ax.tick_params(length=2)
def title(ax, t): ax.set_title(t, loc='left', fontsize=8, color=INK2)
def logy(ax, lo, hi, ticks):
    ax.set_yscale('log'); ax.set_ylim(lo, hi); ax.set_yticks(ticks); ax.set_yticklabels([f'{t:g}' for t in ticks])
    ax.yaxis.set_minor_formatter(matplotlib.ticker.NullFormatter())

# a  high-perturbation O/E
ax = fig.add_subplot(gs[0, 0]); lab(ax, 'a', -0.3)
h = R[R.analysis == 'high'].set_index('group')
for i, (g, mk, col, fc) in enumerate([('germline', 'o', GER, 'white'), ('TCGA', 's', TUM, None), ('PCAWG', '^', TUM, None)]):
    r = h.loc[g]
    ax.plot([i, i], [r.lo, r.hi], color=col, lw=1.1)
    ax.scatter(i, r.oe, marker=mk, s=36, facecolor=fc or col, edgecolor=col, lw=1.2, zorder=3)
    ax.text(i + 0.14, r.oe, f'{r.oe:.2f}', va='center', fontsize=6.3, color=INK)
    ax.text(i, r.hi * 1.08, f'P={r.p:.1g}', ha='center', va='bottom', fontsize=5.8, color=INK2)
ax.axhline(1, color=MUTED, lw=0.6, ls='--')
ax.set_xticks(range(3)); ax.set_xticklabels(['germline', 'TCGA', 'PCAWG'], fontsize=6.6); ax.set_xlim(-0.5, 2.6)
logy(ax, 0.3, 3, [0.5, 1, 2]); ax.set_ylabel('observed / expected'); clean(ax)
title(ax, 'ET-perturbing missense')

# b  dose-response
ax = fig.add_subplot(gs[0, 1]); lab(ax, 'b')
bins = ['<0.1', '0.1–0.3', '0.3–1', '≥1']
for g, mk, col, fc, ls in [('germline', 'o', GER, 'white', '--'), ('tumour', 's', TUM, None, '-')]:
    d = R[(R.analysis == 'bin') & (R.group == g)].set_index('bin').loc[bins]
    x = np.arange(4) + (-0.08 if g == 'germline' else 0.08)
    ax.plot(x, d.oe, color=col, lw=1.2, ls=ls)
    for xi, r in zip(x, d.itertuples()):
        ax.plot([xi, xi], [r.lo, r.hi], color=col, lw=0.9)
    ax.scatter(x, d.oe, marker=mk, s=28, facecolor=fc or col, edgecolor=col, lw=1.1, zorder=3)
    ax.text(x[-1] + 0.15, d.oe.iloc[-1], 'tumour' if g == 'tumour' else 'germline', color=col, fontsize=6.6, va='center')
ax.axhline(1, color=MUTED, lw=0.6, ls='--')
ax.set_xticks(range(4)); ax.set_xticklabels(bins); ax.set_xlim(-0.4, 4.0)
ax.set_xlabel('predicted ET-rate change, |Δlog$_{10}$ k|'); logy(ax, 0.3, 3.5, [0.5, 1, 2])
ax.set_ylabel('observed / expected'); clean(ax)
title(ax, 'Dose–response')

# c  sensitivity heatmaps
sub = gs[0, 2].subgridspec(1, 2, wspace=0.12)
cmap = LinearSegmentedColormap.from_list('div', [GER, '#f1f0ec', TUM])
norm = TwoSlopeNorm(vcenter=0.0, vmin=-1.0, vmax=1.0)
for k, g in enumerate(['germline', 'tumour']):
    ax = fig.add_subplot(sub[k])
    if k == 0: lab(ax, 'c', -0.55)
    s = R[(R.analysis == 'sens') & (R.group == g)].pivot_table(index='eps', columns='lam_scale', values='oe')
    ax.imshow(np.log2(s.values), cmap=cmap, norm=norm, aspect='auto')
    for i in range(3):
        for j in range(3):
            ax.text(j, i, f'{s.values[i, j]:.2f}', ha='center', va='center', fontsize=6, color=INK)
    ax.set_xticks(range(3)); ax.set_xticklabels(['0.7λ', 'λ', '1.4λ'], fontsize=6)
    ax.set_yticks(range(3)); ax.set_yticklabels([f'ε={e}' for e in s.index] if k == 0 else [], fontsize=6)
    ax.tick_params(length=0)
    for sp in ax.spines.values(): sp.set_visible(False)
    ax.set_title(g, fontsize=7, color=INK2)
fig.text(0.715, 0.965, 'Parameter sensitivity (O/E)', fontsize=8, color=INK2)

# d  network vs proximity, with tumour variants
ax = fig.add_subplot(gs[1, :2]); lab(ax, 'd', -0.1)
ax.scatter(V.prox_eV, V.network, s=3, color='#cfcdc7', lw=0, zorder=1, label='all possible missense')
b1, b0 = np.polyfit(V.prox_eV, V.network, 1)
xx = np.linspace(0, V.prox_eV.max(), 50); ax.plot(xx, b0 + b1 * xx, color=MUTED, lw=0.8, ls='--')
T = V[(V.tumours > 0) & (V.dq.abs() >= 0.9)].copy()
T['res'] = T.network - (b0 + b1 * T.prox_eV)
ax.scatter(T.prox_eV, T.network, s=8 + 6 * T.tumours, facecolor='none', edgecolor=TUM, lw=0.9, zorder=3, label='tumour-observed, charge-changing')
names = {(116, 'D'), (48, 'E'), (34, 'D'), (100, 'Q'), (202, 'K'), (80, 'H'), (231, 'D')}
for r in T.itertuples():
    if (r.aa_pos, r.aa_alt) in names:
        off = {116: (-0.07, 0.25), 48: (0.05, 0.3), 34: (-0.06, 0.2), 100: (-0.06, 0.25), 202: (0.06, -0.35), 80: (-0.02, -0.45), 231: (0.04, -0.3)}[r.aa_pos]
        ax.annotate(f'{r.aa_ref}{r.aa_pos}{r.aa_alt}', (r.prox_eV, r.network), (r.prox_eV + off[0], r.network + off[1]),
                    fontsize=6.3, ha='right' if off[0] < 0 else 'left', va='center', color=INK, arrowprops=dict(arrowstyle='-', color=MUTED, lw=0.5))
ax.text(0.02, 0.97, f'r = {np.corrcoef(V.prox_eV, V.network)[0, 1]:.2f} across all possible missense', transform=ax.transAxes, va='top', fontsize=6.5, color=INK2)
ax.set_xlabel('electrostatic shift at nearest redox centre, |ΔE$_m$| (eV)'); ax.set_ylabel('Q-cycle ET perturbation, |Δlog$_{10}$ k|')
ax.set_ylim(-0.1, 3.3); clean(ax)
ax.legend(frameon=False, fontsize=6.3, loc='lower right', markerscale=1.3, handletextpad=0.3)
title(ax, 'ET perturbation is collinear with local electrostatics; outliers discriminate')

# e  ET-specific component
ax = fig.add_subplot(gs[1, 2]); lab(ax, 'e', -0.3)
q = Q[(Q.eps == 8) & (Q.lam_scale == 1.0)].set_index('cohort').loc[['germline', 'TCGA', 'PCAWG']]
for i, (g, r) in enumerate(q.iterrows()):
    se = abs(r.b_diff_only) / max(stats.norm.isf(r.p_diff_only / 2), 1e-6)
    col = GER if g == 'germline' else TUM
    ax.plot([r.b_diff_only - 1.96 * se, r.b_diff_only + 1.96 * se], [2 - i] * 2, color=col, lw=1.1)
    ax.scatter(r.b_diff_only, 2 - i, s=30, color=col, edgecolors='white', lw=0.6, zorder=3)
    ax.text(0.98, 2 - i + 0.22, f'P = {r.p_diff_only:.2f}', transform=ax.get_yaxis_transform(), ha='right', fontsize=6.2, color=INK2)
ax.axvline(0, color=MUTED, lw=0.6, ls='--')
ax.set_yticks([2, 1, 0]); ax.set_yticklabels(['germline', 'TCGA', 'PCAWG']); ax.set_ylim(-0.6, 2.7)
ax.set_xlabel('effect per s.d. (log rate ratio)'); clean(ax)
ax.set_title('ET-specific component', loc='left', fontsize=8, color=INK2, pad=12)
ax.text(0.0, 1.0, 'beyond |ΔE$_m$| and chemistry covariates', transform=ax.transAxes, fontsize=6, color=INK2, va='bottom')

fig.savefig(OUTDIR + '/Figure3.png'); fig.savefig(OUTDIR + '/Figure3.pdf')
```
