## Supplementary code for "Relaxed evolutionary constraints on mitochondrial electron-transfer-associated variants in human cancers": fig3_tables.py

```
"""Figure 3 data: Q-cycle (multi-step) Marcus ET-perturbation score for all possible MT-CYB missense SNVs.
Spectrum-aware O/E (each cohort's own 192-channel rates) for score bins, the high-perturbation class
(score >= 0.3, i.e. >= 2-fold change in at least one ET step), and a parameter sensitivity grid."""
import os as _os, sys as _sys
_sys.path.insert(0, _os.path.dirname(_os.path.abspath(__file__)))
from config import DATADIR, DATAROOT, EXTDIR, STRUCTDIR, ASSETDIR, OUTDIR

import sys, json
sys.path.insert(0, _os.path.dirname(_os.path.abspath(__file__)))
import numpy as np, pandas as pd
from scipy import stats
import et_variants as ev, qcycle_marcus as qm

F = ev.features('CIII bL->bH')
C = ev.cohorts(); RT = {k: ev.rates(v) for k, v in C.items()}
MASK = ev.poly_ok(F)
W = {k: np.array([RT[k][(t, a)] for t, a in zip(F.tri, F.alt)]) * (1 if k == 'germline' else MASK) for k in C}
Y = {}
for k, d in C.items():
    key = d.groupby(['pos', 'alt']).size()
    y = np.array([key.get((p, a), 0) for p, a in zip(F.pos, F.alt)])
    Y[k] = (y > 0).astype(int) if k == 'germline' else y


def oe(mask, cohorts):
    n = sum(int(Y[c][mask].sum()) for c in cohorts)
    e = sum(Y[c].sum() * W[c][mask].sum() / W[c].sum() for c in cohorts)
    lo = stats.chi2.ppf(0.025, 2 * n) / 2 / e if n else 0.0
    hi = stats.chi2.ppf(0.975, 2 * n + 2) / 2 / e
    p = stats.poisson.cdf(n, e) * 2 if n < e else stats.poisson.sf(n - 1, e) * 2
    return dict(observed=n, expected=e, oe=n / e, lo=lo, hi=hi, p=min(p, 1.0))


GROUPS = {'germline': ['germline'], 'TCGA': ['TCGA'], 'PCAWG': ['PCAWG'], 'tumour': ['TCGA', 'PCAWG']}
net, blbh, prox = qm.scores(F, 8.0, 1.0)
F['network'], F['prox_eV'] = net, prox
rows = []
for g, cs in GROUPS.items():
    rows.append(dict(analysis='high', group=g, **oe(net >= 0.3, cs)))
    for lo_, hi_, lab in [(0, 0.1, '<0.1'), (0.1, 0.3, '0.1–0.3'), (0.3, 1.0, '0.3–1'), (1.0, 99, '≥1')]:
        rows.append(dict(analysis='bin', bin=lab, group=g, **oe((net >= lo_) & (net < hi_), cs)))
for eps in (4, 8, 16):
    for sl in (0.7, 1.0, 1.4):
        n_, _, _ = qm.scores(F, float(eps), sl)
        for g in ('germline', 'tumour'):
            rows.append(dict(analysis='sens', eps=eps, lam_scale=sl, group=g, **oe(n_ >= 0.3, GROUPS[g])))
R = pd.DataFrame(rows); R.to_csv(OUTDIR + '/fig3_oe.tsv', sep='\t', index=False)
F[['pos', 'alt', 'aa_pos', 'aa_ref', 'aa_alt', 'dq', 'network', 'prox_eV']].assign(
    tumours=Y['TCGA'] + Y['PCAWG'], germline=Y['germline']).to_csv(OUTDIR + '/fig3_variants.tsv', sep='\t', index=False)
pd.set_option('display.width', 200)
print(R[R.analysis != 'sens'].round(3).to_string(index=False))
print(R[R.analysis == 'sens'].pivot_table(index=['eps', 'lam_scale'], columns='group', values='oe').round(2))
print('r(network, prox) =', round(np.corrcoef(net, prox)[0, 1], 3))
```
