## Supplementary code for "Relaxed evolutionary constraints on mitochondrial electron-transfer-associated variants in human cancers": fig4_helix_state.py

```
"""Fig 4b source: germline (HelixMTdb) MT-CYB missense sites classified as homoplasmic in >=1 person
(HOM_COUNT > 0) versus observed only heteroplasmic (HOM_COUNT == 0), split by predicted high-ET
perturbation. Writes fig4_helix_state.tsv."""
import os as _os, sys as _sys
_sys.path.insert(0, _os.path.dirname(_os.path.abspath(__file__)))
from config import DATADIR, OUTDIR

import numpy as np, pandas as pd
import et_variants as ev, qcycle_marcus as qm

F = ev.features('CIII bL->bH')
net, _, _ = qm.scores(F, 8.0, 1.0)
high = {(int(p), a): bool(h) for p, a, h in zip(F.pos, F.alt, net >= 0.3)}

hc = pd.read_csv(DATADIR + 'data_mutations_helix.txt.gz', sep='\t', low_memory=False)
hc = hc[hc.Variant_Type == 'SNP'].copy()
hc['alt'] = hc.Tumor_Seq_Allele2.str.upper()
hc['key'] = list(zip(hc.Start_Position.astype(int), hc.alt))
hc = hc[hc.key.isin(high.keys())].copy()                 # MT-CYB missense sites only
hc['high'] = hc.key.map(high)
hc['state'] = np.where(hc.HOM_COUNT.fillna(0) > 0, 'homoplasmic in >=1 person', 'heteroplasmic only')

out = hc.groupby(['state', 'high']).size().reset_index(name='n')
out.to_csv(OUTDIR + '/fig4_helix_state.tsv', sep='\t', index=False)
print(out.to_string(index=False))
```
