## Supplementary code for "Relaxed evolutionary constraints on mitochondrial electron-transfer-associated variants in human cancers": fig4_missense_table.py

```
import os as _os, sys as _sys
_sys.path.insert(0, _os.path.dirname(_os.path.abspath(__file__)))
from config import DATADIR, DATAROOT, EXTDIR, STRUCTDIR, ASSETDIR, OUTDIR
import sys; sys.path.insert(0, _os.path.dirname(_os.path.abspath(__file__)))
import numpy as np, pandas as pd
from scipy import stats
from statsmodels.stats.contingency_tables import StratifiedTable
import et_variants as ev
R = DATADIR
V = pd.read_csv(OUTDIR + '/fig3_variants.tsv', sep='\t').set_index(['pos', 'alt'])
hc = pd.read_csv(R + 'data_mutations_helix.txt.gz', sep='\t', low_memory=False); hc = hc[hc.Variant_Type == 'SNP']
poly = hc.groupby(['Start_Position', 'Tumor_Seq_Allele2']).TOTAL_COUNT.sum().to_dict()
def load(p, som, ctcol):
    d = pd.read_csv(R + p, sep='\t', low_memory=False)
    if som: d = d[d['class'] == 'somatic']
    d = d.rename(columns={ctcol: 'ct'})
    d['cohort'] = 'TCGA' if som else 'PCAWG'
    return d
T = load('data_mutations_tcga.txt.gz', True, 'CANCER_TYPE'); P = load('data_mutations_pcawg.txt.gz', False, 'project')
P['ct'] = P.ct.str.split('-').str[0]
D = pd.concat([T, P], ignore_index=True)
D['poly'] = [poly.get((a, b), 0) for a, b in zip(D.Start_Position, D.Tumor_Seq_Allele2)]
snv = D[(D.Variant_Type == 'SNP') & (D.poly <= 10)].copy()
cyb = snv[(snv.Hugo_Symbol == 'MT-CYB') & (snv.Variant_Classification == 'Missense_Mutation')].copy()
cyb['score'] = [V.network.get((p, a), np.nan) for p, a in zip(cyb.Start_Position, cyb.Tumor_Seq_Allele2)]
cyb['high'] = cyb.score >= 0.3
syn = snv[(snv.Variant_Classification == 'Silent')]
print('== 1. Heteroplasmy ==')
for c in ['TCGA', 'PCAWG']:
    h, l, s = [x.loc[x.cohort == c, 'TumorVAF'] for x in (cyb[cyb.high], cyb[~cyb.high], syn)]
    print(f'{c}: high-ET median VAF {h.median():.3f} (n={len(h)}) | low-ET {l.median():.3f} (n={len(l)}) | synonymous {s.median():.3f} (n={len(s)}) '
          f'| high vs low MWU P={stats.mannwhitneyu(h, l).pvalue:.3g} | high vs syn P={stats.mannwhitneyu(h, s).pvalue:.3g}')
    print(f'   fraction VAF>=0.5: high {(h>=0.5).mean():.2f} low {(l>=0.5).mean():.2f} syn {(s>=0.5).mean():.2f}')
print('\n== 2. Cancer types (pooled; share of MT-CYB missense that are high-ET; overall', round(cyb.high.mean(), 3), ') ==')
g = cyb.groupby('ct').agg(n=('high', 'size'), high=('high', 'sum'))
g = g[g.n >= 8].sort_values('n', ascending=False)
p0 = cyb.high.mean()
g['frac'] = g.high / g.n; g['p'] = [stats.binomtest(int(h), int(n), p0).pvalue for h, n in zip(g.high, g.n)]
print(g.round(3).to_string())
print('\n== 3. Co-occurrence with Complex I truncating mutations ==')
ci = D[D.Hugo_Symbol.str.startswith('MT-ND') & D.Variant_Classification.isin(['Nonsense_Mutation', 'Frame_Shift_Del', 'Frame_Shift_Ins'])]
for c in ['TCGA', 'PCAWG']:
    samples = pd.Series(pd.concat([T, P]).query('cohort == @c').drop_duplicates('Tumor_Sample_Barcode').set_index('Tumor_Sample_Barcode').ct)
    allS = set(D.loc[D.cohort == c, 'Tumor_Sample_Barcode'])  # tumours with >=1 mtDNA call
    hiS = set(cyb.loc[(cyb.cohort == c) & cyb.high, 'Tumor_Sample_Barcode']); ciS = set(ci.loc[ci.cohort == c, 'Tumor_Sample_Barcode'])
    tabs = []
    for ct_, ss in samples[samples.index.isin(allS)].groupby(samples): 
        s = set(ss.index)
        a = len(s & hiS & ciS); b = len((s & hiS) - ciS); cc = len((s & ciS) - hiS); d = len(s - hiS - ciS)
        if (a + b) and (a + cc): tabs.append(np.array([[a, b], [cc, d]]) + 0.0)
    st = StratifiedTable(tabs) if tabs else None
    print(f'{c}: tumours with mtDNA calls {len(allS)}, high-ET CYB {len(hiS)}, CI-truncating {len(ciS)}, both {len(hiS & ciS)}; '
          + (f'CMH OR {st.oddsratio_pooled:.2f}, P={st.test_null_odds().pvalue:.3g}' if st else 'n/a'))
cyb.to_csv(OUTDIR + '/fig4_cyb_missense.tsv', sep='\t', index=False)
```
