## Supplementary code for "Relaxed evolutionary constraints on mitochondrial electron-transfer-associated variants in human cancers": fig4_plot.py

```
import os as _os, sys as _sys
_sys.path.insert(0, _os.path.dirname(_os.path.abspath(__file__)))
from config import DATADIR, DATAROOT, EXTDIR, STRUCTDIR, ASSETDIR, OUTDIR
import numpy as np, pandas as pd
import matplotlib; matplotlib.use('Agg')
import matplotlib.pyplot as plt, matplotlib.ticker
from matplotlib.lines import Line2D
from matplotlib.patches import FancyArrowPatch
from scipy import stats

plt.rcParams.update({'font.family': 'DejaVu Sans', 'font.size': 7.5, 'axes.linewidth': 0.6, 'axes.edgecolor': '#000000',
                     'xtick.color': '#000000', 'ytick.color': '#000000', 'text.color': '#000000'})
INK, INK2, MUTED, GRID = '#000000', '#000000', '#9a9893', '#e4e3df'
TUM, GER = '#eb6834', '#2a78d6'
VO = pd.read_csv(OUTDIR + '/fig4_vaf_oe.tsv', sep='\t')
cyb = pd.read_csv(OUTDIR + '/fig4_cyb_missense.tsv', sep='\t')
HS = pd.read_csv(OUTDIR + '/fig4_helix_state.tsv', sep='\t', header=0, names=['state', 'high', 'n'])

fig = plt.figure(figsize=(7.2, 5.6), dpi=300)
gs = fig.add_gridspec(2, 3, width_ratios=[1.25, 0.95, 0.9], height_ratios=[1, 1.05], hspace=0.6, wspace=0.55,
                      left=0.08, right=0.97, top=0.93, bottom=0.09)
def lab(ax, s, x=-0.2): ax.text(x, 1.06, s, transform=ax.transAxes, fontsize=11, fontweight='bold', ha='right', va='bottom')
def clean(ax):
    for s in ('top', 'right'): ax.spines[s].set_visible(False)
    ax.tick_params(length=2)
def title(ax, t): ax.set_title(t, loc='left', fontsize=8, color=INK2)

# a  VAF-stratified O/E
ax = fig.add_subplot(gs[0, :2]); lab(ax, 'a', -0.08)
bins = ['<0.1', '0.1–0.5', '≥0.5']
for k, (c, mk) in enumerate([('TCGA', 's'), ('PCAWG', '^')]):
    d = VO[(VO.cohort == c) & (VO.vaf != 'trend')].set_index('vaf').loc[bins]
    x = np.arange(3) + (k - 0.5) * 0.22
    ax.plot(x, d.oe, color=TUM, lw=1.0, ls='-' if k == 0 else ':')
    for xi, r in zip(x, d.itertuples()):
        ax.plot([xi, xi], [r.lo, r.hi], color=TUM, lw=0.9)
        ax.text(xi, r.hi * 1.05, f'n={r.n}', ha='center', va='bottom', fontsize=5.8, color=INK2)
    ax.scatter(x, d.oe, marker=mk, s=34, color=TUM, edgecolors='white', lw=0.6, zorder=3, label=c)
ax.axhline(1, color=MUTED, lw=0.6, ls='--')
ax.axvspan(1.5, 2.5, color=GRID, alpha=0.6, lw=0)
ax.text(2.0, 0.42, 'majority of mtDNA\ncopies mutant', ha='center', fontsize=6.3, color=INK2)
ax.set_xticks(range(3)); ax.set_xticklabels(bins); ax.set_xlim(-0.5, 2.5)
ax.set_xlabel('tumour heteroplasmy (variant allele fraction)')
ax.set_yscale('log'); ax.set_ylim(0.35, 4.5); ax.set_yticks([0.5, 1, 2, 4]); ax.set_yticklabels(['0.5', '1', '2', '4'])
ax.yaxis.set_minor_formatter(matplotlib.ticker.NullFormatter())
ax.set_ylabel('ET-perturbing missense, O/E'); clean(ax)
ax.legend(frameon=False, fontsize=6.6, loc='upper right', ncol=2, handletextpad=0.2)
title(ax, 'Enrichment is confined to sub-threshold heteroplasmy')

# b  germline heteroplasmic vs homoplasmic
ax = fig.add_subplot(gs[0, 2]); lab(ax, 'b', -0.32)
tab = HS.pivot_table(index='state', columns='high', values='n')
order = ['heteroplasmic only', 'homoplasmic in >=1 person']
fr = []
for i, s in enumerate(order):
    k, n = tab.loc[s, True], tab.loc[s].sum(); p = k / n
    lo, hi = stats.binomtest(int(k), int(n)).proportion_ci()
    ax.bar(i, p * 100, width=0.55, color=GER if i == 0 else '#9cc0ea', edgecolor='none')
    ax.plot([i, i], [lo * 100, hi * 100], color=INK2, lw=0.9)
    ax.text(i, hi * 100 + 0.4, f'{int(k)}/{int(n)}', ha='center', va='bottom', fontsize=6.2, color=INK)
mat = np.array([[int(tab.loc[s_, True]), int(tab.loc[s_, False])] for s_ in order]); orr, pv = stats.fisher_exact(mat)
ax.text(0.5, 16.6, f'OR = {orr:.2f}, P = {pv:.2f}', ha='center', fontsize=6.3, color=INK2)
ax.set_xticks([0, 1]); ax.set_xticklabels(['heteroplasmic\nonly', 'homoplasmic\n(≥1 carrier)'], fontsize=6.0)
ax.set_ylim(0, 18); ax.set_ylabel('ET-perturbing (% of missense sites)'); ax.set_xlim(-0.6, 1.6); clean(ax)
title(ax, 'Germline (HelixMTdb)')

# c  cancer types
ax = fig.add_subplot(gs[1, :2]); lab(ax, 'c', -0.08)
g = cyb.groupby('ct').agg(n=('high', 'size'), k=('high', 'sum')); g = g[g.n >= 8].copy()
p0 = cyb.high.mean()
g['f'] = g.k / g.n
ci = [stats.binomtest(int(k), int(n)).proportion_ci() for k, n in zip(g.k, g.n)]
g['lo'] = [c[0] for c in ci]; g['hi'] = [c[1] for c in ci]
g = g.sort_values('f')
chi = stats.chi2_contingency(np.c_[g.k, g.n - g.k])
for i, r in enumerate(g.itertuples()):
    ax.plot([i, i], [r.lo * 100, r.hi * 100], color=MUTED, lw=0.8)
    ax.scatter(i, r.f * 100, s=10 + r.n * 0.6, color=TUM, edgecolors='white', lw=0.5, zorder=3)
ax.axhline(p0 * 100, color=INK2, lw=0.7, ls='--'); ax.text(len(g) - 0.4, p0 * 100 + 1.5, f'pan-cancer {p0*100:.0f}%', ha='right', fontsize=6.2, color=INK2)
ax.set_xticks(range(len(g))); ax.set_xticklabels(g.index, rotation=90, fontsize=6)
ax.set_ylabel('ET-perturbing (% of MT-CYB missense)'); ax.set_ylim(0, 100); clean(ax)
ax.text(0.01, 0.97, f'heterogeneity χ² = {chi.statistic:.1f}, d.f. = {chi.dof}, P = {chi.pvalue:.2f}', transform=ax.transAxes, va='top', fontsize=6.3, color=INK2)
title(ax, 'A pan-cancer pattern (tumour types with ≥8 mutations; point area ∝ n)')

# d  model
ax = fig.add_subplot(gs[1, 2]); lab(ax, 'd', -0.32)
h = np.linspace(0, 1, 200)
thr = 0.6
fit = np.where(h < thr, 1 + 0.15 * h, 1 + 0.15 * thr - 4.0 * np.clip(h - thr, 0, None) ** 1.4)
ax.plot(h, fit, color=TUM, lw=1.4)
ax.plot(h, np.ones_like(h), color=MUTED, lw=0.8, ls='--')
ax.axvspan(thr, 1, color=GRID, alpha=0.6, lw=0)
ax.text(thr / 2 + 0.03, 1.25, 'tolerated:\nET perturbation\ndiluted by WT copies', ha='center', va='bottom', fontsize=6.0, color=INK)
ax.text((thr + 1) / 2, 1.2, 'threshold\nexceeded:\npurged', ha='center', va='bottom', fontsize=6.0, color=INK)
ax.set_xlim(0, 1); ax.set_ylim(-0.2, 1.75); ax.set_yticks([])
ax.set_xlabel('heteroplasmy'); ax.set_ylabel('relative clonal fitness'); clean(ax); ax.spines['left'].set_visible(True)
title(ax, 'Model')

fig.savefig(OUTDIR + '/Figure4.png'); fig.savefig(OUTDIR + '/Figure4.pdf')
print('heterogeneity', chi.statistic, chi.dof, chi.pvalue)
```
