## Supplementary code for "Relaxed evolutionary constraints on mitochondrial electron-transfer-associated variants in human cancers": fig4_purity.py

```
"""Purity check for Figure 4a (TCGA). Purity: consensus purity estimate (CPE) from Aran et al. 2015
(TCGAbiolinks::Tumor.purity); ABSOLUTE as alternative. Cancer-cell heteroplasmy ~ VAF / purity (capped at 1),
assuming similar mtDNA copy number in tumour and admixed normal cells."""
import os as _os, sys as _sys
_sys.path.insert(0, _os.path.dirname(_os.path.abspath(__file__)))
from config import DATADIR, DATAROOT, EXTDIR, STRUCTDIR, ASSETDIR, OUTDIR

import sys; sys.path.insert(0, _os.path.dirname(_os.path.abspath(__file__)))
import numpy as np, pandas as pd
from scipy import stats
import et_variants as ev
pu = pd.read_csv(ASSETDIR + '/tcga_purity_aran2015.tsv', sep='\t')
for c in ['ESTIMATE', 'ABSOLUTE', 'LUMP', 'IHC', 'CPE']: pu[c] = pd.to_numeric(pu[c].astype(str).str.replace(',', '.'), errors='coerce')
pu = pu.drop_duplicates('Sample.ID').set_index('Sample.ID')
cyb = pd.read_csv(OUTDIR + '/fig4_cyb_missense.tsv', sep='\t')
t = cyb[(cyb.cohort == 'TCGA') & cyb.score.notna()].copy()
t['sid'] = t.Tumor_Sample_Barcode.str[:16]
t['CPE'] = t.sid.map(pu.CPE); t['ABS'] = t.sid.map(pu.ABSOLUTE)
print(f'TCGA MT-CYB missense with score: {len(t)}; with CPE purity: {t.CPE.notna().sum()}; with ABSOLUTE: {t.ABS.notna().sum()}')
F = ev.features('CIII bL->bH'); V = pd.read_csv(OUTDIR + '/fig3_variants.tsv', sep='\t')
rate = ev.rates(ev.cohorts()['TCGA']); w = np.array([rate[(a, b)] for a, b in zip(F.tri, F.alt)]) * ev.poly_ok(F)
f = w[(V.network >= 0.3).values].sum() / w.sum()
out = []
def strat(d, col, label):
    for lo, hi, lab in [(0, 0.1, '<0.1'), (0.1, 0.5, '0.1–0.5'), (0.5, 9, '≥0.5')]:
        s = d[(d[col] >= lo) & (d[col] < hi)]; n = len(s); k = int(s.high.sum())
        if n == 0: continue
        bt = stats.binomtest(k, n, f); ci = bt.proportion_ci()
        out.append(dict(analysis=label, bin=lab, n=n, high=k, oe=k / (n * f), lo=ci[0] / f, hi=ci[1] / f, p=bt.pvalue))
d = t[t.CPE.notna()].copy(); d['h_cpe'] = (d.TumorVAF / d.CPE).clip(upper=1)
strat(d, 'TumorVAF', 'raw VAF (tumours with purity)')
strat(d, 'h_cpe', 'purity-adjusted (CPE)')
da = t[t.ABS.notna()].copy(); da['h_abs'] = (da.TumorVAF / da.ABS).clip(upper=1); strat(da, 'h_abs', 'purity-adjusted (ABSOLUTE)')
strat(d[d.CPE >= 0.7], 'TumorVAF', 'raw VAF, high-purity tumours (CPE >= 0.7)')
r = pd.DataFrame(out); r.to_csv(OUTDIR + '/fig4_purity_oe.tsv', sep='\t', index=False)
print(r.round(3).to_string(index=False))
# does purity differ between high-ET and other missense?
print('median CPE high-ET', d[d.high].CPE.median().round(3), 'other', d[~d.high].CPE.median().round(3), 'MWU P', round(stats.mannwhitneyu(d[d.high].CPE, d[~d.high].CPE).pvalue, 3))
```
