## Supplementary code for "Relaxed evolutionary constraints on mitochondrial electron-transfer-associated variants in human cancers": fig4_tables.py

```
"""Heteroplasmy-stratified O/E of high-ET-perturbation MT-CYB missense (score >= 0.3).
Within each cohort the spectrum-expected fraction f of high-ET among all missense is constant;
O/E in a VAF stratum = observed high / (n missense in stratum * f)."""
import os as _os, sys as _sys
_sys.path.insert(0, _os.path.dirname(_os.path.abspath(__file__)))
from config import DATADIR, DATAROOT, EXTDIR, STRUCTDIR, ASSETDIR, OUTDIR

import sys; sys.path.insert(0, _os.path.dirname(_os.path.abspath(__file__)))
import numpy as np, pandas as pd
from scipy import stats
import et_variants as ev
cyb = pd.read_csv(OUTDIR + '/fig4_cyb_missense.tsv', sep='\t')
F = ev.features('CIII bL->bH'); V = pd.read_csv(OUTDIR + '/fig3_variants.tsv', sep='\t')
C = ev.cohorts(); out = []
for c in ['TCGA', 'PCAWG']:
    rate = ev.rates(C[c]); w = np.array([rate[(t, a)] for t, a in zip(F.tri, F.alt)]) * ev.poly_ok(F)
    f = w[(V.network >= 0.3).values].sum() / w.sum()
    d = cyb[(cyb.cohort == c) & cyb.score.notna()]
    for lo, hi, lab in [(0, 0.1, '<0.1'), (0.1, 0.5, '0.1–0.5'), (0.5, 1.01, '≥0.5')]:
        s = d[(d.TumorVAF >= lo) & (d.TumorVAF < hi)]; n = len(s); k = int(s.high.sum())
        bt = stats.binomtest(k, n, f); ci = bt.proportion_ci()
        out.append(dict(cohort=c, vaf=lab, n=n, high=k, exp_frac=f, oe=k / (n * f), lo=ci[0] / f, hi=ci[1] / f, p=bt.pvalue))
    # trend: logistic of high ~ VAF
    import statsmodels.api as sm
    m = sm.Logit(d.high.astype(int), sm.add_constant(d.TumorVAF)).fit(disp=0)
    out.append(dict(cohort=c, vaf='trend', n=len(d), oe=np.exp(m.params['TumorVAF']), p=m.pvalues['TumorVAF']))
r = pd.DataFrame(out); r.to_csv(OUTDIR + '/fig4_vaf_oe.tsv', sep='\t', index=False)
print(r.round(3).to_string(index=False))
```
