## Supplementary code for "Relaxed evolutionary constraints on mitochondrial electron-transfer-associated variants in human cancers": functional_evidence.py

```
"""Functional-evidence analyses for high-ET-perturbation MT-CYB missense variants (TCGA).
(1) Heteroplasmy (VAF) distribution: do high-ET missense behave like truncating (loss-of-function)
    rather than like low-ET missense or synonymous (neutral)?
(2) Complex I truncating co-occurrence: are tumours carrying high-ET MT-CYB missense enriched for
    truncating mutations in Complex I (ND) genes?
(3) Overlap with MITOMAP-annotated pathogenic MT-CYB variants.
"""
import os as _os, sys as _sys
_sys.path.insert(0, _os.path.dirname(_os.path.abspath(__file__)))
from config import DATADIR, DATAROOT, EXTDIR, STRUCTDIR, ASSETDIR, OUTDIR

import sys
sys.path.insert(0, _os.path.dirname(_os.path.abspath(__file__)))
import numpy as np, pandas as pd
from scipy import stats

R = DATADIR
t = pd.read_csv(R + 'data_mutations_tcga.txt.gz', sep='\t', low_memory=False)
t['PAT'] = t.PATIENT_ID.astype(str)
CYB = (14747, 15887)
ND = {'MT-ND1', 'MT-ND2', 'MT-ND3', 'MT-ND4', 'MT-ND4L', 'MT-ND5', 'MT-ND6'}

# --- CYB missense with ET score ---
miss = pd.read_csv(OUTDIR + '/fig4_cyb_missense.tsv', sep='\t')
miss = miss[miss['class'] == 'somatic'].copy()
miss['grp'] = np.where(miss['high'], 'high-ET missense', 'low-ET missense')

# CYB truncating and synonymous from raw
cyb = t[t.Hugo_Symbol == 'MT-CYB'].copy()
trunc = cyb[cyb.truncating == True].copy()
syn = cyb[cyb.Variant_Classification == 'Silent'].copy()

print('=== (1) Heteroplasmy (VAF) by functional group, TCGA MT-CYB ===')
groups = {
    'high-ET missense': miss[miss.high].TumorVAF.dropna().values,
    'low-ET missense': miss[~miss.high].TumorVAF.dropna().values,
    'synonymous': syn.TumorVAF.dropna().values,
    'truncating (LoF)': trunc.TumorVAF.dropna().values,
}
for g, v in groups.items():
    print(f'  {g:20s} n={len(v):3d}  median VAF={np.median(v):.3f}  frac<0.5={np.mean(v<0.5):.2f}')
# key contrasts (Mann-Whitney, two-sided)
def mw(a, b):
    return stats.mannwhitneyu(groups[a], groups[b], alternative='two-sided').pvalue
print('  high-ET vs synonymous  (VAF): P=%.3g' % mw('high-ET missense', 'synonymous'))
print('  high-ET vs truncating  (VAF): P=%.3g' % mw('high-ET missense', 'truncating (LoF)'))
print('  high-ET vs low-ET      (VAF): P=%.3g' % mw('high-ET missense', 'low-ET missense'))

print('\n=== (2) Complex I (ND) truncating co-occurrence ===')
nd_trunc_pat = set(t[(t.Hugo_Symbol.isin(ND)) & (t.truncating == True)].PAT)
all_pat = set(t.PAT)
hi_pat = set(miss[miss.high].PATIENT_ID.astype(str))
other_pat = all_pat - hi_pat
a = len(hi_pat & nd_trunc_pat); b = len(hi_pat) - a
c = len(other_pat & nd_trunc_pat); d = len(other_pat) - c
OR, p = stats.fisher_exact([[a, b], [c, d]])
print(f'  tumours with high-ET CYB missense: {len(hi_pat)}, carrying ND-truncating: {a} ({a/len(hi_pat):.1%})')
print(f'  other tumours: {len(other_pat)}, carrying ND-truncating: {c} ({c/len(other_pat):.1%})')
print(f'  Fisher OR={OR:.2f}, P={p:.3g}')

print('\n=== (3) MITOMAP pathogenic MT-CYB overlap ===')
mito = cyb[cyb.Pathogenic_Reason == 'MITOMAP']
print(f'  MITOMAP-flagged MT-CYB somatic calls: {len(mito)}')
if len(mito):
    # join to ET score by position
    sc = miss[['Start_Position', 'score', 'high', 'HGVSp_Short']].drop_duplicates('Start_Position')
    mj = mito.merge(sc, on='Start_Position', how='left', suffixes=('', '_m'))
    print(mj[['HGVSp_Short', 'Start_Position', 'TumorVAF', 'score', 'high']].to_string(index=False))

# save source tables
out = pd.DataFrame([dict(group=g, n=len(v), median_vaf=np.median(v), frac_lt_0p5=np.mean(v < 0.5)) for g, v in groups.items()])
out.to_csv(OUTDIR + '/func_vaf_groups.tsv', sep='\t', index=False)
pd.DataFrame([dict(analysis='CplxI_truncating_cooccur', hiET_n=len(hi_pat), hiET_ndtrunc=a,
                   other_n=len(other_pat), other_ndtrunc=c, OR=OR, P=p)]).to_csv(OUTDIR + '/func_cooccur.tsv', sep='\t', index=False)
print('\nsaved func_vaf_groups.tsv, func_cooccur.tsv')
```
