## Supplementary code for "Relaxed evolutionary constraints on mitochondrial electron-transfer-associated variants in human cancers": impact_replication.py

```
"""Independent replication of the two headline endpoints in MSK-IMPACT (deep targeted panel, distinct
platform from TCGA exomes and PCAWG genomes). Same pipeline: strand-aware 192-channel spectrum null,
population-rare site restriction (HelixMTdb carriers <=10), expectation over the restricted universe."""
import os as _os, sys as _sys
_sys.path.insert(0, _os.path.dirname(_os.path.abspath(__file__)))
from config import DATADIR, DATAROOT, EXTDIR, STRUCTDIR, ASSETDIR, OUTDIR

import sys, os
sys.path.insert(0, _os.path.dirname(_os.path.abspath(__file__)))
import numpy as np, pandas as pd
from scipy import stats
import et_variants as ev, fig2_pipeline as fp, qcycle_marcus as qm

R = ev.R
seq = ev.seq

# population polymorphism table (same as ev.cohorts)
hc = pd.read_csv(R + 'data_mutations_helix.txt.gz', sep='\t', low_memory=False); hc = hc[hc.Variant_Type == 'SNP']
poly = hc.groupby(['Start_Position', 'Tumor_Seq_Allele2']).TOTAL_COUNT.sum().to_dict()

im = pd.read_csv(R + 'data_mutations_impact.txt.gz', sep='\t', low_memory=False)
im = im[im['class'].isin(['somatic', 'likely somatic'])]
im = im[im.Variant_Type == 'SNP']
o = pd.DataFrame(dict(sample=im.Tumor_Sample_Barcode.values,
                      pos=im.Start_Position.astype(int).values,
                      alt=im.Tumor_Seq_Allele2.str.upper().values))
o = o[[poly.get((a, b), 0) <= 10 for a, b in zip(o.pos, o.alt)]]
IMP = o
print(f'MSK-IMPACT somatic SNV calls after rare-site filter: {len(IMP)}  (samples: {IMP["sample"].nunique()})')

F = ev.features('CIII bL->bH')
MASK = ev.poly_ok(F)
net, blbh, prox = qm.scores(F, 8.0, 1.0)

# haem charge-change mask
cls = pd.read_csv(ASSETDIR + '/MT-CYB_residue_classes.tsv', sep='\t'); cm = dict(zip(cls.aa_pos, cls.category))
grp = {'Qo site / bL leak path': 'Qo/ISP', 'ISP interface': 'Qo/ISP', 'Qi site': 'Qi', 'heme b environment': 'haem', 'other': 'other'}
F['cls'] = F.aa_pos.map(cm).fillna('other').map(grp)
haem_charge = (F.cls == 'haem').values & (F.dq.abs() >= 0.9).values
high_et = (net >= 0.3)

# count IMPACT MT-CYB missense in region
cybmiss = 0
for r in IMP.itertuples():
    if 14747 <= r.pos <= 15887:
        k, a, b, eff = ev.annotate(r.pos, r.alt, 14747)
        if eff == 'missense': cybmiss += 1
print(f'MSK-IMPACT MT-CYB missense (rare-site): {cybmiss}')

rate = ev.rates(IMP)
W = np.array([rate[(t, a)] for t, a in zip(F.tri, F.alt)]) * MASK
key = IMP.groupby(['pos', 'alt']).size()
Y = np.array([key.get((p, a), 0) for p, a in zip(F.pos, F.alt)])
Ntot = Y.sum()


def oe(mask):
    n = int(Y[mask].sum())
    e = Ntot * W[mask].sum() / W.sum()
    lo = stats.chi2.ppf(0.025, 2 * n) / 2 / e if n else 0.0
    hi = stats.chi2.ppf(0.975, 2 * n + 2) / 2 / e
    p = stats.poisson.cdf(n, e) * 2 if n < e else stats.poisson.sf(n - 1, e) * 2
    return dict(observed=n, expected=round(e, 2), oe=round(n / e, 3), lo=round(lo, 3), hi=round(hi, 3), p=min(p, 1.0))


print('\n=== MSK-IMPACT replication (independent third cohort) ===')
src_rows = []
for lab, m in [('haem-env charge-change missense', haem_charge), ('high-ET missense (score>=0.3)', high_et)]:
    r = oe(m)
    print(f'  {lab:34s} O/E={r["oe"]:.2f}  [{r["lo"]:.2f}-{r["hi"]:.2f}]  obs={r["observed"]} exp={r["expected"]}  P={r["p"]:.2e}')
    src_rows.append(dict(cohort='MSK-IMPACT', endpoint=lab, **r))

# VAF-stratified high-ET O/E (threshold-model check)
IMP['up'] = IMP['alt']
hi_sites = {(int(p), a) for p, a, h in zip(F.pos, F.alt, high_et) if h}
cyb = IMP[(IMP.pos >= 14747) & (IMP.pos <= 15887)].copy()
cyb = cyb[[ev.annotate(int(p), a, 14747)[3] == 'missense' for p, a in zip(cyb.pos, cyb.up)]]
cyb['high'] = [(int(p), a) in hi_sites for p, a in zip(cyb.pos, cyb.up)]
# merge VAF from the raw IMPACT table
imraw = pd.read_csv(R + 'data_mutations_impact.txt.gz', sep='\t', low_memory=False)
vafmap = dict(zip(zip(imraw.Tumor_Sample_Barcode, imraw.Start_Position.astype(int), imraw.Tumor_Seq_Allele2.str.upper()), imraw.TumorVAF))
cyb['vaf'] = [vafmap.get((s, int(p), a), np.nan) for s, p, a in zip(cyb['sample'], cyb.pos, cyb.up)]
expfrac = W[high_et].sum() / W.sum()
for lab, mm in [('high-ET missense, VAF<0.5', cyb.vaf < 0.5), ('high-ET missense, VAF>=0.5', cyb.vaf >= 0.5)]:
    sub = cyb[mm]; n = int(sub.high.sum()); N = len(sub); e = N * expfrac
    p = stats.poisson.sf(n - 1, e) * 2 if n > e else stats.poisson.cdf(n, e) * 2
    print(f'  {lab:28s} n={N} high={n} exp={e:.1f} O/E={n/e:.2f} P={min(p,1):.3g}')
    src_rows.append(dict(cohort='MSK-IMPACT', endpoint=lab, observed=n, expected=round(e, 2),
                         oe=round(n / e, 3), lo=None, hi=None, p=round(min(p, 1), 4)))

pd.DataFrame(src_rows).rename(columns={'p': 'P'}).to_csv(OUTDIR + '/impact_sourcedata.tsv', sep='\t', index=False)
pd.DataFrame(src_rows).to_csv(OUTDIR + '/impact_repl.tsv', sep='\t', index=False)
```
