## Supplementary code for "Relaxed evolutionary constraints on mitochondrial electron-transfer-associated variants in human cancers": multiple_testing_fdr.py

```
"""Benjamini-Hochberg FDR across the primary confirmatory test family.

The haem-environment charge-change and high-ET endpoints are recomputed from raw here (self-contained);
the per-class relaxed-constraint odds-ratio P values are read from fig2_tables output and the
heteroplasmy-threshold P from fig4_tables output. Run fig2_tables.py and fig4_tables.py first.
Writes fdr_primary.tsv.
"""
import os as _os, sys as _sys
_sys.path.insert(0, _os.path.dirname(_os.path.abspath(__file__)))
from config import OUTDIR

import numpy as np, pandas as pd
from scipy import stats
from statsmodels.stats.multitest import multipletests
import et_variants as ev, qcycle_marcus as qm

F = ev.features('CIII bL->bH')
C = ev.cohorts()
MASK = ev.poly_ok(F)
net, _, _ = qm.scores(F, 8.0, 1.0)

cls = pd.read_csv(ev.ASSETDIR + '/MT-CYB_residue_classes.tsv', sep='\t') if hasattr(ev, 'ASSETDIR') \
    else pd.read_csv(OUTDIR + '/../inputs/assets/MT-CYB_residue_classes.tsv', sep='\t')
cm = dict(zip(cls.aa_pos, cls.category))
grp = {'Qo site / bL leak path': 'Qo/ISP', 'ISP interface': 'Qo/ISP', 'Qi site': 'Qi',
       'heme b environment': 'haem', 'other': 'other'}
F['cls'] = F.aa_pos.map(cm).fillna('other').map(grp)
haem_charge = (F.cls == 'haem').values & (F.dq.abs() >= 0.9).values
high_et = (net >= 0.3)

RT = {k: ev.rates(v) for k, v in C.items()}
W = {k: np.array([RT[k][(t, a)] for t, a in zip(F.tri, F.alt)]) * (1 if k == 'germline' else MASK) for k in C}
Y = {}
for k, d in C.items():
    key = d.groupby(['pos', 'alt']).size()
    y = np.array([key.get((p, a), 0) for p, a in zip(F.pos, F.alt)])
    Y[k] = (y > 0).astype(int) if k == 'germline' else y


def pois_p(mask, cohorts):
    n = sum(int(Y[c][mask].sum()) for c in cohorts)
    e = sum(Y[c].sum() * W[c][mask].sum() / W[c].sum() for c in cohorts)
    return (stats.poisson.cdf(n, e) * 2 if n < e else stats.poisson.sf(n - 1, e) * 2), n, e


def binom_p(mask, cohort):
    N = int(Y[cohort].sum()); n = int(Y[cohort][mask].sum())
    f = W[cohort][mask].sum() / W[cohort].sum()
    return stats.binomtest(n, N, f).pvalue


p_haem_g = binom_p(haem_charge, 'germline')
p_haem_t = pois_p(haem_charge, ['TCGA', 'PCAWG'])[0]
p_high_g = pois_p(high_et, ['germline'])[0]
p_high_t = pois_p(high_et, ['TCGA', 'PCAWG'])[0]

# relaxed-constraint Fisher ORs from fig2_tables output
g = pd.read_csv(OUTDIR + '/fig2_out_poly10/fig2_somatic_vs_germline.tsv', sep='\t')
p_haem_or = g.loc[g.category == 'heme b environment', 'p'].iloc[0]
p_qi_or = g.loc[g.category == 'Qi site', 'p'].iloc[0]

# heteroplasmy low-VAF (<0.5) pooled from fig4_tables output
v = pd.read_csv(OUTDIR + '/fig4_vaf_oe.tsv', sep='\t')
low = v[v.vaf.isin(['<0.1', '0.1–0.5'])]
obs = low['high'].sum(); exp = (low['n'] * low['exp_frac']).sum()
p_het = stats.poisson.sf(obs - 1, exp) * 2 if obs > exp else stats.poisson.cdf(obs, exp) * 2

fam = [
    ('Germline depletion: haem-env charge-change missense (binomial)', min(p_haem_g, 1)),
    ('Tumour enrichment: haem-env charge-change missense, pooled (Poisson)', min(p_haem_t, 1)),
    ('Germline depletion: high-ET missense, score>=0.3 (Poisson)', min(p_high_g, 1)),
    ('Tumour enrichment: high-ET missense, pooled (Poisson)', min(p_high_t, 1)),
    ('Relaxed constraint: haem class, somatic vs germline (Fisher)', max(p_haem_or, 1e-5)),
    ('Relaxed constraint: Qi class, somatic vs germline (Fisher)', p_qi_or),
    ('Heteroplasmy threshold: high-ET enrichment at VAF<0.5, pooled (Poisson)', min(p_het, 1)),
]
lab = [x[0] for x in fam]; p = np.array([x[1] for x in fam])
rej, q, _, _ = multipletests(p, method='fdr_bh')
out = pd.DataFrame({'test': lab, 'P_value': p, 'q_BH': q, 'significant_q_lt_0.05': rej})
out.to_csv(OUTDIR + '/fdr_primary.tsv', sep='\t', index=False)
pd.set_option('display.width', 200)
print(out.to_string(index=False))
print('\nAll primary tests significant at q<0.05:', bool(rej.all()))
```
