## Supplementary code for "Relaxed evolutionary constraints on mitochondrial electron-transfer-associated variants in human cancers": nuclear_sbs.py

```
"""Nuclear SBS signature exposures for TCGA tumours (GDC WXS ensemble-masked MAFs).

1. SNVs -> SBS96 matrix using the MAF CONTEXT column (pyrimidine-centred).
2. Fit COSMIC v3.4 GRCh38 *exome* signatures per tumour: NNLS on all non-artefact
   signatures, drop signatures < 3% of mutations, then backward elimination while the
   cosine similarity drops by < 0.01 (SigProfilerAssignment-like decomposition).
3. Oxidative-damage readouts: SBS18 and SBS36 exposures; C>A (G>T) at GG/GGG-type
   guanine runs as a sequence-level proxy for hole-trapping sites.
Output: tcga_sbs_exposures.csv (PCAWG-style wide table), tcga_sbs96.tsv, tcga_sbs_qc.tsv
"""
import os as _os, sys as _sys
_sys.path.insert(0, _os.path.dirname(_os.path.abspath(__file__)))
from config import DATADIR, DATAROOT, EXTDIR, STRUCTDIR, ASSETDIR, OUTDIR

import glob, re, sys
import numpy as np, pandas as pd
from scipy.optimize import nnls

FILES = sorted(glob.glob(STRUCTDIR + '/tcga/TCGA_*_mutations*.csv.gz'))
SIG = pd.read_csv(OUTDIR + '/ext/spa/COSMIC_v3.4_SBS_GRCh38_exome.txt', sep='\t', index_col=0)
ARTEFACT = {'SBS27', 'SBS43', 'SBS95'} | {f'SBS{i}' for i in range(45, 61)}
SIG = SIG[[c for c in SIG.columns if c not in ARTEFACT]]
CH = list(SIG.index)
COMP = str.maketrans('ACGT', 'TGCA')
rc = lambda s: s.translate(COMP)[::-1]


def channel(ctx, ref, alt):
    if len(ctx) < 11 or ctx[5] != ref: return None
    up, dn = ctx[4], ctx[6]
    if ref in 'GA':
        up, ref, alt, dn = rc(dn), rc(ref), rc(alt), rc(up)
    return f'{up}[{ref}>{alt}]{dn}'


def guanine_run_ox(ctx, ref, alt):
    """C>A/G>T where the mutated G sits in a run of >=2 G on its own strand (GG/GGG hole traps)."""
    if (ref, alt) == ('G', 'T'): s = ctx
    elif (ref, alt) == ('C', 'A'): s = rc(ctx)
    else: return 0
    return int(s[4] == 'G' or s[6] == 'G')


def fit(v, S, names):
    x, _ = nnls(S, v)
    tot = v.sum()
    keep = [i for i in range(len(names)) if x[i] >= 0.03 * tot]
    if not keep: keep = [int(np.argmax(x))]
    cos = lambda k: (lambda e: (S[:, k] @ e) @ v / (np.linalg.norm(S[:, k] @ e) * np.linalg.norm(v) + 1e-12))(nnls(S[:, k], v)[0])
    cur = cos(keep)
    while len(keep) > 1:
        trials = [(cos([j for j in keep if j != i]), i) for i in keep]
        best, drop = max(trials)
        if cur - best < 0.01:
            keep.remove(drop); cur = best
        else:
            break
    e, _ = nnls(S[:, keep], v)
    out = np.zeros(len(names)); out[keep] = e
    out *= tot / max(out.sum(), 1e-9)
    return out, cur


rows, ox = {}, {}
meta = {}
for f in FILES:
    ct = re.search(r'TCGA_([A-Z]+)_', f).group(1)
    d = pd.read_csv(f, usecols=['Tumor_Sample_Barcode', 'Variant_Type', 'Reference_Allele', 'Tumor_Seq_Allele2', 'CONTEXT'])
    d = d[(d.Variant_Type == 'SNP')]
    d = d[d.Tumor_Sample_Barcode.str[13:15] < '10']           # tumour samples only
    for r in d.itertuples(index=False):
        ctx = str(r.CONTEXT).upper()
        ch = channel(ctx, r.Reference_Allele, r.Tumor_Seq_Allele2)
        if ch is None: continue
        s = r.Tumor_Sample_Barcode
        rows.setdefault(s, {}); rows[s][ch] = rows[s].get(ch, 0) + 1
        ox[s] = ox.get(s, 0) + guanine_run_ox(ctx, r.Reference_Allele, r.Tumor_Seq_Allele2)
        meta[s] = ct
    print(ct, d.Tumor_Sample_Barcode.nunique(), file=sys.stderr)

M = pd.DataFrame.from_dict(rows, orient='index').reindex(columns=CH).fillna(0).astype(int)
# one tumour per patient: keep the aliquot with most SNVs
M['patient'] = M.index.str[:12]; M['n'] = M[CH].sum(1)
M = M.sort_values('n', ascending=False).drop_duplicates('patient')
pat = M.pop('patient'); M.pop('n')
M.T.to_csv(OUTDIR + '/tcga_sbs96.tsv', sep='\t')

S = SIG.values; names = list(SIG.columns)
E, qc = [], []
for s, v in M.iterrows():
    e, c = fit(v.values.astype(float), S, names)
    E.append(e); qc.append(dict(sample=s, patient=pat[s], cancer_type=meta[s], n_snv=int(v.sum()), cosine=round(c, 4),
                                gg_CtoA=ox[s], gg_CtoA_frac=ox[s] / v.sum() if v.sum() else np.nan))
E = pd.DataFrame(E, index=M.index, columns=names)
out = E.copy(); out.insert(0, 'Sample Names', out.index); out.insert(0, 'Cancer Types', [meta[s] for s in out.index])
out.insert(2, 'Patient', [pat[s] for s in out.index])
out.to_csv(OUTDIR + '/tcga_sbs_exposures.csv', index=False)
qc = pd.DataFrame(qc); qc.to_csv(OUTDIR + '/tcga_sbs_qc.tsv', sep='\t', index=False)
print('tumours', len(E), 'median SNVs', int(qc.n_snv.median()), 'median cosine', qc.cosine.median())
```
