## Supplementary code for "Relaxed evolutionary constraints on mitochondrial electron-transfer-associated variants in human cancers": qcycle_marcus.py

```
"""Multi-step (Q-cycle) Marcus model for MT-CYB missense variants vs a 'charge near cofactor' control.

Redox centres in cytochrome b (9CG3 monomer J): Qo semiquinone (Qo pocket centroid), haem bL Fe,
haem bH Fe, Qi quinone (Qi pocket centroid). ET steps (dG0 / lambda, eV; literature-range values):
  Qo SQ -> bL   -0.20 / 0.8
  bL   -> bH   -0.10 / 0.7
  bH   -> Qi   -0.02 / 0.7
A charge change dq shifts each centre's potential by 14.4*dq/(eps*r) (eV); each step's ddG is the
acceptor-minus-donor differential. Pathways |T_DA| change is included for bL->bH (side-chain knockout).
Scores per variant:
  network = sqrt(sum_i dlog10k_i^2)  (any ET step perturbed)
  bLbH    = |dlog10k| for bL->bH only (previous score)
  prox    = |dEm| at the nearest centre (no differential; control)
Discrimination: same-df comparison of (controls + prox) vs (controls + network) vs both, quasi-Poisson,
spectrum offset, per cohort; plus 'differential-only' variable = network residualised on prox.
"""
import os as _os, sys as _sys
_sys.path.insert(0, _os.path.dirname(_os.path.abspath(__file__)))
from config import DATADIR, DATAROOT, EXTDIR, STRUCTDIR, ASSETDIR, OUTDIR

import sys, json, math
import numpy as np, pandas as pd, gemmi
import statsmodels.api as sm
sys.path.insert(0, _os.path.dirname(_os.path.abspath(__file__)))
import et_variants as ev

kT = 8.617e-5 * 310
STEPS = [('Qo', 'bL', -0.20, 0.8), ('bL', 'bH', -0.10, 0.7), ('bH', 'Qi', -0.02, 0.7)]
st = gemmi.read_structure(STRUCTDIR + '/9CG3.cif'); st.remove_hydrogens(); m = st[0]
pk = json.load(open(ASSETDIR + '/pockets.json'))
CEN = {'Qo': np.mean(pk['Qo']['points'], axis=0), 'Qi': np.mean(pk['Qi']['points'], axis=0),
       'bL': ev.centre(m, ('J', 401, 'FE')), 'bH': ev.centre(m, ('J', 402, 'FE'))}
geo = {}
for r in m['J']:
    if r.het_flag == 'H': continue
    sc = [a for a in r if a.name not in ('N', 'CA', 'C', 'O')] or [a for a in r if a.name == 'CA']
    x = np.mean([[a.pos.x, a.pos.y, a.pos.z] for a in sc], axis=0)
    geo[r.seqid.num] = {k: np.linalg.norm(x - v) for k, v in CEN.items()}


def fc(g, lam): return -((g + lam) ** 2) / (4 * lam * kT) / math.log(10)


def scores(F, eps=8.0, scale_lam=1.0):
    dEm = {c: 14.4 * F.dq / (eps * F.aa_pos.map(lambda k: geo[k][c]).clip(lower=3)) for c in CEN}
    parts = []
    for d, a, g0, lam in STEPS:
        lam *= scale_lam
        ddG = -(dEm[a] - dEm[d])
        dl = fc(g0 + ddG, lam) - fc(g0, lam)
        if (d, a) == ('bL', 'bH'): dl = dl - 2 * F.ko
        parts.append(dl)
    P = np.vstack(parts).T
    net = np.sqrt((P ** 2).sum(1)).clip(max=3)
    prox = np.max(np.abs(np.vstack([dEm[c] for c in CEN]).T), axis=1)
    return net, np.abs(P[:, 1]).clip(max=3), prox


if __name__ == '__main__':
    C = ev.cohorts(); RT = {k: ev.rates(v) for k, v in C.items()}
    F = ev.features('CIII bL->bH')
    out = []
    for eps, sl in [(8, 1.0), (4, 1.0), (16, 1.0), (8, 0.7), (8, 1.4)]:
        net, blbh, prox = scores(F, eps, sl)
        X = pd.DataFrame(dict(network=net, bLbH=blbh, prox=prox, dcharge=F.dq.abs(), dKD=F.dKD, dvol=F.dvol,
                              burial=F.burial, mind=np.minimum(F.rD, F.rA)))
        X = (X - X.mean()) / X.std()
        # differential-only component: network orthogonalised to prox (and dcharge)
        b = np.linalg.lstsq(np.c_[np.ones(len(X)), X.prox, X.dcharge], X.network, rcond=None)[0]
        X['net_resid'] = X.network - np.c_[np.ones(len(X)), X.prox, X.dcharge] @ b
        X['net_resid'] /= X.net_resid.std()
        ctrl = ['dcharge', 'dKD', 'dvol', 'burial', 'mind']
        r_np = np.corrcoef(X.network, X.prox)[0, 1]
        for name, d in C.items():
            key = d.groupby(['pos', 'alt']).size()
            y = np.array([key.get((p, a), 0) for p, a in zip(F.pos, F.alt)])
            if name == 'germline': y = (y > 0).astype(int)
            off = np.log([RT[name][(t, a)] for t, a in zip(F.tri, F.alt)])
            keep = np.ones(len(F), bool) if name == 'germline' else ev.poly_ok(F)   # same site universe as the filtered calls
            def dev(cols):
                return sm.GLM(y[keep], sm.add_constant(X.loc[keep, cols]), family=sm.families.Poisson(), offset=off[keep]).fit(scale='X2')
            rP, rN, rB, rR = dev(ctrl + ['prox']), dev(ctrl + ['network']), dev(ctrl + ['prox', 'network']), dev(ctrl + ['prox', 'net_resid'])
            out.append(dict(eps=eps, lam_scale=sl, cohort=name, n=int(y.sum()), r_network_prox=r_np,
                            dev_prox=rP.deviance, dev_network=rN.deviance, dev_both=rB.deviance,
                            b_net_both=rB.params['network'], p_net_both=rB.pvalues['network'],
                            b_prox_both=rB.params['prox'], p_prox_both=rB.pvalues['prox'],
                            b_diff_only=rR.params['net_resid'], p_diff_only=rR.pvalues['net_resid']))
    res = pd.DataFrame(out); res.to_csv(OUTDIR + '/qcycle_validation.tsv', sep='\t', index=False)
    pd.set_option('display.width', 250); print(res.round(4).to_string(index=False))
```
