## Supplementary code for "Relaxed evolutionary constraints on mitochondrial electron-transfer-associated variants in human cancers": robustness_dedup.py

```
"""Patient-level dedup of TCGA/PCAWG overlap, then recompute the two headline pooled O/E endpoints.
PCAWG uses TCGA patient barcodes directly for the TCGA-US cases; any PCAWG sample whose normalized
patient barcode is already in the TCGA cohort is dropped (kept in TCGA) to avoid double-counting."""
import os as _os, sys as _sys
_sys.path.insert(0, _os.path.dirname(_os.path.abspath(__file__)))
from config import DATADIR, DATAROOT, EXTDIR, STRUCTDIR, ASSETDIR, OUTDIR

import sys, re
sys.path.insert(0, _os.path.dirname(_os.path.abspath(__file__)))
import numpy as np, pandas as pd
from scipy import stats
import et_variants as ev

R = ev.R
def norm(x):
    m = re.match(r'(TCGA-[0-9A-Za-z]+-[0-9A-Za-z]+)', str(x))
    return m.group(1) if m else str(x)

# TCGA patient set
ts = pd.read_csv(R + 'data_samples_tcga.txt.gz', sep='\t', low_memory=False)
tcga_pat = set(norm(x) for x in ts.PATIENT_ID.astype(str))

F = ev.features('CIII bL->bH')
C = ev.cohorts()
MASK = ev.poly_ok(F)

# build deduped PCAWG: drop samples whose normalized barcode is a TCGA patient
pc = C['PCAWG'].copy()
pc['pat'] = pc['sample'].map(norm)
dup_mask = pc['pat'].isin(tcga_pat)
n_dropped_rows = int(dup_mask.sum())
n_dropped_samples = pc.loc[dup_mask, 'sample'].nunique()
C_dedup = dict(C); C_dedup['PCAWG'] = pc[~dup_mask].drop(columns='pat')

print(f'PCAWG rows dropped as TCGA duplicates: {n_dropped_rows} (from {n_dropped_samples} overlapping samples)')
print(f'PCAWG MT-CYB region kept: {len(C_dedup["PCAWG"])} / {len(C["PCAWG"])} total SNV rows')

import qcycle_marcus as qm
net, blbh, prox = qm.scores(F, 8.0, 1.0)

# charge-class mapping for haem endpoint
cls = pd.read_csv(ASSETDIR + '/MT-CYB_residue_classes.tsv', sep='\t'); cm = dict(zip(cls.aa_pos, cls.category))
grp = {'Qo site / bL leak path': 'Qo/ISP', 'ISP interface': 'Qo/ISP', 'Qi site': 'Qi', 'heme b environment': 'haem', 'other': 'other'}
F['cls'] = F.aa_pos.map(cm).fillna('other').map(grp)
haem_charge = (F.cls == 'haem').values & (F.dq.abs() >= 0.9).values
high_et = (net >= 0.3)


def recompute(Cx):
    RT = {k: ev.rates(v) for k, v in Cx.items()}
    W = {k: np.array([RT[k][(t, a)] for t, a in zip(F.tri, F.alt)]) * (1 if k == 'germline' else MASK) for k in Cx}
    Y = {}
    for k, d in Cx.items():
        key = d.groupby(['pos', 'alt']).size()
        y = np.array([key.get((p, a), 0) for p, a in zip(F.pos, F.alt)])
        Y[k] = (y > 0).astype(int) if k == 'germline' else y

    def oe(mask, cohorts):
        n = sum(int(Y[c][mask].sum()) for c in cohorts)
        e = sum(Y[c].sum() * W[c][mask].sum() / W[c].sum() for c in cohorts)
        lo = stats.chi2.ppf(0.025, 2 * n) / 2 / e if n else 0.0
        hi = stats.chi2.ppf(0.975, 2 * n + 2) / 2 / e
        p = stats.poisson.cdf(n, e) * 2 if n < e else stats.poisson.sf(n - 1, e) * 2
        return dict(observed=n, expected=round(e, 2), oe=round(n / e, 3), lo=round(lo, 3), hi=round(hi, 3), p=min(p, 1.0))

    return dict(
        haem_charge_tumour=oe(haem_charge, ['TCGA', 'PCAWG']),
        haem_charge_germline=oe(haem_charge, ['germline']),
        high_et_tumour=oe(high_et, ['TCGA', 'PCAWG']),
        high_et_germline=oe(high_et, ['germline']),
    )


orig = recompute(C)
dedup = recompute(C_dedup)
print('\n=== ORIGINAL (with TCGA/PCAWG overlap) ===')
for k, v in orig.items(): print(f'  {k:24s} O/E={v["oe"]:.2f}  [{v["lo"]:.2f}-{v["hi"]:.2f}]  obs={v["observed"]} exp={v["expected"]}  P={v["p"]:.2e}')
print('\n=== DEDUPLICATED (overlapping patients counted once) ===')
for k, v in dedup.items(): print(f'  {k:24s} O/E={v["oe"]:.2f}  [{v["lo"]:.2f}-{v["hi"]:.2f}]  obs={v["observed"]} exp={v["expected"]}  P={v["p"]:.2e}')

pd.DataFrame({'endpoint': list(orig), 'oe_original': [orig[k]['oe'] for k in orig],
              'oe_dedup': [dedup[k]['oe'] for k in orig],
              'obs_original': [orig[k]['observed'] for k in orig],
              'obs_dedup': [dedup[k]['observed'] for k in orig],
              'p_dedup': [dedup[k]['p'] for k in orig]}).to_csv(OUTDIR + '/dedup_results.tsv', sep='\t', index=False)
print('\nsaved', OUTDIR + '/dedup_results.tsv')
```
