## Supplementary code for "Relaxed evolutionary constraints on mitochondrial electron-transfer-associated variants in human cancers": source_data_extras.py

```
"""Source-data tables for figure panels that had no table of their own.

Fig. 4c  share of somatic MT-CYB missense that are ET-perturbing, per tumour type (>=8 mutations),
         with the chi-squared heterogeneity test reported in the legend.
Supp Fig. 1  the no-polymorphism-filter counterpart of Fig. 2, stacked into one table.

Run after fig4_missense_table.py, and after fig2_tables.py / fig2_charge_split.py have been run a
second time with the filter disabled:
    POLY_MAX=1000000000 python3 fig2_tables.py
    ET_NOFILTER=1 CHARGE_OUT=$MTCYB_OUT/fig2_charge_split_nofilter.tsv python3 fig2_charge_split.py
"""
import os as _os, sys as _sys
_sys.path.insert(0, _os.path.dirname(_os.path.abspath(__file__)))
from config import OUTDIR

import numpy as np, pandas as pd
from scipy import stats

O = OUTDIR + '/'

# --- Fig 4c ---------------------------------------------------------------
# NOTE: do not filter on 'class' here -- the PCAWG rows carry no class column and would be dropped.
m = pd.read_csv(O + 'fig4_cyb_missense.tsv', sep='\t')
p0 = m.high.mean()
g = m.groupby('ct').agg(n=('high', 'size'), high=('high', 'sum')).reset_index()
g = g[g.n >= 8].sort_values('n', ascending=False)
g['pct_high'] = 100 * g.high / g.n
g['pancancer_pct'] = 100 * p0
g['p_vs_pancancer'] = [stats.binomtest(int(k), int(n), p0).pvalue for k, n in zip(g.high, g.n)]
chi2, pchi, dof, _ = stats.chi2_contingency(np.array([g.high, g.n - g.high]))
g['heterogeneity_chi2'] = round(chi2, 3); g['heterogeneity_df'] = dof
g['heterogeneity_P'] = round(pchi, 4)
g.rename(columns={'ct': 'cancer type'}).to_csv(O + 'fig4_cancertype.tsv', sep='\t', index=False)
print('Fig 4c: %d tumour types; pan-cancer share %.3f; chi2=%.2f d.f.=%d P=%.4f'
      % (len(g), p0, chi2, dof, pchi))

# --- Supp Fig 1 -----------------------------------------------------------
parts = []
for fn, lab in [('fig2_out/fig2_obs_exp_real.tsv', 'class O/E'),
                ('fig2_out/fig2_somatic_vs_germline.tsv', 'somatic vs germline OR'),
                ('fig2_charge_split_nofilter.tsv', 'class x charge O/E')]:
    d = pd.read_csv(O + fn, sep='\t'); d.insert(0, 'analysis', lab); parts.append(d)
s1 = pd.concat(parts, ignore_index=True)
s1.to_csv(O + 'suppfig1_nofilter.tsv', sep='\t', index=False)
a = parts[0]
tot = a[a.cohort.str.startswith('Somatic') & (a.category != 'Qo+ISP')].groupby('cohort')['observed'].sum()
print('Supp Fig 1: somatic MT-CYB missense without the filter =', int(tot.sum()), dict(tot))
```
