## Supplementary code for "Relaxed evolutionary constraints on mitochondrial electron-transfer-associated variants in human cancers": supp_figures.py

```
"""Supplementary figures S2 (synonymous calibration), S3 (Complex IV replication),
S4 (nuclear signature fitting QC), S6 (purity-adjusted heteroplasmy)."""
import os as _os, sys as _sys
_sys.path.insert(0, _os.path.dirname(_os.path.abspath(__file__)))
from config import DATADIR, DATAROOT, EXTDIR, STRUCTDIR, ASSETDIR, OUTDIR

import sys; sys.path.insert(0, _os.path.dirname(_os.path.abspath(__file__)))
import numpy as np, pandas as pd
from scipy import stats
import matplotlib; matplotlib.use('Agg')
import matplotlib.pyplot as plt, matplotlib.ticker
from matplotlib.lines import Line2D
import et_variants as ev, fig2_pipeline as fp

plt.rcParams.update({'font.family': 'DejaVu Sans', 'font.size': 7.5, 'axes.linewidth': 0.6, 'axes.edgecolor': '#000000',
                     'xtick.color': '#000000', 'ytick.color': '#000000', 'text.color': '#000000'})
INK, MUTED, GRID = '#000000', '#9a9893', '#e4e3df'
TUM, GER = '#eb6834', '#2a78d6'
COL = dict(fp.COL); COL['other'] = '#a8a7a2'
SH = {'Qo site / bL leak path': 'Qo', 'ISP interface': 'ISP', 'Qi site': 'Qi', 'heme b environment': 'haem', 'other': 'other'}
def clean(ax):
    for s in ('top', 'right'): ax.spines[s].set_visible(False)
    ax.tick_params(length=2)
def lab(ax, s, x=-0.15): ax.text(x, 1.06, s, transform=ax.transAxes, fontsize=11, fontweight='bold', ha='right', va='bottom')
def title(ax, t): ax.set_title(t, loc='left', fontsize=8, color=INK)
def logy(ax, lo, hi, ticks):
    ax.set_yscale('log'); ax.set_ylim(lo, hi); ax.set_yticks(ticks); ax.set_yticklabels([f'{t:g}' for t in ticks])
    ax.yaxis.set_minor_formatter(matplotlib.ticker.NullFormatter())
def pci(n, e):
    return (stats.chi2.ppf(0.025, 2 * n) / 2 / e if n else 0.0), stats.chi2.ppf(0.975, 2 * n + 2) / 2 / e

C = ev.cohorts(); RT = {k: ev.rates(v) for k, v in C.items()}
seq = ev.seq
cls = pd.read_csv(ASSETDIR + '/MT-CYB_residue_classes.tsv', sep='\t'); cm = dict(zip(cls.aa_pos, cls.category))

# ---------------- S2: synonymous calibration ----------------
syn = []
for pos in range(fp.CYB_START, fp.CYB_END - 2):
    t = fp.tri(seq, pos)
    for alt in 'ACGT':
        if alt == t[1]: continue
        k, a, b, eff = fp.annotate_cyb(pos, alt, seq)
        if eff == 'synonymous': syn.append((pos, alt, t, cm.get(k, 'other')))
S = pd.DataFrame(syn, columns=['pos', 'alt', 'tri', 'cls'])
import os as _os
_os.environ['ET_NOFILTER'] = '1'; C2 = ev.cohorts(); RT2 = {k: ev.rates(v) for k, v in C2.items()}; _os.environ.pop('ET_NOFILTER')
rows = []
for name, d in C2.items():
    w = np.array([RT2[name][(t, a)] for t, a in zip(S.tri, S.alt)])   # unfiltered calls: whole site universe
    key = d.groupby(['pos', 'alt']).size()
    y = np.array([key.get((p, a), 0) for p, a in zip(S.pos, S.alt)]); y = (y > 0).astype(int) if name == 'germline' else y
    for c in fp.CLASSES:
        m = (S.cls == c).values; n = int(y[m].sum()); e = y.sum() * w[m].sum() / w.sum(); lo, hi = pci(n, e)
        rows.append(dict(cohort=name, cls=c, observed=n, expected=e, oe=n / e if e else np.nan, lo=lo, hi=hi,
                         p=min(1, 2 * (stats.poisson.cdf(n, e) if n < e else stats.poisson.sf(n - 1, e)))))
S2 = pd.DataFrame(rows); S2.to_csv(OUTDIR + '/supp_S2_synonymous.tsv', sep='\t', index=False)
fig, ax = plt.subplots(figsize=(4.6, 2.8), dpi=300); fig.subplots_adjust(left=0.14, right=0.97, top=0.86, bottom=0.14)
for i, c in enumerate(fp.CLASSES):
    for j, (g, mk, fc) in enumerate([('germline', 'o', 'white'), ('TCGA', 's', None), ('PCAWG', '^', None)]):
        r = S2[(S2.cohort == g) & (S2.cls == c)].iloc[0]; x = i + (j - 1) * 0.24
        ax.plot([x, x], [max(r.lo, 0.06), r.hi], color=COL[c], lw=0.9)
        ax.scatter(x, max(r.oe, 0.06), marker=mk, s=24, facecolor=fc or COL[c], edgecolor=COL[c], lw=1.0, zorder=3)
ax.axhline(1, color=MUTED, lw=0.6, ls='--'); ax.set_xticks(range(5)); ax.set_xticklabels([SH[c] for c in fp.CLASSES])
logy(ax, 0.05, 8, [0.1, 0.25, 0.5, 1, 2, 4]); ax.set_ylabel('synonymous observed / expected'); clean(ax)
ax.legend(handles=[Line2D([], [], marker=m, ls='', mfc=fc or INK, mec=INK, markersize=5, label=l) for l, m, fc in
                   [('germline', 'o', 'white'), ('TCGA', 's', None), ('PCAWG', '^', None)]], frameon=False, fontsize=6.5, ncol=3, loc='upper left')
title(ax, f'Synonymous calibration of the mutational null, all calls (min P = {S2.p.min():.2f})')
fig.savefig(OUTDIR + '/SuppFig_S2_synonymous.png'); fig.savefig(OUTDIR + '/SuppFig_S2_synonymous.pdf'); plt.close(fig)

# ---------------- S3: Complex IV replication ----------------
def step_oe(step, dG0, lam):
    F = ev.features(step); F['score'], _ = ev.marcus(F, dG0, lam, 8.0); MSK = ev.poly_ok(F)
    out = []
    for g, cs in [('germline', ['germline']), ('tumour', ['TCGA', 'PCAWG'])]:
        for lo_, hi_, lb in [(0, 0.1, '<0.1'), (0.1, 0.3, '0.1–0.3'), (0.3, 1, '0.3–1'), (1, 99, '≥1')]:
            m = ((F.score >= lo_) & (F.score < hi_)).values; n = 0; e = 0.0
            for c in cs:
                w = np.array([RT[c][(t, a)] for t, a in zip(F.tri, F.alt)]) * (1 if c == 'germline' else MSK)
                key = C[c].groupby(['pos', 'alt']).size()
                y = np.array([key.get((p, a), 0) for p, a in zip(F.pos, F.alt)]); y = (y > 0).astype(int) if c == 'germline' else y
                n += int(y[m].sum()); e += y.sum() * w[m].sum() / w.sum()
            l, h = pci(n, e); out.append(dict(step=step, group=g, bin=lb, observed=n, expected=e, oe=n / e if e else np.nan, lo=l, hi=h))
    return out
S3 = pd.DataFrame(step_oe('CIII bL->bH', -0.10, 0.70) + step_oe('CIV CuA->heme a', -0.05, 0.40))
S3.to_csv(OUTDIR + '/supp_S3_complexIV.tsv', sep='\t', index=False)
fig, axs = plt.subplots(1, 2, figsize=(6.4, 2.8), dpi=300, sharey=True); fig.subplots_adjust(left=0.1, right=0.97, top=0.84, bottom=0.2, wspace=0.12)
bins = ['<0.1', '0.1–0.3', '0.3–1', '≥1']
for k, (ax, step, ttl) in enumerate(zip(axs, ['CIII bL->bH', 'CIV CuA->heme a'], ['Complex III: haem b$_L$ → b$_H$ (MT-CYB)', 'Complex IV: Cu$_A$ → haem a (MT-CO1/CO2)'])):
    for g, mk, col, fc, ls in [('germline', 'o', GER, 'white', '--'), ('tumour', 's', TUM, None, '-')]:
        d = S3[(S3.step == step) & (S3.group == g)].set_index('bin').reindex(bins)
        x = np.arange(4) + (-0.08 if g == 'germline' else 0.08)
        ax.plot(x, d.oe, color=col, ls=ls, lw=1.1)
        for xi, r in zip(x, d.itertuples()): ax.plot([xi, xi], [max(r.lo, 0.12), r.hi], color=col, lw=0.8)
        ax.scatter(x, d.oe.clip(lower=0.12), marker=mk, s=24, facecolor=fc or col, edgecolor=col, lw=1.0, zorder=3, label=g)
    ax.axhline(1, color=MUTED, lw=0.6, ls='--'); ax.set_xticks(range(4)); ax.set_xticklabels(bins)
    ax.set_xlabel('predicted ET-rate change, |Δlog$_{10}$ k|'); clean(ax); title(ax, ttl); lab(ax, 'ab'[k], -0.08 if k else -0.14)
logy(axs[0], 0.1, 4, [0.25, 0.5, 1, 2]); axs[0].set_ylabel('observed / expected'); axs[1].legend(frameon=False, fontsize=6.5, loc='upper left')
fig.savefig(OUTDIR + '/SuppFig_S3_complexIV.png'); fig.savefig(OUTDIR + '/SuppFig_S3_complexIV.pdf'); plt.close(fig)

# ---------------- S4: nuclear signature fitting QC ----------------
import os as _os3, sys as _sys3
if not _os3.path.exists(OUTDIR + '/tcga_sbs_exposures.csv'):
    print('[skip] Supp Fig S4 (signature QC): tcga_sbs_exposures.csv not found (run nuclear_sbs.py)')
    _sys3.exit(0)
E = pd.read_csv(OUTDIR + '/tcga_sbs_exposures.csv'); Qc = pd.read_csv(OUTDIR + '/tcga_sbs_qc.tsv', sep='\t')
sig = [c for c in E.columns if c.startswith('SBS')]
Fr = E[sig].div(E[sig].sum(axis=1), axis=0); Fr['ct'] = E['Cancer Types']
M = Fr.groupby('ct')[sig].mean()
top = M.max().sort_values(ascending=False).index[:16].tolist()
if 'SBS18' not in top: top[-1] = 'SBS18'
fig = plt.figure(figsize=(7.2, 5.4), dpi=300); g = fig.add_gridspec(2, 1, height_ratios=[1.5, 1], hspace=0.55, left=0.08, right=0.92, top=0.94, bottom=0.08)
ax = fig.add_subplot(g[0]); lab(ax, 'a', -0.04)
im = ax.imshow(M[top].T.values, aspect='auto', cmap=matplotlib.colors.LinearSegmentedColormap.from_list('seq', ['#f7f6f3', '#2a78d6', '#0d3a73']), vmin=0, vmax=0.5)
ax.set_xticks(range(len(M))); ax.set_xticklabels(M.index, rotation=90, fontsize=6); ax.set_yticks(range(len(top))); ax.set_yticklabels(top, fontsize=6)
ax.tick_params(length=0); cb = fig.colorbar(im, ax=ax, fraction=0.025, pad=0.01); cb.set_label('mean fraction of SNVs', fontsize=6.5); cb.ax.tick_params(labelsize=6)
title(ax, f'COSMIC v3.4 exome signature activities, TCGA (n = {len(E):,} tumours)')
ax = fig.add_subplot(g[1]); lab(ax, 'b', -0.04)
order = Qc.groupby('cancer_type').cosine.median().sort_values().index
ax.boxplot([Qc.loc[Qc.cancer_type == c, 'cosine'] for c in order], widths=0.6, showfliers=False, medianprops=dict(color=TUM, lw=1.2),
           boxprops=dict(lw=0.6), whiskerprops=dict(lw=0.6), capprops=dict(lw=0.6))
ax.set_xticks(range(1, len(order) + 1)); ax.set_xticklabels(order, rotation=90, fontsize=6); ax.set_ylabel('reconstruction cosine'); ax.set_ylim(0.4, 1.0); clean(ax)
title(ax, f'Reconstruction accuracy (median cosine {Qc.cosine.median():.2f}; median {int(Qc.n_snv.median())} SNVs per exome)')
fig.savefig(OUTDIR + '/SuppFig_S4_signatureQC.png'); fig.savefig(OUTDIR + '/SuppFig_S4_signatureQC.pdf'); plt.close(fig)

# ---------------- S6: purity ----------------
P6 = pd.read_csv(OUTDIR + '/fig4_purity_oe.tsv', sep='\t')
fig, ax = plt.subplots(figsize=(5.2, 2.9), dpi=300); fig.subplots_adjust(left=0.12, right=0.97, top=0.86, bottom=0.17)
an = P6.analysis.unique(); mk = ['s', 'D', 'v', 'o']; cols = [TUM, '#b5461a', '#7a2d0f', '#f2a57f']
bins = ['<0.1', '0.1–0.5', '≥0.5']
for k, a in enumerate(an):
    d = P6[P6.analysis == a].set_index('bin').reindex(bins); x = np.arange(3) + (k - 1.5) * 0.15
    for xi, r in zip(x, d.itertuples()):
        if np.isnan(r.oe): continue
        ax.plot([xi, xi], [max(r.lo, 0.15), r.hi], color=cols[k], lw=0.9)
    ax.scatter(x, d.oe, marker=mk[k], s=24, color=cols[k], edgecolors='white', lw=0.5, zorder=3, label=a)
ax.axhline(1, color=MUTED, lw=0.6, ls='--'); ax.axvspan(1.5, 2.5, color=GRID, alpha=0.6, lw=0)
ax.set_xticks(range(3)); ax.set_xticklabels(bins); ax.set_xlim(-0.5, 2.5); ax.set_xlabel('heteroplasmy (VAF or purity-adjusted)')
logy(ax, 0.15, 5, [0.25, 0.5, 1, 2, 4]); ax.set_ylabel('ET-perturbing missense, O/E'); clean(ax)
ax.legend(frameon=False, fontsize=5.8, loc='upper right'); title(ax, 'TCGA: robustness of Fig. 4a to tumour purity')
fig.savefig(OUTDIR + '/SuppFig_S6_purity.png'); fig.savefig(OUTDIR + '/SuppFig_S6_purity.pdf'); plt.close(fig)
print(S2.round(2).to_string(index=False)); print(S3.round(2).to_string(index=False))
```
